# Synthetic multicolor antigen-stabilizable nanobody platform for intersectional labelling and functional imaging

**DOI:** 10.1101/2025.10.27.684934

**Authors:** Natalia V. Barykina, Erin M. Carey, Olena S. Oliinyk, Juliana M Mendonça-Gomes, Sofia de Oliveira, Axel Nimmerjahn, Vladislav V. Verkhusha

**Affiliations:** Department of Genetics, and Gruss-Lipper Biophotonics Center, Albert Einstein College of Medicine, Bronx, NY 10461, USA; Waitt Advanced Biophotonics Center, Salk Institute for Biological Studies, La Jolla, CA 92037, USA; Medicum, Faculty of Medicine, University of Helsinki, Helsinki 00290, Finland; Department of Developmental and Molecular Biology, Albert Einstein College of Medicine, Bronx, NY, USA; Department of Developmental and Molecular Biology, Harold and Muriel Block Institute for Clinical and Translational Research, and Cancer Dormancy Institute, Albert Einstein College of Medicine, Bronx, NY, USA

## Abstract

We present a synthetic toolkit of antigen-stabilizable fluorescent nanobodies (VIS-Fbs) spanning the entire visible spectrum from 450 nm to 660 nm. By engineering over twenty fluorescent proteins (FPs) and biosensors into eight nanobodies, we established a generalizable design of VIS-Fbs, which fluoresce brightly only upon binding to cognate antigens. Our synthetic approach includes constitutive, photoactivatable and photoswitchable FPs, and intensiometric FP-based biosensors. VIS-Fbs carrying biosensors enable simultaneous monitoring of two metabolites at confined locations, while FP-based VIS-Fbs targeting biosensors allow ratiometric functional imaging in the mouse brain. We further used VIS-Fbs to track endogenous β-catenin dynamics in zebrafish embryos during normal development and under Wnt/β-catenin signaling modulation. VIS-Fbs provide background-free visualization of intracellular proteins, multicolor detection of multiple antigens, and selective targeting of defined cell populations and compartments. This synthetic biology-driven platform enables precise studies of protein dynamics, cellular processes, and complex biological systems with high specificity and minimal background.

## Introduction

The development of multicolor fluorescent proteins (FPs) with a wide variety of photochemical properties^1-3^ has revolutionized biological sciences, making it possible to visualize gene expression, protein dynamics and interactions in live cells and whole organisms. While direct fusions of proteins with FPs enable imaging of protein tracking, FP tags can interfere with the protein’s function, localization, expression, and turnover^4^. Incorporation of unnatural amino acids for biorthogonal labeling can be an alternative for high resolution imaging of endogenous proteins^5^, however this method can be less efficient than expression of FPs and requires careful optimization. Nanobodies (Nbs)^6^ offer an alternative to tagging endogenous proteins to address these challenges. Nbs derived from camelid antibodies are small (~15 kDa) genetically encoded VHH (variable heavy domain of heavy chain) domains, consisting of three complementarity determining regions (CDRs), which are the hypervariable loops responsible for antigen binding, and four framework regions (FRs), which are conserved sequences that connect and support the CDRs^7^. Nbs bind specific antigens with high affinity and specificity and can be engineered to carry fluorescent tags to visualize intracellular antigens^8, 9^. Recombinantly produced Nbs can be conjugated with various synthetic dyes^10^ and applied for high-resolution labeling of intracellular structures^11^ and transsynaptic neuronal contacts^12^. However, Nbs conjugated with dyes require delivery to cells and may result in high fluorescence background and abnormal accumulation in non-targeted compartments^11^.

Fusions of Nbs with FPs, known as chromobodies, allow protein tracking in live cells^13^. However, genetically encoded chromobodies are produced in mammalian cells regardless of their cognate antigen, leading to an accumulation of unbound fluorescent Nbs, which again increases background fluorescence and hinders visualization of intracellular targets^10, 13^. To reduce the background fluorescence caused by unbound chromobodies, one can express them under weaker promoters^14^ or generate cell lines expressing very low levels of chromobodies^10^. While these strategies help in some applications, they are unsuitable for imaging populations that express target antigens heterogeneously. It has also been proposed to introduce point mutations in the FRs of Nbs to cause chromobodies to degrade when unbound to an antigen^15^; however, these mutations may negatively affect their binding affinity and cellular interactions^16^.

Recently, we engineered antigen-stabilized near-infrared (NIR) fluorescent Nbs, termed NIR-Fbs^8, 9^. They consist of small NIR FPs, based on single GAF domains of the cyanobacteriochrome photoreceptors^17^, inserted into the FRs of various Nbs. When bound to a cognate antigen, NIR-Fbs are highly stabilized and, consequently, fluorescent. However, in the absence of an antigen, they rapidly degrade and do not produce fluorescence. It has been hypothesized that without the bound antigen, the NIR-Fb construct becomes allosterically unstable and targeted by ubiquitinases due to the presence of hydrophobic patches^8, 9^. These patches are thought to appear due to the Nb structure being stretched by the rigid, closely located N- and C-helical termini of the inserted NIR FP. NIR-Fbs were successfully used for background-free visualization of various overexpressed and genomically encoded intracellular proteins. However, NIR-Fbs technology has been limited to the NIR spectral range since all current NIR FPs exhibit emissions above 670 nm^18-21^.

Expanding NIR-Fb approach to GFP-like FPs is a major challenge due to their much larger β-can structure, which is twice as large as the GAF domain, and flexible heterogeneous termini separated by ~2.5 nm. Inserting such bulky and stable FPs into nanobodies risked disrupting antigen binding and antigen-dependent stabilization. In this work, we extended the fluorescence range of these synthetic antigen-dependent fluorescent Nbs to the visible spectrum to allow multicolor protein tagging and multiplexed imaging. To harness the β-can scaffold of GFP-like FPs for this purpose, we engineered linkers to achieve the unique property of antigen-dependent stabilization while preserving the photophysical and biochemical properties of the inserted GFP-like FPs. These consist of constitutively fluorescent FPs (including FPs with large Stokes shift (LSS)^22-24^), dark-to-fluorescent state photoactivatable FPs, FPs photoconvertible from one fluorescent color to another, and FP-based biosensors to various metabolites. By engineering various FPs and biosensors into visible fluorescent antigen-stabilizable Nbs, termed VIS-Fbs, we demonstrated the generalizability of our design. The developed VIS-Fbs were thoroughly characterized individually and validated in mammalian cells. We applied VIS-Fbs in an intersectional approach to selectively target and trace behavior-evoked calcium activity in specific cell types and compartments, including excitatory pyramidal neurons, GABAergic interneurons, and astrocytes, in the somatosensory cortex of transgenic mice. Using VIS-Fb, we captured *in vivo* modulation of endogenous β-catenin levels before gastrulation, following pharmacological activation and inhibition. This provides novel insight into the timing and sensitivity of β-catenin regulation during the early zebrafish embryogenesis.

## Results

### Design of destabilized fluorescent nanobody with inserted mCherry

First, we inserted mCherry^25^ into a nanobody to GFP (Nb_GFP_)^16^ between residues Ser65 and Val66, a position that has been shown to tolerate insertion of miRFP670nano3 in NIR-Fbs^9^, without linkers, with both N- and C-terminal Gly_2_Ser linkers, or 5 deleted amino acid (a.a.) residues from both N- and C-termini of mCherry (**Fig. 1a, Supplementary Figs. 1 and 2**). All variants of the Nb_GFP_ with inserted mCherry exhibited red fluorescence in the cells co-expressing mEGFP as a model antigen, but to various extents (**Supplementary Fig. 1**). Without mEGFP, fluorescence of the insertions was not detected (**Supplementary Fig. 1**), suggesting that it depends on the presence of the antigen. The highest contrast between antigen-bound and destabilized forms was detected for the insertion with mCherry truncated by 5 a.a. from both termini (**Supplementary Fig. 1**). We subjected this variant to further optimization.

**Figure 1.**
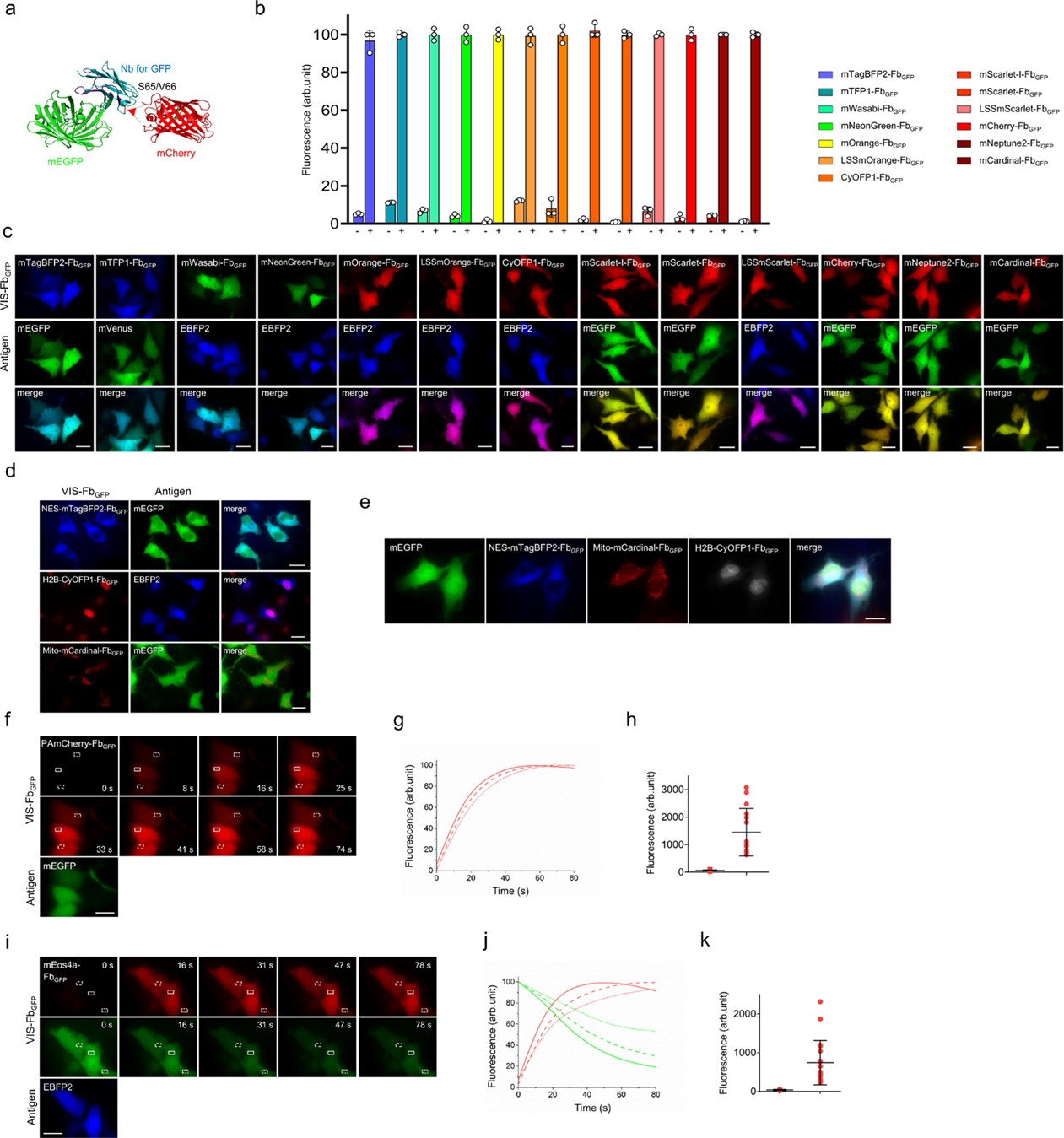
A series of fluorescent nanobodies to GFP (VIS-Fbs_GFP_) spanning the visible spectrum. **(a)** Scheme of a nanobody with inserted red FP (PDB ID: 1ZGO) bound to GFP (PDB ID: 3OGO). Complementarity-determining regions (CDRs) are highlighted in violet. The position of mCherry insertion to the anti-GFP nanobody is indicated with a red arrow. **(b)** Fluorescence intensity of cells transfected with 13 different VIS-Fbs to GFP. The following FPs were co-expressed with VIS-Fbs as positive controls: *mEGFP* for mTagBFP2-Fb_GFP_, mScarlet-I-Fb_GFP_, mScarlet-Fb_GFP_, mCherry-Fb_GFP_, mNeptune2-Fb_GFP_, mCardinal-Fb_GFP_; *mVenus* for mTFP1-Fb_GFP_; *EBFP2* for mWasabi-Fb_GFP_, mNeonGreen-Fb_GFP_, mOrange-Fb_GFP_, LSSmOrange-Fb_GFP_, CyOFP1-Fb_GFP_, LSSmScarlet-Fb_GFP_. The following FPs were co-expressed with VIS-Fbs as negative controls: *mCherry* for mTagBFP2-Fb_GFP_, mTFP1-Fb_GFP_; *mTagBFP2* for other VIS-Fbs. **(c)** Fluorescence images of HeLa cells co-expressing VIS-Fbs from (a) with their cognate antigens. **(d)** HeLa cells co-expressing NES-mTagBFP2-Fb_GFP_ with mEGFP, H2B-CyOFP1-Fb_GFP_ with mEGFP, mito-mCardinal-Fb_GFP_ with mEGFP. **(e)** Multicolor fluorescence images for HeLa cells co-expressing mEGFP antigen with three nanobodies in different compartments: NES-mTagBFP2-Fb_GFP_ in the cytoplasm, mito-Cardinal-Fb_GFP_ in the mitochondria and H2B-CyOFP1-Fb_GFP_ in the nucleus. **(f)** Change in fluorescence intensity of HeLa cells co-expressing PAmCherry-based VIS-Fb_GFP_ and mEGFP in response to 390 nm irradiation. **(g)** Change in fluorescence intensity of representative cells co-expressing PAmCherry-based VIS-Fb_GFP_ and mEGFP in response to 390 nm irradiation. Change of fluorescence for 3 ROIs from (a) is presented. **(h)** Contrast of PAmCherry-based VIS-Fb_GFP_ co-expressed with either mTagBFP2 (*n=18*) or mEGFP (*n=15*) after 390 nm irradiation. **(i)** Change in fluorescence intensity of HeLa cells co-expressing mEos4a-Fb_GFP_ (red and green forms) in response to 390 nm irradiation. **(j)** Change in fluorescence intensity of representative cells co-expressing mEos4a-Fb_GFP_ and EBFP2 in response to 390 nm irradiation. Change of fluorescence for 3 ROIs from (d) is presented. **(k)** Contrast of mEos4a-Fb_GFP_ co-expressed with either mTagBFP2 (*n=14*) or EBFP2 (*n=20*) after 390 nm irradiation. In (b), fluorescence intensity was analyzed by flow cytometry using a 405 nm excitation laser and 450/50 nm emission filter for mTagBFP2, EBFP2, mTagBFP2-Fb_GFP_, 510/40 nm emission filter for mTFP1-Fb_GFP_, 595/30 emission filter for LSSmOrange-Fb_GFP_; a 488 nm excitation laser and 525/50 nm emission filter for mEGFP, mWasabi-Fb_GFP_, mNeonGreen-Fb_GFP_, 537/32 nm for mVenus; 582/15 nm emission filter for CyOFP1-Fb_GFP_, 610/20 nm emission filter for LSSmScarlet-Fb_GFP_; a 561 nm excitation laser and 582/15 nm emission filter for mOrange-Fb_GFP_, mScarlet-Fb_GFP_ and mScarlet-I-Fb_GFP_; 610/20 nm emission filter for mCherry and mCherry-Fb_GFP_; 670/30 nm emission filter for mNeptune2-Fb_GFP_ and mCardinal-Fb_GFP_. The maximal fluorescence of antigen-bound form for each VIS-Fb_GFP_ was assumed to be 100%. Data are presented as mean values ± s.d. for *n* = 3 transfection experiments. In (c – f and i) the following filters were used: for imaging mTagBFP2-Fb_GFP_ and EBFP2 390/40 nm excitation and 460/40 nm emission; for imaging mEGFP, mVenus, mWasabi-Fb_GFP_, mNeonGreen-Fb_GFP_ and green form mEos4a-Fb_GFP_ 480/40 nm excitation and 535/40 nm emission; for imaging mTFP1-Fb_GFP_ 436/20 nm excitation and 480/40 nm emission; for imaging mOrange-Fb_GFP_ 543/22 nm excitation and 580/30 nm emission; for imaging LSSmOrange-Fb_GFP_ 436/20 nm excitation and 575/30 nm emission; for imaging CyOFP1-Fb_GFP_ 523/20 nm excitation and 588/21 nm emission; for imaging LSSmScarlet-Fb_GFP_ 475/42 nm excitation and 615/30 nm emission; for imaging mCherry-Fb_GFP_, mScarlet-I-Fb_GFP_, mScarlet-Fb_GFP_, red form of mEos4a-Fb_GFP_ and PAmCherry-based VIS-Fb_GFP_ 575/25 nm excitation and 615/30 nm emission; for imaging mNeptune2-Fb_GFP_ and mCardinal-Fb_GFP_ 605/30 nm excitation and 667/30 nm emission. Scale bar, 40 μm.

Compared to miRFP670nano3, GFP-like FPs have long, flexible termini that vary in length and a.a. contents depending on their origin. The termini may affect the folding, oligomeric state, and stability of GFP-like FPs^26^. However, it was shown that a minimal fluorescent GFP-like domain tolerates the deletion of up to 8-9 a.a. residues from both termini^27, 28^. To improve the contrast of mCherry insertions into Nb_GFP_ in a stabilized versus destabilized state, we next performed systematic deletion of a.a. residues from N- and C-termini of mCherry (**Supplementary Fig. 3a, b**). The highest contrast was observed with the mCherry variant truncated by 9 a.a. residues from the N-terminus (ΔMVSKGEEDN) (**Supplementary Fig. 3b, c**). We propose that truncating N-terminal residues from the mCherry inserted into Nb_GFP_ reduces its stability, potentially shortens its half-life in antigen-free state and therefore improves background in the absence of cognate antigen. We termed this insertion mCherry-Fb_GFP_ and used it as a framework to engineer other VIS-Nbs for GFP.

To assess whether an antigen concentration affects the protein level of mCherry-Fb_GFP_, we co-transfected cells with the fixed amount of a plasmid encoding mCherry-Fb_GFP_ and decreasing amounts of a plasmid encoding mEGFP antigen. Though the amount of the mCherry-Fb_GFP_ plasmid was the same in all cell transfections, the fluorescence intensity of mCherry-Fb_GFP_ varied and depended on the mEGFP level **(Supplementary Fig. 4 and 5)**, confirming that mCherry-Fb_GFP_ is antigen-dependent and occurs in a dose-dependent manner.

To find how long it takes to detect mCherry-Fb_GFP_ after the start of antigen expression, we co-expressed mCherry-Fb_GFP_ together with mEGFP antigen under the tetracycline-inducible (TRE) promoter along with Tet On plasmid. We then treated cells pre-expressing mCherry-Fb_GFP_ with 1 μg/mL doxycycline (Dox) for 0, 1.5, 4, 18, 24, or 48 h and analyzed the fluorescence by flow cytometry (**Supplementary Fig. 6a, b**). A minimal increase in mCherry-Fb_GFP_ red fluorescence was observed as early as 1.5 h after Dox induction, with a maximal increase detected after 48 h of Dox treatment, which correlated well with a green fluorescence of mEGFP under the Dox-inducible promoter (**Supplementary Fig. 6c and d**).

### Series of constitutively fluorescent nanobodies spanning the visible spectrum

We next analyzed the databases of modern GFP-like FPs and chose 12 bright monomeric constitutive FPs, having emission maxima across the entire visible light spectrum, from blue to far-red: mTagBFP2^29^, mTFP1^30^, mWasabi^31^, mNeonGreen^32^, mOrange^25^, LSSmOrange^33^, CyOFP1^34^, mScarlet-I^35^, mScarlet^35^, LSSmScarlet^36^, mNeptune2^37^, and mCardinal^37^. Since the selected FPs are heterogeneous in terms of their N- and C-termini, we aligned their a.a. sequences with that of the truncated mCherry variant utilized in mCherry-Fb_GFP_ and adjusted the N- and C-termini of FPs accordingly (**Supplementary Fig. 7**). Specifically, we truncated the N-terminal regions of the FPs aligned with the nine N-terminal a.a. residues of mCherry (MVSKGEEDN), resulting in truncations of 3 to 9 residues depending on the FP sequence. Additionally, we incorporated 9th and 10th a.a. residues (MA) from mCherry to the N-termini of the FPs to achieve full alignment (**Supplementary Fig. 7**). Regarding the C-termini, most FPs remained intact, except *Entacmaea quadricolor*-derived FPs (mTagBFP2, mCardinal, mNeptune2) and mEos4a. For these, we deleted C-terminal amino acids to align their C-ends with that of mCherry, ensuring they all terminate with a Lys (**Supplementary Fig. 7**). The resulting nucleotide sequences of FPs were inserted between residues Ser65 and Val66 of Nb_GFP_^16^.

All VIS-Fbs were co-expressed with the cognate antigens in HeLa cells: with mEGFP for the blue, red, and far-red VIS-Fbs and with EBFP2 for the green, orange VIS-Fbs or VIS-Fbs with LSS phenotype (**Fig. 1b, c**). EBFP2 is a blue FP derived from *Aequoria victoria* GFP, and due to the structural similarity between FPs, it can also be recognized by Nb_GFP_. In contrast, mTagBFP2, which is derived from *Entacmaea quadricolor*, is not recognized by Nb_GFP_. Using flow cytometry and fluorescence microscopy, we found that in the absence of the antigen, the cells transfected with VIS-Fbs were not fluorescent. However, when mEGFP or EBFP2 was co-expressed, we observed bright fluorescence at the range specific to the inserted FP, indicating that the engineered VIS-Fb_GFP_s are stabilized (**Fig. 1b, c; Supplementary Fig. 8**). The contrast of various VIS-Fbs in stabilized versus destabilized states ranged ~10-110-fold for different FPs.

To assess how insertion of FP affects brightness of VIS-Fbs for GFP, comparing to the brightness of the parental FPs, we co-transfected mammalian cells with the representative VIS-Fb_GFP_s: mTagBFP2-Fb_GFP_, mOrange-Fb_GFP_, mCherry-Fb_GFP_, mScarlet-Fb_GFP_ or mCardinal-Fb_GFP_ under CMV promoter together with mEGFP or EBFP2 antigen. In the same experiment, mTagBFP2, mOrange, mCherry, mScarlet and mCardinal FPs under CMV promoter were co-transfected with mEGFP or EBFP2 (**Supplementary Figs. 9 and 10**). The flow cytometry data showed that representative VIS-Fb_GFP_s are 35-70% as bright as incorporated FPs in mammalian cells (**Supplementary Figs. 9 and 10**).

Finally, when we fused one of the VIS-Fb_GFP_s, such as mCardinal-Fb_GFP_, to mTagBFP2 via a long flexible linker, its fluorescence was dependent on the presence of mEGFP antigen, similarly to the non-fused mCardinal-Fb_GFP_ fluorescence, confirming that VIS-Fbs are destabilized in the absence of antigen (**Supplementary Fig. 11a, b**). Notably, the fluorescence of the fused mTagBFP2 was also antigen-dependent, suggesting that fusion partners can adopt the same stabilization–degradation behaviors as the VIS-Fb. This highlights the potential to fuse other proteins to VIS-Fbs, enabling antigen-dependent control of their stability.

### Single-antigen-dependent multicolor cell imaging

One of the advantages of multicolor VIS-Fbs is that they can visualize multiple targets in a cell or a single target in different cellular organelles. To visualize an antigen in different cellular compartments simultaneously, we first fused VIS-Fbs with various subcellular localization signals. We fused mTagBFP2-Fb_GFP_ with nuclear exclusion signal (NES) to localize it in the cytoplasm, CyOFP1-Fb_GFP_ with H2B to direct it to the nucleus, and mCardinal-Fb_GFP_ with a mitochondrial localization signal to target it to the mitochondrial matrix. We co-expressed each VIS-Fb fusion in mammalian cells together with mEGFP and observed bright blue, LSS red, or far-red emission in the cytoplasm, nucleus and mitochondria, respectively (**Fig. 1d**). When the fusions were co-expressed with mCherry or mTagBFP2 as negative controls, no fluorescence was observed, confirming the antigen-stabilized property of VIS-Fbs fused with localization signals (**Supplementary Fig. 12**). We next co-expressed all three VIS-Fbs with mEGFP and simultaneously detected their spatially localized respective multicolor fluorescence signals (**Fig. 1e**). We predict that because of high affinity of Nb_GFP_ (Kd < 1nM), the portion of mEGFP antigen may be redirected to certain organelles by VIS-Fb_GFP_s, when VIS-Fb_GFP_ is fused with a specific localization signal. Similarly, it has been shown that a low nanomolar Nb for survivin (SVVNb8) fused with a mitochondria localization signal, peroxisome targeting signal, nuclear localization and nuclear exclusion signals delocalizes endogenous survivin to mitochondria, peroxisomes, nucleus and cytoplasm, respectively^38^.

Using this approach, it is potentially possible to spatially control the activity of GFP-tagged proteins. By targeting VIS-Fb_GFP_s to various organelles and fusing VIS-Fb_GFP_s with functional effectors, such as degradation domains or enzymes, it would be possible to study and manipulate organelle-specific protein functions or specific pathways. This strategy could be especially valuable in GFP-expressing transgenic mouse lines, enabling precise spatial control of protein activity *in vivo*. Another possible application is intersectional labeling of multiple cell populations in GFP-expressing transgenic mice. By targeting VIS-Fb_GFP_s with distinct subcellular localization signals, it could be possible to study the connections between specific cohorts of cells during behavior.

### Photoactivatable and photoswitchable fluorescent nanobodies

Spatiotemporal labeling and tracking of proteins in cells and cells in tissues demand VIS-Fbs with optical highlighters, such as photoactivatable and photoconvertible FPs^3, 39, 40^. To engineer photoactivatable fluorescent Nb, we selected widely used PAmCherry, which photoactivates from a dark-to-red state with violet light^41, 42^. We inserted PAmCherry in Nb_GFP_ between residues Ser65 and Val66 (**Supplementary Fig. 7**) and co-expressed it in HeLa cells with either mEGFP (**Fig. 1f**) or mTagBFP2. Upon irradiation with 390 nm light, the red fluorescence of PAmCherry-based VIS-Fb_GFP_ was activated over 60 s, which is comparable to the activation kinetics of the parental PAmCherry FP (**Fig. 1g; Supplementary Fig. 13a**). Its red fluorescence intensity increased more than 20-fold (**Fig. 1h**). We also compared contrasts between dark and light-activated states for PAmCherry-Fb_GFP_ and its parental PAmCherry, and observed no significant difference (**Supplementary Fig. 13b**).

For designing photoconvertible VIS-Fb, we inserted mEos4a^43^, which photoconverts from green to red fluorescent states by violet light, in the Nb_GFP_ between residues Ser65 and Val66. As the C-terminus of mEos4a differs significantly from that of mCherry, we made a series of modifications to align the mEos4a sequence with the mCherry sequence. Specifically, we truncated three amino acid residues (ΔMVS) and inserted two amino acid residues (MA) at the N-terminus and truncated one residue (ΔR) and inserted two amino acid residues (YK) at the C-terminus (**Supplementary Fig. 7**). The resulting mEos4a-based VIS-Fb_GFP_ exhibited two forms, green and red (**Fig. 1i**). When co-expressed with GFP-derived EBFP2 model antigen, which is also recognized by Nb_GFP_, the mEos4a-Fb_GFP_ construct exhibits green fluorescence, whereas the 60-80 s exposure to 390 nm light caused it to shift its emission to red, demonstrating photoconversion kinetics comparable to those of the parental mEos4a FP (**Fig. 1i, j; Supplementary Fig. 13c**). Likewise, the green forms of mEos4a-Fb_GFP_ and the parental mEos4a FP exhibited comparable photoconversion contrasts (**Supplementary Fig. 13d**). Notably, when co-expressed with mTagBFP2 negative control, no mEos4a-Fb_GFP_ fluorescence was detected (**Supplementary Fig. 14**). The red fluorescence contrast between stabilized (EBFP2-bound) and destabilized conditions was 19.3-fold (**Fig. 1k**). These results demonstrated the preservation of the phenotypes and photophysics of the inserted FPs while demonstrating the versatility of our VIS-Fb engineering approach.

### Expanding the engineering platform to other nanobodies

To study whether our approach can be expanded to Nbs other than Nb_GFP_, we applied it to four other Nbs to different antigens: LAM2 Nb for mCherry^44^, 59H10 Nb specific for p24 HIV protein^45^, 2E7 Nb targeting the gp41 HIV protein^46^, and Nb against ALFA-tag^46^.

We first inserted various FPs, truncated at the N-terminus, into these Nbs. As a result, we observed no or relatively low fluorescence when the engineered VIS-Fbs were co-expressed with their cognate antigens. The most promising results were obtained for blue-green FPs (mTagBFP2, mWasabi, mClover3^47^) inserted into the LAM2 Nb against mCherry (VIS-Fbs_LAM2_) (**Supplementary Figs. 15a, 16a-c, 17**).

Since the contrasts between antigen-bound and antigen-free states of VIS-Fbs_LAM2_ ranged from 2 to 3 (**Supplementary Figs. 16b and 17**), we refined our strategy by recovering the N-terminal sequences of FPs and even incorporating additional Gly_2_Ser linkers. Using this approach, we first engineered a new blue mTagBFP2(GGS)-Fb_LAM2_ with a 5-fold improved difference between antigen-stabilized and destabilized forms (**Supplementary Figs. 15b, 16d, e, 17**). We then applied the refined strategy of recovering FP’s N-terminus and insertion of flexible linkers for engineering the far-red and blue Nbs for p24 and gp41 proteins, mCardinal-Fb_59H10_ and mTagBFP2-Fb_2E7_ (**Supplementary Figs. 18a, b and 19a, b**), achieving contrasts between the antigen-bound and destabilized states 110- and 76-fold, respectively (**Fig. 2a, b**). Next, we recovered the N- and C-termini of mTagBFP2 using the same approach and inserted it into Nb_ALFA_, resulting in a blue fluorescent VIS-Fb to the ALFA-tag peptide^48^ (**Fig. 2c**). We then swapped mTagBFP2 for mStayGold^49^, CyOFP1, TagRFP-T^50^, and mScarlet, creating green, orange, red, and far-red VIS-Fbs to ALFA-tag (**Fig. 2c, Supplementary Figs. 18c and 19c**). All VIS-Fbs were brightly fluorescent in the presence of their cognate antigen in HeLa cells (**Fig. 2a-c**). Thus, by aligning the N- and C-termini of various FPs with those of the FP that successfully functions with a specific Nb, we can create a set of fluorescent Nbs for a particular antigen by simply swapping the original FP for others.

**Figure 2.**
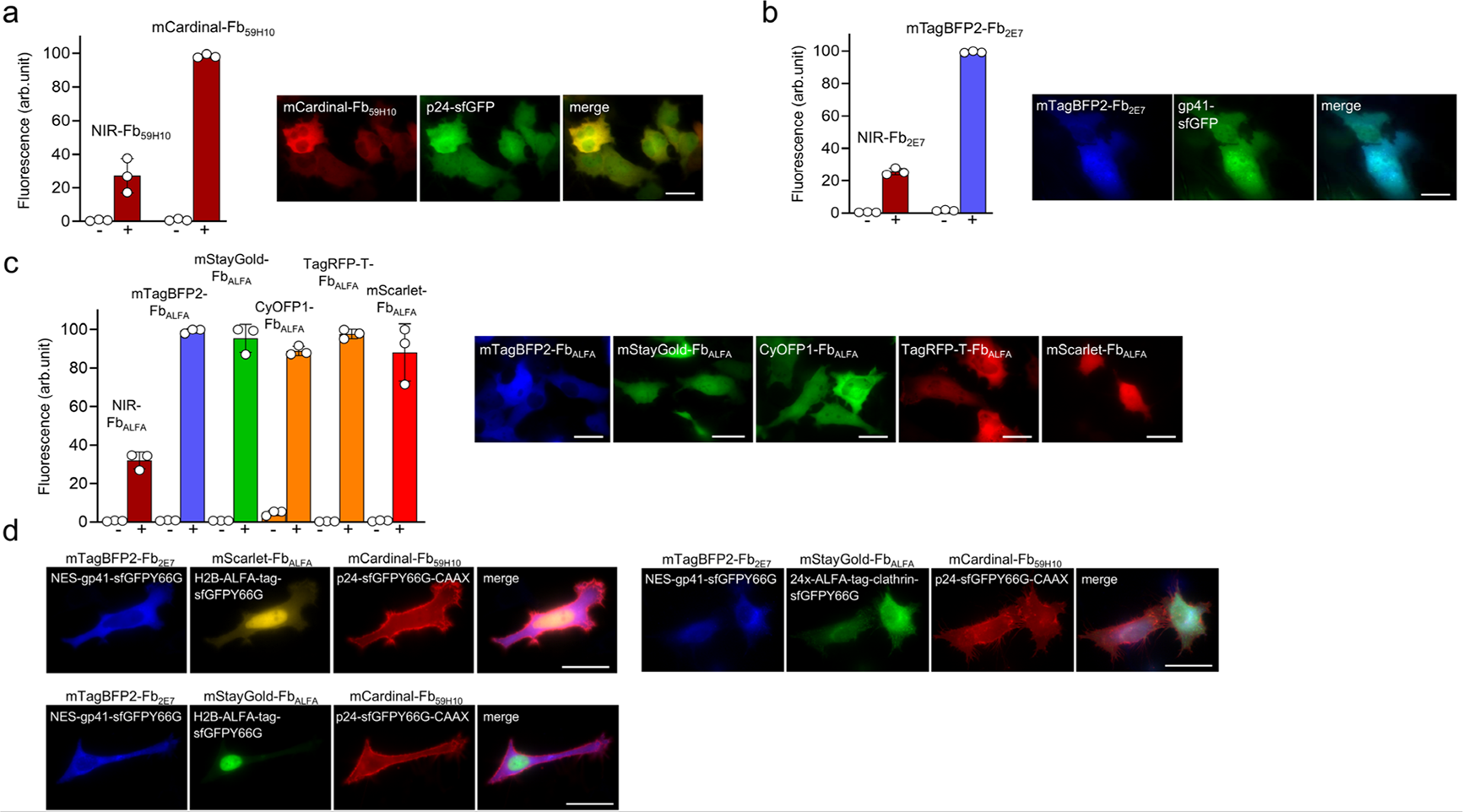
Multicolor imaging with three VIS-Fbs in HeLa cells. **(a)** Fluorescence intensity and images of cells expressing mCardinal-Fb_59H10_ or NIR-Fb_59H10_ for HIV gp41 protein. Co-transfection with a plasmid encoding gp41-sfGFP was used as a positive control (+), and co-transfection with pmEGFP-N1 was used as a negative control (−). **(b)** Fluorescence intensity and images of cells expressing mTagBFP2-Fb_2E7_ or NIR-Fb_2E7_ for HIV p24 protein. Co-transfection with a plasmid encoding p24-sfGFP was used as a positive control (+), and co-transfection with pmEGFP-N1 was used as a negative control (−). **(c)** Fluorescence intensity and images of cells expressing mTagBFP2-Fb_ALFA_, mStayGold-Fb_ALFA_, CyOFP1-Fb_ALFA_, TagRFP-T-Fb_ALFA_, mScarlet-Fb_ALFA_ or NIR-Fb_ALFA_ for ALFA tag. Co-transfection with a plasmid encoding 24x ALFA-tag repeats was used as a positive control (+), and co-transfection with pmEGFP-N1 was used as a negative control (−). **(d)** Multicolor images of HeLa cells co-expressing three different VIS-Fbs: mTagBFP2-Fb_2E7_, mScarlet-Fb_ALFA_/mStayGold-Fb_ALFA_ and mCardinal-Fb_59H10_ with their cognate antigens fused to nuclear exclusion signal, nuclear localization signal, clathrin or membrane localization signal, respectively: NES-gp41-sfGFPY66G, H2B-ALFA-tag-sfGFPY66G/24xALFA-tag-clathrin and p24-sfGFPY66G-CAAX. In (a-d) the following filters were used: for imaging mTagBFP2-Fb_2E7_ and mTagBFP2-Fb_ALFA_ 390/40 nm excitation and 460/40 nm emission; for imaging sfGFP and mStayGold-Fb_ALFA_ 480/40 nm excitation and 535/40 nm emission; for imaging CyOFP1-Fb_ALFA_ 523/20 nm excitation and 588/21 nm emission; for imaging TagRFP-T-Fb_ALFA_ and mScarlet-Fb_ALFA_ 575/25 nm excitation and 615/30 nm emission; for imaging mCardinal-Fb_59H10_ 605/30 nm excitation and 667/30 nm emission. Scale bar, 40 μm. In (a-c) fluorescence intensity was analyzed by flow cytometry using a 405 nm excitation laser and 450/50 nm emission filter for mTagBFP2-Fb_2E7_ and mTagBFP2-Fb_ALFA_; a 488 nm excitation laser and 525/50 nm emission filter for mEGFP, sfGFP and 510/20 nm emission filter for mStayGold-Fb_ALFA_, 582/15 nm emission filter for CyOFP1-Fb_ALFA_; a 561 nm excitation laser and 602/40 nm emission filter for mScarlet-Fb_ALFA_, 610/20 nm emission filter for TagRFP-T-Fb_ALFA,_ 670/30 nm emission filter for mCardinal-Fb_59H10_; a 640 nm excitation laser and a 660/20 nm emission filter for NIR-Fb_59H10_, NIR-Fb_2E7_ and NIR-Fb_ALFA_. (a, b) The maximal fluorescence of antigen-bound form for each VIS-Fb was assumed to be 100%. In (c) for VIS-Fb_ALFA_s, fluorescence of mTagBFP2-Fb_ALFA_ was assumed to be 100%. (a-c) Data are presented as mean values ± s.d. for *n* = 3 transfection experiments.

### Multicolor imaging of several antigens simultaneously

To evaluate the potential of targeting different antigens with VIS-Fbs, we co-expressed mTagBFP2-Fb_2E7_, mScarlet-Fb_ALFA_, and mCardinal-Fb_59H10_ constructs with gp41, ALFA-tag, and p24 cognate antigens, localized to the cytoplasm, nucleus, and plasma membrane of HeLa cells, respectively (**Fig. 2d**). For this, we fused HIV gp41 antigen with a NES signal, ALFA-tag with H2B nuclear localization tag, and HIV p24 antigen with a membrane-localization tag (CAAX). Alternatively, we used mStayGold_ALFA_ instead of mScarlet-Fb_ALFA_, co-expressed with 24x ALFA-tag repeats fused with the clathrin signal, resulting in mStayGold-Fb_ALFA_ localization in the coated vesicles (**Fig. 2d**). As a result, bright fluorescence of corresponding VIS-Fbs was observed in the membrane, cytoplasm, and nucleus (or coated vesicles) (**Fig. 2d**), demonstrating the ability to background-free detect several antigens in single cells using spectrally distinct VIS-Fbs. This capability should be extremely useful in complex biological systems with multiple markers, enabling multi-parameter cell mapping. Notably, achieving similar multi-antigen cell discrimination would be challenging with NIR-Fbs in terms of spectral resolution, as they operate in a narrow part of the NIR spectral range (670-718 nm)^8, 9, 51^.

### Fluorescent nanobodies with the ability to detect cellular metabolites

To determine whether Nb_GFP_ tolerates the insertion of not only GFP-like FPs but also larger constructs and to extend the number of Nbs in our engineering platform, we inserted a mApple-based calcium biosensor jRGECO1a^52^ into an anti-GFP Nb, LAG30 (Nb_LAG30_)^44^ (**Supplementary Fig. 20**). Nb_LAG30_ binds to GFP-derived FPs at a side opposite to that recognized by the Nb_GFP_ used to develop multicolor VIS-Fb above. Nb_LAG30_ binding does not block the β7–β8 loop region of the GFP’s β-can, which serves as the insertion point for metabolite-sensing domains in the GFP-based biosensors^53^. Since jRGECO1a consists of a circularly permuted mApple sandwiched between myosin light chain kinase at the N-terminus and calmodulin at the C-terminus, we did not incorporate any linkers when inserting jRGECO1a into Nb_LAG30_ (**Supplementary Fig. 20**). When co-expressed with mEGFP, the resulting jRGECO1a-Fb_LAG30_ construct exhibited dim red fluorescence, which intensified upon increasing calcium levels after adding 5 μM ionomycin (**Fig. 3a**).

**Figure 3.**
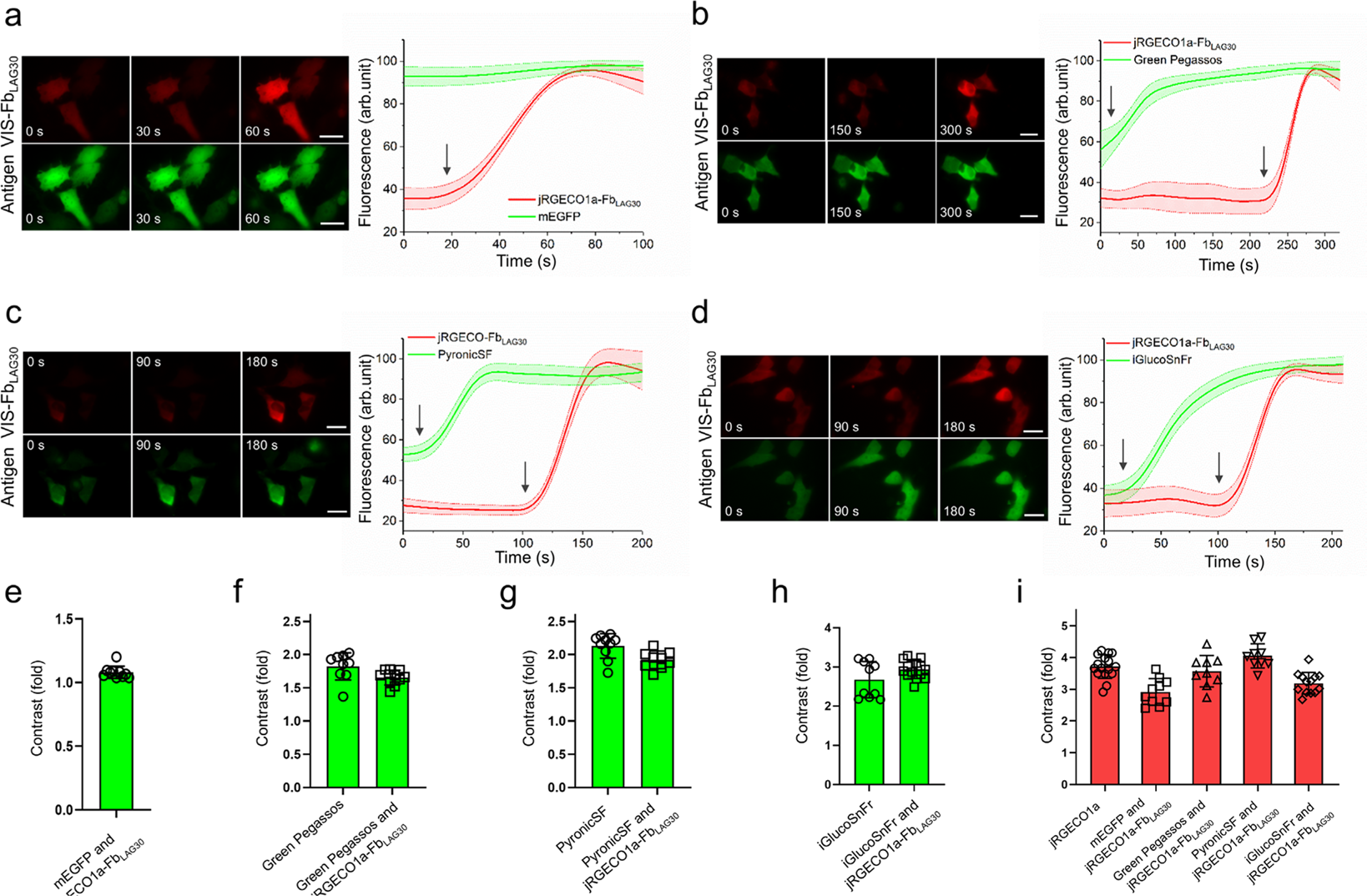
Simultaneous detection of different metabolites with jRGECO1a-Fb_LAG30_ in live GFP-expressing cells. **(a)** Change in fluorescence intensity of the cell co-expressing jRGECO1a-Fb_LAG30_ (red) and mEGFP (green) in response to 5 μM ionomycin. **(b)** Change in fluorescence intensity of the cell co-expressing jRGECO1a-Fb_LAG30_ (red) and Green Pegassos pyruvate biosensor (green) in response to 1 mM pyruvate. **(c)** Change in fluorescence intensity of the cell co-expressing jRGECO1a-Fb_LAG30_ (red) and PyronicSF pyruvate biosensor (green) in response to 10 mM pyruvate. **(d)** Change in fluorescence intensity of the cell co-expressing jRGECO1a-Fb_LAG30_ (red) and iGlucoSnFr biosensor (green) in response to 20 mM pyruvate. (a-d) Scale bar, 40 μm. **(e)** Contrast of mEGFP co-expressed with jRGECO1a-Fb_LAG30_ (*n=10*) after addition of 5 μM ionomycin. **(f)** Contrast of Green Pegassos only (*n=9*) and Green Pegassos co-expressed with jRGECO1a-Fb_LAG30_ (*n=9*) after addition of 1 mM pyruvate. **(g)** Contrast of PyronicSF only (*n=10*) and PyronicSF co-expressed with jRGECO1a-Fb_LAG30_ (*n=9*) after the addition of 10 mM pyruvate. **(h)** Contrast of iGlucoSnFr only (*n=11*) and iGlucoSnFr co-expressed with jRGECO1a-Fb_LAG30_ (*n=13*) after the addition of 20 mM glucose. **(i)** Contrast of jRGECO1a (*n=16*) and jRGECO1a-Fb_LAG30_ only (*n=10*) or jRGECO1a-Fb_LAG30_ co-expressed with Green Pegassos (*n=9*), PyronicSF (*n=9*) or iGlucoSnFr (*n=13*). In (b-d), the following filters were used: for imaging mEGFP, Green Pegassos, PyronicSF, and iGlucoSnFr 480/40 nm excitation and 535/40 nm emission; for imaging jRGECO1a-Fb_LAG30_ 575/25 nm excitation and 615/30 nm emission.

We then co-expressed jRGECO1a-based VIS-Fb_LAG30_ with GFP-based genetically encoded biosensors for pyruvate (GreenPegassos^54^ and PyronicSF^55^) and glucose (iGlucoSnFr^56^). Without stimulation, for all three biosensors, the dim fluorescence of jRGECO1a-Fb_LAG30_ demonstrated overlap with the green fluorescence of the corresponding co-transfected biosensor (**Fig. 3b-d**). We then added respective metabolites (pyruvate or glucose) to the mammalian cultures and observed a fast increase in the green fluorescence for each biosensor (**Fig. 3b-d**). Simultaneously, we stimulated calcium release from intracellular compartments and recorded an increase in red fluorescence of jRGECO1a-Fb_LAG30_ (**Fig. 3b-d**). The performance of jRGECO1a, as well as the fluorescence kinetics of the antigen biosensors (Green Pegassos, PyronicSF, and iGlucoSnFr), were not substantially affected by insertion into Nb_LAG30_ or co-expression with jRGECO1a-Fb_LAG30_ (**Supplementary Fig. 21**), respectively. We observed no difference between the green fluorescence of biosensors only or biosensors co-expressed with jRGECO1a-Fb_LAG30_ (**Fig. 3e-h**). Similarly, there were no changes in red fluorescence of jRGECO1a or jRGECO1a-Fb_LAG30_ co-expressed with either mEGFP or GFP-based biosensors (**Fig. 3i**).

This indicates that by utilizing our VIS-Fb technology, we could bring two different genetically encoded biosensors into proximity, potentially allowing for simultaneous real-time monitoring of both metabolites within overlapping cellular regions. However, further quantitative and high-resolution analysis is required to confirm precise stoichiometry. Moreover, these results suggest the possibility of inserting other non-circular-permuted biosensors with original N- and C-termini into Nbs. These advances are impossible with NIR-Fbs because of the lack of GAF-domain-based NIR biosensors.

### Fluorescent nanobody for selective labeling and imaging of neuronal soma

Inspired by our success in the orthogonal labeling of the GreenPegassos, PyronicSF, and iGlucoSnFr biosensors with Nb_LAG30_ while preserving their functionality, we used another anti-GFP Nb, LAG16 (Nb_LAG16_)^44^ to target GFP biosensors with constitutively red fluorescent dTomato. Nb_LAG16_ recognizes the GFP’s β-can at an epitope close to that of Nb_LAG30_, thus preserving the biosensors’ functionality^53^. dTomato was selected due to the relative ease of its two-photon imaging with a standard Ti:Sapphire laser rather than an optical parametric oscillator. We tested dTomato insertion at Nb_LAG16_ position Ser65/Val66 with no linkers or Gly_2_Ser flexible linkers. The insertion with Gly_2_Ser linkers had the highest contrast between antigen-bound and destabilized forms (**Fig. 4a, Supplementary Fig. 22**). We then co-expressed the dTomato-Fb_LAG16_ construct with mEGFP (**Fig. 4b**) and calcium biosensor GCaMP6s^57^ and monitored fluorescence changes in response to the calcium increase (**Fig. 4c, d**). A fast increase in green fluorescence of GCaMP6s was observed while there were no changes in red fluorescence of dTomato-Fb_LAG16_ (**Fig. 4d**). There was no difference between green fluorescence of GCaMP6s only or GCaMP6s co-expressed with dTomato-based VIS-Fb_LAG16_ (**Fig. 4e**).

**Figure 4.**
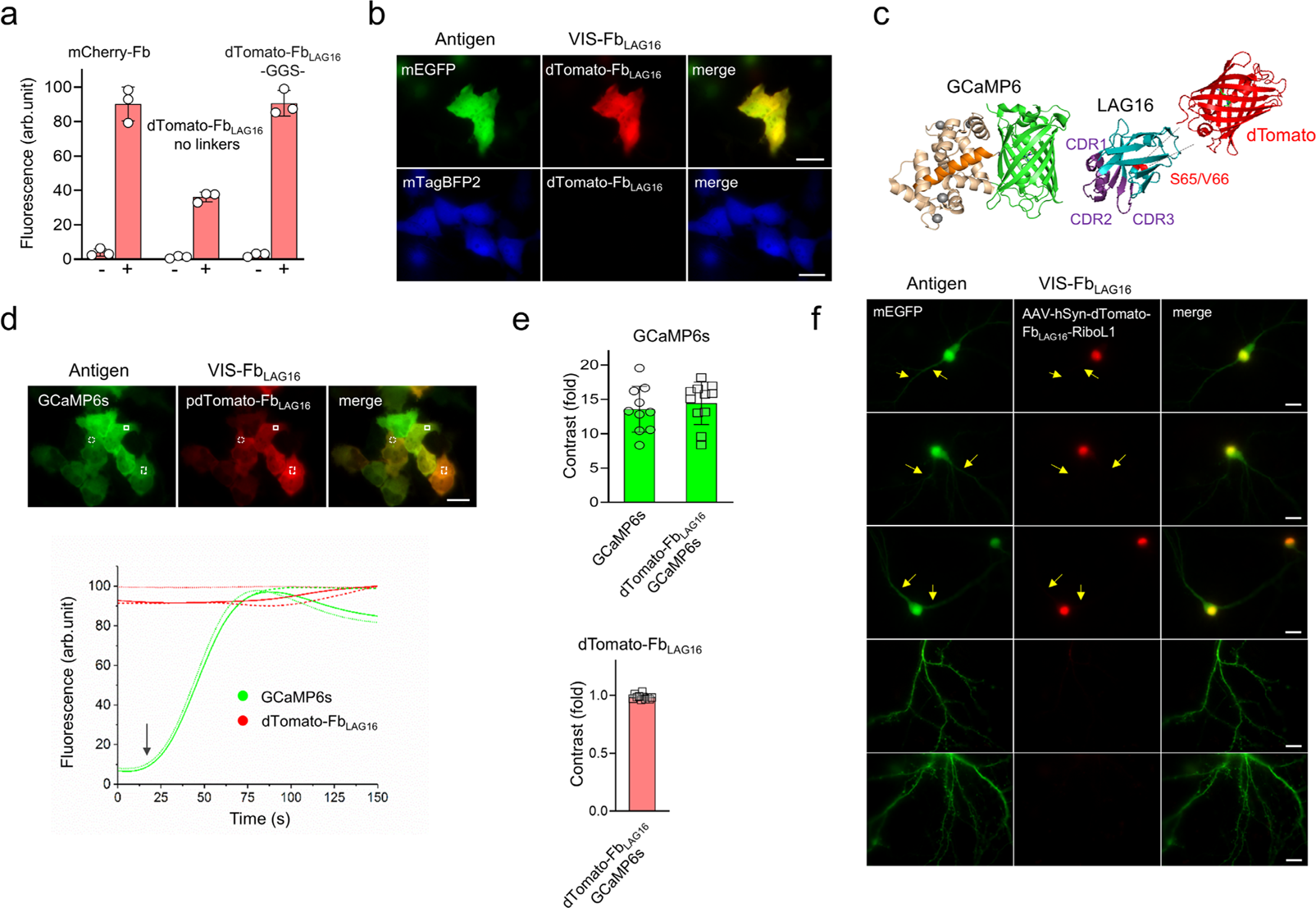
Performance of dTomato-Fb_LAG16_ in HeLa cells and cultured primary neurons. **(a)** Red fluorescence intensity of cells transfected with mCherry-Fb_GFP_, dTomato-Fb_LAG16_ without linkers, or dTomato-Fb_LAG16_ with –GGS-linkers and co-expressed with mEGFP (right column (+)) or mTagBFP2 (left column (−)). **(b)** Fluorescence images of HeLa cells co-expressing dTomato-Fb_LAG16_ with mTagBFP2 (negative control) or mEGFP (positive control). **(c)** Scheme of a VIS-Fb with a red FP (PDB ID: 1ZGO) inserted into LAG16 anti-GFP nanobody (PDB ID: 6LR7) bound to GFP-based biosensor GCaMP6m (PDB ID: 3WLD). Complementarity-determining regions (CDRs) are highlighted in violet. The position of dTomato insertion to the anti-GFP nanobody is indicated with a red arrow. **(d)** Upper, representative image of HeLa cells co-expressing GCaMP6s and dTomato-Fb_LAG16_. Three regions of interest (ROIs) are indicated with white squares. Lower, changes in fluorescence intensity of the same cell co-expressing GCaMP6s (green) and dTomato-Fb_LAG16_ (red) in response to 5 μM ionomycin. Fluorescence changes for three ROIs are shown. **(e)** Upper, contrast of GCaMP6s only (*n=10*) and GCaMP6s co-expressed with dTomato-Fb_LAG16_ (*n=11*) after addition of 5 μM ionomycin. Lower, contrast of dTomato-Fb_LAG16_ (*n=11*) for the data presented in the left graph. **(f)** Co-expression of dTomato-Fb_LAG16_ fused to RiboL1 tag and mEGFP in the soma of hippocampal neurons. In (a) fluorescence intensity was analyzed by flow cytometry using a 405 nm excitation laser and 450/50 nm emission filter for mTagBFP2; a 488 nm excitation laser and 525/50 nm emission filter for mEGFP; a 561 nm excitation laser and 610/20 nm emission filter for mCherry-Fb_GFP_ and dTomato-Fb_LAG16_. The maximal fluorescence of antigen-bound form for dTomato(GGS)-Fb_LAG16_ was assumed to be 100%. Data are presented as mean values ± s.d. for *n* = 3 transfection experiments. In (b, d, and f), the following filters were used: for imaging mEGFP and GCaMP6s 480/40 nm excitation and 535/40 nm emission; for imaging dTomato-Fb_LAG16_ and dTomato-Fb_LAG16_-RiboL1 575/25 nm excitation and 615/30 nm emission. (b, d) Scale bar, 40 μm. (f) Scale bar, 20 μm.

To determine whether dTomato-Fb_LAG16_ tolerates soma-localization tags, we fused it with several peptide signals known to localize proteins to the neuronal soma: ribo-tags RiboL1^58^ and RPL10^59^, and coiled-coil peptide EE-RR^60^. All soma-localization peptides were fused to the C-terminus of the dTomato-Fb_LAG16._ We screened the engineered constructs in hippocampal mouse neuronal cultures co-transfected with mEGFP (**Fig. 4f, Supplementary Fig. 23**). dTomato-Fb_LAG16_ fused with RiboL1 targeted the neuronal soma with high precision and was selected for further *in vivo* experiments. In contrast, the dTomato-Fb_LAG16_ fused with RPL10 and EE-RR signals showed prominent red fluorescence in the neuronal processes (**Supplementary Fig. 23**).

### Intersectional labeling and functional imaging in behaving mice

To assess the effectiveness and versatility of VIS-Fbs for intersectional labeling and ratiometric calcium imaging of various cell compartments and populations, we conducted two-photon imaging in mice expressing the calcium indicator GCaMP6f and dTomato-Fb_LAG16_ under different neuronal and glial promoters.

We first delivered an adeno-associated viral (AAV) vector driving dTomato-Fb_LAG16_ expression under the control of the hSyn promoter and the soma-targeting peptide RiboL1^58^ into the somatosensory cortex of adult Thy1-GCaMP6f mice, which preferentially express the calcium indicator in a subset of excitatory pyramidal neurons (**Fig. 5a**). As expected, dTomato-Fb_LAG16_ expression was mainly confined to neuronal cell bodies (**Fig. 5b**, showing 91.2% ± 7.0% colocalization with GCaMP6f expression and 98.1% ± 1.9% with NeuN immunostaining (**Fig. 5b** and **Supplementary Fig. 24a-d**; see also **Methods**). Dual-color two-photon imaging in spontaneously active Thy1-GCaMP6f mice on a spherical treadmill revealed that dTomato-Fb_LAG16_ fluorescence remained stable across ~10-min recording periods (**Supplementary Fig. 25a-c**). This stability enabled ratiometric imaging (**Fig. 5c**), which can improve the signal-to-noise ratio of calcium transients. The calcium spiking in dTomato-positive and -negative GCaMP6f-expressing neurons was comparable (**Fig. 5c**, center), indicating that VIS-Fb binding does not significantly interfere with GCaMP conformational changes, consistent with our previous work^9^.

**Figure 5.**
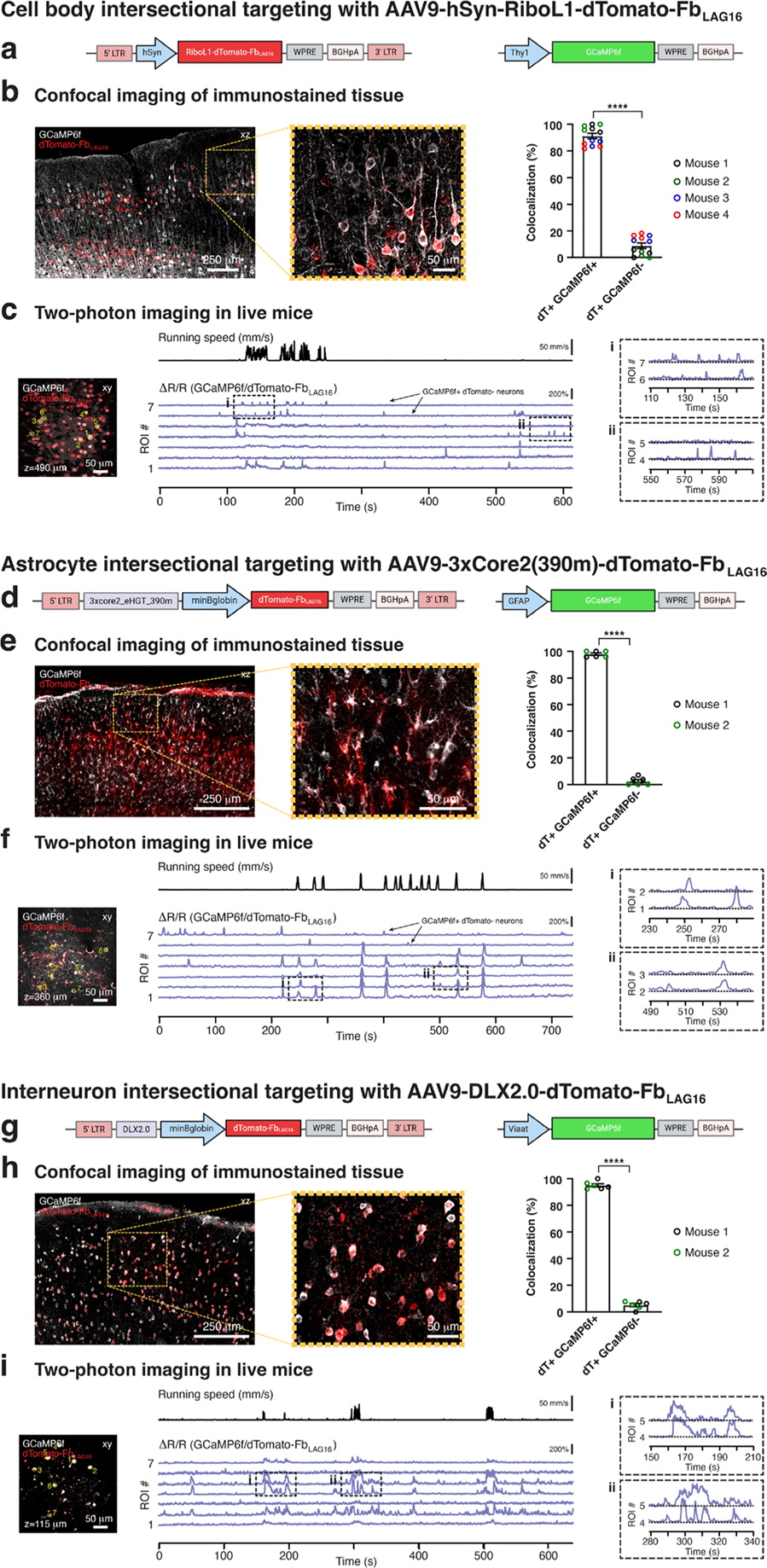
Intersectional targeting of specific cell compartments and populations with dTomato-Fb_LAG16_ in GCaMP6f reporter mice. **(a)** Schematic of the experimental approach. An AAV vector driving dTomato-Fb_LAG16_ expression under the control of the hSyn promoter and the soma-targeting peptide RiboL1 was stereotactically injected into the somatosensory cortex of Thy1-GCaMP6f mice with preferential calcium indicator expression in a subset of excitatory pyramidal neurons. **(b)** Immunostaining validation. Left, example confocal fluorescence images showing GCaMP6f-(gray) and dTomato-expressing cells (red) in a cortical tissue section from an injected *Thy1*-GCaMP6f mouse. Center, zoom-in of the indicated region. Scale bars, 250 μm (left) and 50 μm (center). Right, population analysis (*n=12* tissue sections from four mice). Data are presented as mean values ± SD. **(c)** *In vivo* validation. Left, example two-photon fluorescence image from a dual-color time-lapse recording showing GCaMP6f-(gray) and dTomato-expressing cells (red) in the somatosensory cortex of a behaving Thy1-GCaMP6f mouse. Recording depth (z) from the pial surface and seven somatic regions of interest (ROIs) is indicated. Center, fluorescence transients in the indicated ROIs are shown as ΔR/R (blue) for the combined channels. The simultaneously recorded mouse’s locomotor activity on a spherical treadmill is shown above the fluorescence traces. Scale bars, 50 μm (left), 50 mm/s and 200% (center). Right, zoom-ins of the two periods indicated in (c, center). **(d)** Schematic of the experimental approach. An AAV vector driving dTomato-Fb_LAG16_ expression under the control of the astrocyte enhancer 3xCore2(390m) was stereotactically injected into the somatosensory cortex of GFAP-GCaMP6f mice with preferential calcium indicator expression in astrocytes. **(e)** Immunostaining validation. Left, example confocal fluorescence images showing GCaMP6f-(gray) and dTomato-expressing cells (red) in a cortical tissue section from an injected *GFAP*-GCaMP6f mouse. Center, zoom-in of the indicated region. Scale bars, 250 μm (left) and 50 μm (center). Right, population analysis (*n=6* tissue sections from two mice). Data are presented as mean values ± SD. **(f)** *In vivo* validation. Left, example two-photon fluorescence image from a dual-color time-lapse recording showing GCaMP6f-(gray) and dTomato-expressing cells (red) in the somatosensory cortex of a behaving GFAP-GCaMP6f mouse. Recording depth (z) from the pial surface and seven somatic regions of interest (ROIs) is indicated. Center, fluorescence transients in the indicated ROIs are shown as ΔR/R (blue) for the combined channels. The simultaneously recorded mouse’s locomotor activity on a spherical treadmill is shown above the fluorescence traces. Scale bars, 50 μm (left), 50 mm/s and 200% (center). Right, zoom-ins of the two periods indicated in (f, center). **(g)** Schematic of the experimental approach. An AAV vector driving dTomato-Fb_LAG16_ expression under the control of the DLX2.0 enhancer was stereotactically injected into the somatosensory cortex of Viaat-GCaMP6f mice with calcium indicator expression in inhibitory interneurons. **(h)** Immunostaining validation. Left, example confocal fluorescence images showing GCaMP6f-(gray) and dTomato-expressing cells (red) in a cortical tissue section from an injected *Viaat*-GCaMP6f mouse. Center, zoom-in of the indicated region. Scale bars, 250 μm (left) and 50 μm (center). Right, population analysis (*n=6* tissue sections from two mice). Data are presented as mean values ± SD. **(i)** *In vivo* validation. Left, example two-photon fluorescence image from a dual-color time-lapse recording showing GCaMP6f-(gray) and dTomato-expressing cells (red) in the somatosensory cortex of a behaving Viaat-GCaMP6f mouse. Recording depth (z) from the pial surface and seven somatic regions of interest (ROIs) is indicated. Center, fluorescence transients in the indicated ROIs are shown as ΔR/R (blue) for the combined channels. The simultaneously recorded mouse’s locomotor activity on a spherical treadmill is shown above the fluorescence traces. Scale bars, 50 μm (left), 50 mm/s and 200% (center). **(i)** Zoom-ins of the two periods indicated in (i, center).

Next, we delivered an AAV vector to drive dTomato-Fb_LAG16_ expression under the control of the astrocyte enhancer 3xCore2(390m)^61^ into the somatosensory cortex of adult GFAP-GCaMP6f mice (**Fig. 5d**). These mice preferentially express the calcium indicator in astrocytes, with some off-target expression in neurons. We observed that 98.0% ± 2.2% of dTomato-positive cells colocalized with GCaMP6f, while 93.9% ± 2.4% colocalized with Sox9, an astrocyte marker (**Fig. 5e** and **Supplementary Fig. 24e-i**; see also **Methods**). In contrast, 2.4% ± 1.7% colocalized with NeuN immunostaining (**Supplementary Fig. 24i**). This demonstrates that our intersectional approach successfully distinguished between GCaMP6f-expressing astrocytes and neurons in this mouse line. Further, dual-color two-photon imaging in spontaneously active GFAP-GCaMP6f mice on a spherical treadmill showed that dTomato-negative GCaMP6f-expressing cells exhibited neuron-like morphology and calcium spiking with spatiotemporal features that were distinct from the astrocytic syncytium calcium transients triggered by running behavior^62, 63^ (**Fig. 5f**, **Supplementary Fig. 25d-f**).

Finally, we delivered an AAV vector driving dTomato-Fb_LAG16_ expression under the control of the DLX2.0 enhancer^64^, targeting GABAergic interneurons, into the somatosensory cortex of adult Viaat-GCaMP6f mice that express calcium indicators in all inhibitory interneurons (both GABAergic and glycinergic) (**Fig. 5g**). We found that 94.9% ± 2.9% of dTomato-positive cells colocalized with GCaMP6f and 97.7% ± 1.9% colocalized with NeuN immunostaining (**Fig. 5h** and **Supplementary Fig. 24j-m**; see also **Methods**). This suggests that our approach effectively targets GABAergic interneurons in the adult cortex. Dual-color two-photon imaging in spontaneously active Viaat-GCaMP6f mice on a spherical treadmill demonstrated that dTomato-Fb_LAG16_ expression enabled high-fidelity ratiometric measurements of interneuron calcium spiking (**Fig. 5i**, **Supplementary Fig. 25g-i**).

Collectively, these results reveal the high potential of VIS-Fbs for multiplexed, antigen-dependent functional imaging across various cell types in behaving mice.

### Real-time tracking of endogenous β-catenin in early embryogenesis

Currently, only a few well-characterized nanobodies, antibodies, or other small fluorescent tools can specifically recognize endogenous proteins in zebrafish *in vivo* with high affinity. To address this limitation, we applied the VIS-Fb technology to engineer a new antigen-stabilized fluorescent Nb against β-catenin based on BC2 Nb^65^. For this, we inserted green sfGFP between Ser64 and Val65 of BC2 Nb via Ser_2_Gly linkers (**Supplementary Fig. 26**)^65^. BC2 Nb has a high affinity to β-catenin (1.9 nM) and recognizes a linear peptide at its N-terminal domain. Importantly, while the central armadillo repeat domain and the C-terminal domain of β-catenin interact with numerous binding partners, the N-terminal domain has relatively few interactions^66^. Thus, VIS-Fb_BC2_ offers a particular advantage by enabling β-catenin targeting and imaging with minimal perturbation of its function and localization.

Validation of the sfGFP-Fb_BC2_ construct in mammalian cultured cells co-expressing β-catenin showed that the sfGFP-Fb_BC2_ was brightly fluorescent and antigen-stabilized. In contrast, no fluorescence was observed when blue mTagBFP2 was co-expressed, as these constructs were destabilized (**Supplementary Fig. 27a**). Expression of sfGFP-Fb_BC2_ was predominantly nuclear in cultured mammalian cells (**Supplementary Fig. 27b**), consistent with previous findings that BC2 Nb precipitates more β-catenin from nuclear than from cytoplasmic fractions^65^.

The β-catenin plays a critical role both in cell signaling, as a key protein in the Wnt signaling pathway, and in cell adhesion, as a component of adhesion complexes^66^. To assess whether our engineered, antigen-stabilized VIS-Fb_BC2_ targets endogenous β-catenin *in vivo*, we expressed it in a mosaic way in wild-type *Danio rerio* (zebrafish) embryos under the control of the constitutive Ef1a promoter. We injected pTol2 plasmid encoding sfGFP-Fb_BC2_ under Ef1a promoter into zebrafish eggs at the one-cell stage. Embryos were screened for fluorescence from 5 h post-injection (hpi) until 72 hpi in at least two biological replicates with over 200 eggs injected in each experiment. Fluorescence was not observed between 5-8 h post-injection. At 24 hpi, zebrafish embryos showed mosaic expression of sfGFP-Fb_BC2_ for β-catenin in cells at different regions and tissues, such as head, dorsal trunk, skin epithelium, eyes, muscle fibers, and neural tube (**Fig. 6a, b and Supplementary Fig. 28**).

**Figure 6.**
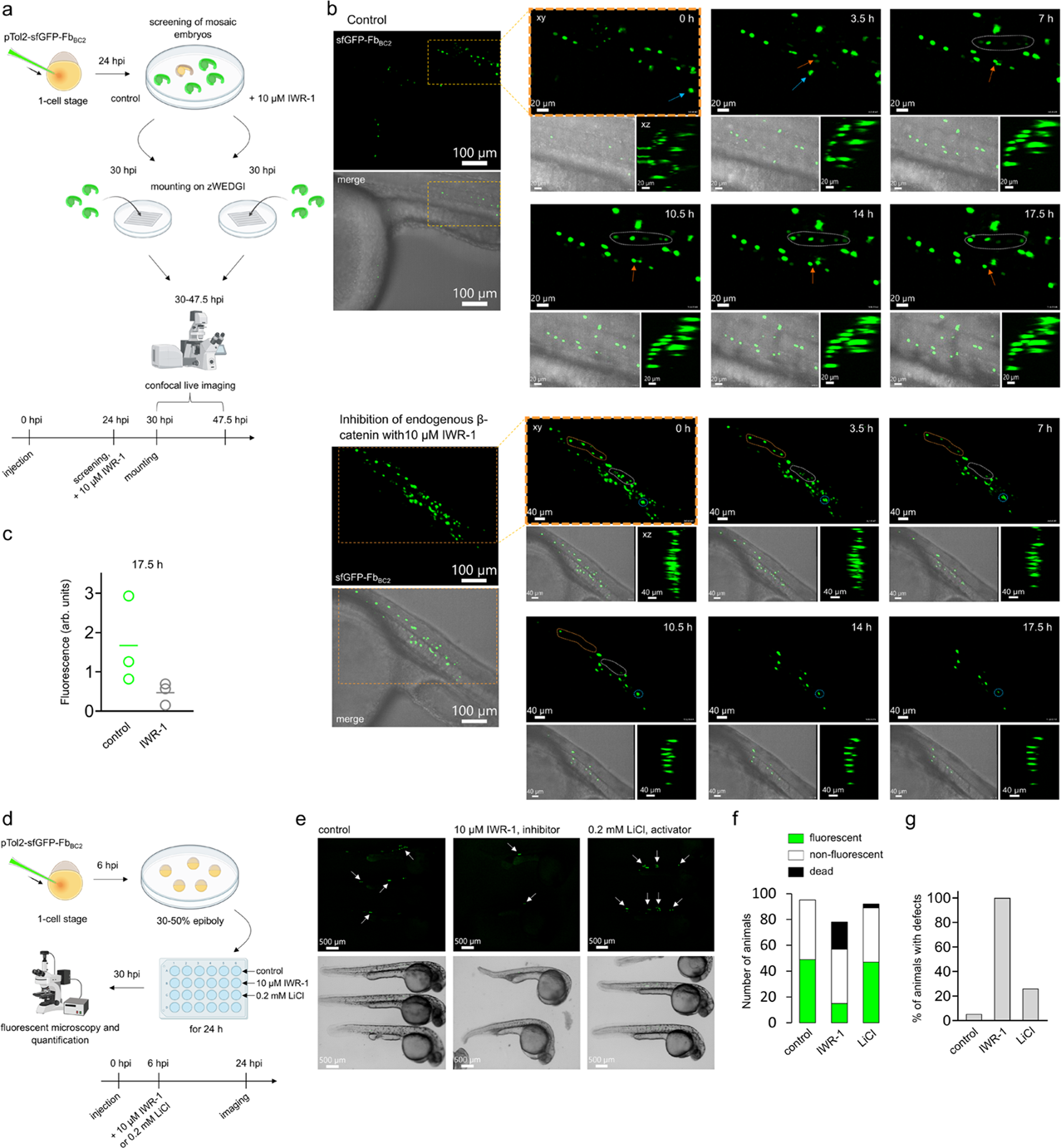
*In vivo* tracking of endogenous β-catenin in zebrafish. **(a)** A scheme of experiment: the pTol2-sfGFP-Fb_BC2_ plasmid encoding VIS-Fb for β-catenin was injected into wild-type zebrafish embryos at the one-cell stage. At 24 hours post-injection (hpi), larvae were screened for green fluorescence and then divided into two groups. One group remained untreated (control), while the other one was incubated with 10 μM of β-catenin inhibitor IWR-1 for 6 h. At 30 hpi, larvae exhibiting mosaic expression of sfGFP-Fb_BC2_ were mounted on a zWEDGI chamber and imaged using a spinning disk confocal microscope for up to 17.5 h. Zebrafish pre-treated with 10 μM IWR-1 were also imaged in continuous presence of 10 μM IWR-1. **(b)** Upper part, 3D view time-lapsed images from intravital spinning disk fluorescent confocal microscopy of 30 hpi wild-type untreated zebrafish larvae previously injected at the one-cell stage with pTol2-sfGFP-Fb_BC2_ for β-catenin. Time-lapse images acquired every 3.5 h are presented. Shown are zoom-ins for xy and xz projections for the sfGFP-Fb_BC2_ channel, as well as xy projection of the merged brightfield and green fluorescence channels. Dashed white ROIs and orange arrows indicate cells displaying an increase in endogenous β-catenin signal over time. Blue arrows highlight a migrating cell. Lower part, 3D view time-lapsed images from intravital spinning disk fluorescent confocal microscopy of 30 hpi wild-type zebrafish larvae pre-treated with 10 μM of β-catenin inhibitor IWR-1, previously injected at the one-cell stage with pTol2-sfGFP-Fb_BC2_ for β-catenin. Time-lapse images acquired every 3.5 h are presented. Shown are zoom-ins for xy and xz projections for the sfGFP-Fb_BC2_ channel and xy projection of the merged brightfield and green fluorescence channels. Dashed white, orange, and blue ROIs indicate cells showing a decrease in endogenous β-catenin signal over time in response to IWR-1 treatment. **(c)** Fluorescence intensity of sfGFP-Fb_BC2_ was quantified in control (untreated) zebrafish (*n=3*) and IWR-1–treated zebrafish (*n=3*) after 17.5 h of live imaging. Intensities at 17.5 h were normalized to the corresponding values at 0 h for each group. **(d)** A scheme of experiment: the pTol2-sfGFP-Fb_BC2_ plasmid encoding VIS-Fb for β-catenin was injected into wild-type zebrafish embryos at the one-cell stage. At 6 hpi, larvae were divided into three groups in 24-well plate: control (untreated), treated with 10 μM of β-catenin inhibitor IWR-1 or 0.2 mM of β-catenin activator LiCl. After 24 h of incubation, zebrafish larvae were imaged using fluorescent microscope. **(e)** Representative fluorescent images of 24 hpi larvae with mosaic expression of sfGFP-Fb_BC2_, untreated, treated with 10 μM IWR-1 or 0.2 mM LiCl. White arrows indicate β-catenin-expressing cells. **(f)** Proportion of larvae exhibiting fluorescence, no fluorescence, and mortality in each treatment condition (control, 10 μM IWR-1 and 0.2 mM LiCl). **(g)** Percentage of zebrafish larvae with developmental defects across treatment conditions (control, 10 μM IWR-1 and 0.2 mM LiCl). Scalebar, (b) 100 μm, 20 μm, 40 μm (e) 500 μm. (a, d) Created with BioRender.com.

To validate if VIS-Fb_BC2_ can be used as a specific *in vivo* reporter of β-catenin dynamics during normal embryonic development and upon modulation of Wnt/β-catenin signaling pathway, zebrafish embryos injected with pTol2-sfGFP-Fb_BC2_ at the one-stage cell were screened for green fluorescence at 24 hpi and divided into two groups (**Fig. 6a**). One group remained untreated (control), while the other one was treated with 10 μM IWR-1 for 6 h (**Fig. 6a**). IWR-1 targets the Wnt/β-catenin signaling pathway by stabilizing the complex responsible for β-catenin degrading, leading to reduced β-catenin levels and suppression of Wnt signaling^67^. At 30 hpi, embryos from both groups showed strong mosaic fluorescence across various regions (**Fig. 6b** and **Supplementary Figs. 28-29**).

Long-term intravital fluorescent confocal microscopy was performed at 30 hpi for up to 17.5 h in 8-15 zebrafish embryos per group, mounted on a zWEDGI device. A representative time-lapse image series showing sfGFP-Fb_BC2_ targeting endogenous β-catenin in the trunk of control and IWR-1-treated zebrafish is presented in **Figure 6b** (see also **Supplementary Fig. 28 and 29**). During the 17.5 h imaging period of untreated embryos, we observed bright β-catenin-expressing cells within the trunk and in the epidermis, likely corresponding to muscle- and neural-derived cells, as well as keratinocyte-like cells. These cells displayed stable fluorescence with no signs of photobleaching (**Fig. 6b**, upper panel). Using sfGFP-Fb_BC2_, we detected the appearance of additional β-catenin-expressing cells along the developing spinal cord in real time *in vivo* (**Fig. 6b**, upper panel; white dashed ROI and orange arrow**; Supplementary Fig. 28**), while other cells – presumably of blood origin such as hematopoietic stem cells – were observed migrating over time (**Fig. 6a, upper panel; blue arrow**). While imaging zebrafish embryos treated with 10 μM IWR-1 inhibitor, we observed a continuous and rapid loss of the endogenous β-catenin signal, consistent with the known mechanism of IWR-1, which promotes β-catenin degradation (**Fig. 6b**, lower panel; orange, white, and blue dashed ROIs**; Supplementary Fig. 29**). We next compared the fluorescence intensity in control and IWR-1-treated zebrafish embryos at 17.5 h, with each value normalized to the corresponding fluorescence at 0 h. Treatment with the IWR-1 inhibitor resulted in a 3.5-fold decrease in β-catenin levels targeted by sfGFP-Fb_BC2_ (**Fig. 6c**).

While treatment of zebrafish embryos at 24 hpi is more suitable for targeting specific, later-stage effects on β-catenin patterning, we next asked if sfGFP-Fb_BC2_ could also be used to study Wnt/β-catenin signaling pathway during earlier stages of embryonic development. To test this, we injected zebrafish embryos with pTol2-sfGFP-Fb_BC2_ at the one-stage cell and randomly divided them into three groups at the stage of 30-50% epiboly (~6 hpi, before fluorescent signal is observed) in a 24-well plate: i) control (untreated) embryos, ii) embryos treated with 10 μM β-catenin inhibitor IWR-1, and iii) embryos treated with 0.2 mM β-catenin activator LiCl (**Fig. 6d**). After 24 h of incubation, zebrafish were imaged using fluorescence microscopy (**Fig. 6e**). Morphological analysis revealed typical defects that were especially pronounced in the IWR-1-treated embryos, including a shortened posterior body axis, curved tails, yolk sac edema, and reduced pigmentation (**Fig. 6e**). Regarding targeted β-catenin, IWR-1 treatment resulted in the fewest embryos with detectable fluorescence (**Fig. 6f**). Additionally, IWR-1 treatment caused increased embryos mortality and a higher percentage of animals with morphological defects (**Fig. 6f and g**). While the number of embryos with detectable fluorescence was comparable in the control and LiCl-treated group (**Fig. 6f**), we observed an increased number of β-catenin-positive cells in embryos treated with LiCl (**Fig. 6e**).

These results establish sfGFP-Fb_BC2_ as a non-invasive *in vivo* reporter for endogenous β-catenin and demonstrate normal and pharmacologically-induced dynamics of β-catenin in zebrafish embryogenesis, uncovering spatiotemporal signaling patterns not previously accessible with existing Wnt/β-catenin tools.

## Discussion

By genetically inserting GFP-like FPs into eight different Nbs frameworks, we developed a series of fully genetically encoded VIS-Fb binders with a molecular weight of ~42 kDa (**Fig. 1, Supplementary Fig. 8**). Overall, we have developed more than twenty antigen-stabilizable VIS-Fbs with emissions ranging from ~450 nm to ~660 nm and various photochemical properties. Importantly, VIS-Fbs fully preserve properties of the parental FPs, allowing engineering of not only constitutively fluorescent but also photoactivatable and photoconvertible VIS-Fbs, as well as VIS-Fbs with FP-based biosensors (**Figs. 1f-k, 3**).

It has been suggested that insertion of a GAF-domain-based NIR FP, having rigid α-helical termini, into an Nb may cause a stretch in the Nb structure, leading to the exposure of hydrophobic patches on the protein surface in an antigen-free state^8, 9^. Consequently, these patches are recognized by ubiquitinases that label NIR-Fbs for proteasomal degradation. We hypothesize that our engineered linkers between GFP-like FPs and Nbs likewise induce structural tension in the Nb scaffold, resulting in a similar dependence on NIR-Fb antigen. Upon binding to their cognate antigens, both VIS-Fbs and NIR-Fbs likely undergo conformational changes, which relieve the structural tension, burying the hydrophobic patches and stabilizing the overall structure. However, future structural studies are required to clarify the mechanism of antigen-dependent stabilization. Importantly, the demonstrated possibility to use proteins other than GAF-domain-based NIR FPs as destabilizing components in the Nb scaffold opens opportunities for inserting a wide range of other functional proteins into Nbs, including luciferases, enzymes, growth factors, and transcriptional activators. The activity of these proteins inserted into the resulting antigen-dependent nanobodies can be modulated by the presence of the cognate antigens.

Different GFP-like FPs used for the VIS-Fbs engineering exhibit notable variations in the length and amino acid composition of their N- and C-termini (**Supplementary Fig. 7**). By aligning the N- and C-terminal amino acids, we determined the consensus N- and C-termini required to engineer antigen-dependent VIS-Fbs (**Supplementary Figs. 7, 15, 18**). For example, to create a lineage of VIS-Fbs to GFP, we first optimized the termini of mCherry (**Supplementary Fig. 3**), resulting in a red fluorescent VIS-Fb. Then, by aligning mCherry termini with 14 other FPs, such as mTagBFP2, mTFP1, mWasabi, mNeonGreen, mOrange, LSSmOrange, CyOFP1, mScarlet, mScarlet-I, LSSmScarlet, mCardinal, mNeptune2, mEos4a, PAmCherry (**Supplementary Fig. 7**), we engineered fourteen other VIS-Fbs of different colors and photochemical properties to the same antigen (**Fig. 1b, c**). This VIS-Fb palette was then applied to multicolor imaging (**Fig. 1d, e**), showcasing the potential of VIS-Fbs for a single antigen to study multiple intracellular structures in live cells.

Applying a similar linker design principle, we next developed multicolor VIS-Fbs for p24, gp41, ALFA-tag, and mCherry (**Fig. 2**). For example, to engineer multicolor VIS-Fbs against ALFA-tag, we first restored the N- and C-termini for mTagBFP2 and introduced Gly_2_Ser linkers between FP and Nb, resulting in a blue fluorescent VIS-Fb for ALFA-tag. Then we swapped the mTagBFP2 for mStayGold, CyOFP1, TagRFP-T, and mScarlet using the same consensus N- and C-linkers found for mTagBFP2, resulting in the green, orange, red, and far-red VIS-Fbs to ALFA-tag, respectively (**Fig. 2c, Supplementary Figs. 18c, 19**). Consequently, we achieved multicolor imaging of several different antigens in live cells, demonstrating the versatility of VIS-Fbs (**Fig. 2d**).

For the standardized N- and C-termini, first introduced in mCherry and then in many other GFP-like FPs listed above, we identified generalizable principles of their modification, allowing us to copy and paste these FPs into Nbs. Once the N- and C-termini are optimized for a VIS-Fb for one member of this FP subclass, making VIS-Fbs for other FP subfamily members is a straightforward process. Moreover, all FPs have been inserted into the same Ser65/Val66 position of the third FR region of eight different Nbs. Due to the high homology between the FRs of various Nbs, once the N- and C-termini of an FP are designed to make one VIS-Fb, the same termini can be used to develop other VIS-Fbs. For example, mTagBFP2 with the termini optimized for an Nb to gp41 was then directly transferred into an Nb to ALFA-tag, resulting in the efficient VIS-Fb_ALFA_ (**Fig. 2a-c, Supplementary Figs. 18, 19**). Therefore, we implemented a generalized improvement by restoring the native N- and C-termini of the FPs and introducing flexible Gly_3_Ser linkers between the FPs and Nbs for a set of representative targets, including Nb_LAM2_ for mCherry, Nb_59H10_ for HIV p24 protein, Nb_2E7_ for HIV gp41 protein, Nb_ALFA_ for ALFA-tag peptide, Nb_LAG16_ for GFP, and Nb_BC2_ for β-catenin. This two-step approach, first optimizing Nb_GFP_ and subsequently applying these adjustments to enhance the fluorescence performance across multiple Nbs, does not require optimization for each new chimera. Thus, our approach establishes a versatile and expandable platform for efficiently creating various antigen-specific VIS-Fbs, advancing protein labeling and imaging. As we aimed to establish the feasibility and broad applicability of the VIS-Fb synthetic platform across diverse protein pairs, the VIS-Fbs were not individually optimized. However, quantitative comparisons of properties such as turn-on efficiency and absolute brightness may be necessary in future studies aimed at specific applications.

Even though all Nbs have structural similarities, they bind cognate antigens using different regions. Thus, all three CDRs and FR3 of LaG16 (PDB ID 6LR7) form specific contacts with GFP^68^. All three CDRs of LAM2 (PDB ID 7SAJ) bind mCherry^53^. To interact with HIV protein p24, Nb_59H10_ (PDB ID 5O2U) uses CDRs1-3 and FR2^45^, while binding of Nb2E7 (PDB ID 5HM1) to HIV protein gp41 occurs via CDR1, CDR2, and FR2^46^. Connection of Nb_ALFA_ (PDB ID 6I2G) with ALFA-tag peptide is mainly built from CDR2 and CDR3, also involving residues within FR2^48^. BC2 Nb (PDB ID 5IVN) engages β-catenin epitope mainly via CDR3 and FRs 2 and 3^69^. Compared to all other used Nbs, Nb_GFP_ (PDB ID 3OGO) is a unique Nb that has a notably shorter CDR3: 9 a.a. *vs.* 13-20 a.a. residues for other Nbs^16, 70^. Unlike all other Nbs listed, which bind external surfaces or peptide epitopes, short and compact CDR3 of Nb_GFP_ penetrates GFP barrel and stabilizes the chromophore. We hypothesize that this compactness of Nb_GFP_ CDR3 allows it to tolerate insertion of truncated GFP-like FPs, while insertion of truncated FPs in Nbs with long CDR3, which are more critical for binding and stability, may disrupt folding or binding of respective VIS-Fbs.

Our data from different GCaMP calcium indicator-expressing transgenic mice demonstrate that the AAV-mediated dTomato-Fb_LAG16_ expression is well tolerated across various cell types, including excitatory and inhibitory neurons and astrocytes (**Fig. 5** and **Supplementary Figs. 24-25**). Moreover, VIS-Fb binding to GCaMP did not significantly affect its activity. Compared to the direct fusion of FP to the targeted protein, VIS-Fb does not directly affect target protein’s folding and activity. The antigen-dependent stabilization and fluorescence of VIS-Fbs were sufficiently high to enable high-fidelity low-background ratiometric measurements over minutes to hours in living mice. The expression of VIS-Fbs also facilitated computational analyses, such as measuring somatic calcium activity while minimizing neuropil contamination. VIS-Fbs can be easily adapted to different fluorescent colors and binding targets without re-engineering the primary sensor construct. This facilitates rapid development of ratiometric sensor pairs and multiplexed imaging strategies. VIS-Fbs are small enough to fit into AAVs. VIS-Fbs can be expressed under separate promoters, allowing cell-type–specific or inducible labeling without altering the expression of the target protein. Using our intersectional approach, VIS-Fbs enabled tagging and measurement from specific cell compartments and subpopulations in living mice.

The sfGFP-Fb_BC2_ nanobody enabled specific, dynamic, and non-invasive *in vivo* imaging of endogenous β-catenin during zebrafish embryogenesis, including conditions of pharmacological activation and inhibition of the Wnt/β-catenin pathway (**Fig. 6** and **Supplementary Figs. 27–29**). This approach revealed spatiotemporal dynamics of endogenous β-catenin accumulation and degradation *in vivo*, including early developmental stages prior to gastrulation, which could not previously be directly visualized in live animals with high spatiotemporal precision and low background. By capturing real-time changes in distribution and signal intensity of endogenous β-catenin at single-cell resolution, sfGFP-Fb_BC2_ uncovered specific responses and morphological outcomes following modulation of Wnt signaling, patterns that were not accessible using existing transcriptional reporters or immunostaining. These findings demonstrate the ability of VIS-Fb to study the regulation of endogenous proteins in live vertebrates, offering new insight into the spatial and temporal coordination of signaling during development.

We anticipate that the synthetic biology-based VIS-Fb toolbox will find applications in various areas of cell and developmental biology, including live-cell multicolor imaging and protein tracking, studying protein and cellular dynamics, pulse-chase experiments for labeling newly synthesized proteins, tracing different cell lineages, super-resolution microscopy approaches, measuring metabolites in proximity to targeted antigens, and multiplexed labeling and functional imaging *in vivo*^71^.

## Methods

### Construction of mammalian plasmids

To generate mCherry inserted into Nb_GFP_ with Gly_2_Ser linkers, without linkers or with deletions from N- and/or C-termini, miRFP670nano3 gene was swapped with mCherry gene (mCherry-pBAD; Addgene no.54630) in the NIR-Fb plasmid (Addgene no.184675) by NEBuilder HiFi DNA Assembly Kit.

VIS-Fbs to GFP with insertions of mTagBFP2, mTFP1, mWasabi, mNeonGreen, mOrange, LSSmOrange, CyOFP1, mScarlet-I, mScarlet, LSSmScarlet, mNeptune2, mCardinal, PAmCherry or mEos4a were designed by swapping miRFP670nano3 gene with FPs genes in the NIR-Fb plasmid (Addgene no.184675) by NEBuilder HiFi DNA Assembly Kit. All oligonucleotide primers for PCR amplification were purchased from Thermo Fisher Scientific (**Supplementary Table 1**). mTagBFP2, mTFP1, mWasabi, mNeonGreen, mOrange, LSSmOrange, CyOFP1, mScarlet-I, mScarlet, LSSmScarlet, mNeptune2, mCardinal, PAmCherry, and mEos4a were amplified from pBAD-mTagBFP2 (Addgene no.34632), pBAD-mTFP1 (Addgene no.54553), pmWasabi-FAK-5 (Addgene no.56502), pmOrange-N1 (Addgene no.54499), pBAD-LSSmOrange (Addgene no.37129), pKK-CyOFP1-TEV (Addgene no.105782), pmScarlet-I-C1 (Addgene no.85044), pmScarlet-C1 (Addgene no.85042), pcDNA3-mNeptune2 (Addgene no.51309), pcDNA3-mCardinal (Addgene no.51311), PAmCherry1-C1 (Addgene no.54495) and mEoa4a-N1 (Addgene no.54811) plasmids, respectively.

To generate the NES-mTagBFP2-Fb plasmid, the mTagBFP2-Fb_GFP_ gene was swapped with DEVD-mCardinal-3xNLS construct in the pcDNA-NES-DEVD-mCardinal-3xNLS plasmid (Addgene no.164052) at the BamHI/XhoI sites. To construct an H2B-CyOFP1-Fb_GFP_ plasmid, the CyOFP1-Fb gene was amplified and inserted into the pcDNA plasmid with an H2B signal (Addgene no.20972) at the BamHI/XbaI sites. To generate the Mito-mCardinal-Fb_GFP_ plasmid, the mCardinal-Fb gene was amplified and swapped with LSSmKate2 in the pMito-LSSmKate2-N1 (Addgene no.31879) by NEBuilder HiFi DNA Assembly Kit.

VIS-Fb to p24 with inserted mCardinal, VIS-Fb to gp41 with inserted mTagBFP2 and VIS-Fbs to ALFA-tag with insertions of mTagBFP2, TagRFP-T, CyOFP1, mScarlet, mStayGold, and mCardinal were cloned into the pcDNA plasmid by NEBuilder HiFi DNA Assembly Kit by swapping miRFP670nano3 gene by respective FP genes in the following plasmids: NIR-Fb_59H10_ (Addgene no. 184682), NIR-Fb_2E7_ (Addgene no.184681) and NIR-Fb_ALFA_ (Addgene no.184678). TagRFP-T and mStayGold with Gly_2_Ser linkers were PCR amplified from mTagRFP-T-Mito-7 (Addgene no.58023) and pESETB/mStayGold (Addgene no.212017).

HIV p24-sfGFP, gp41-sfGFP, and ALFA-tag-sfGFP from ^8^ were amplified by QuikChange PCR protocol to insert the sfGFPY66G point mutation. Note that gp41-sfGFP has a truncated N-terminus (ΔMGAASLTLTVQARQLLS). For membrane localization, p24-sfGFPY66G was cloned to pKA-142 plasmid (Addgene no.79835) by NEBuilder HiFi DNA Assembly Kit. For cytoplasmic localization, gp41-sfGFPY66G was cloned to pcDNA-NES-DEVD-mCardinal-3xNLS plasmid (Addgene no.164052) by NEBuilder HiFi DNA Assembly Kit. For nuclear localization, ALFA-tag-sfGFPY66G was cloned to the pcDNA-H2B-mCherry plasmid (Addgene no. 20972) by NEBuilder HiFi DNA Assembly Kit.

VIS-Fb to mCherry with insertions of mTagBFP2, mWasabi, and mClover3 were generated by overlap PCR, and the resulting constructs were cloned into the pcDNA plasmid by KpnI/EcoRI restriction sites. Anti-mCherry Nb was amplified from the pGEX6P1-mCherry-Nanobody (LaM-2) (Addgene no.162276), mClover3 was amplified from pNCS-mClover3 (Addgene no.74236).

jRGECO1a biosensor was amplified from a pGP-CMV-NES-jRGECO1a (Addgene no.61563) plasmid and inserted into the NIR-Fb_LAG30_ (Addgene no.220740) plasmid by NEBuilder HiFi DNA Assembly Kit instead of miRFP670nano3.

To generate dTomato-Fb_LAG16_, the dTomato gene was PCR amplified from the pCAG-Kir2.1-T2A-tdTomato (Addgene no.60598) plasmid with Gly_2_Ser linkers and inserted into the NIR-Fb_LAG16_ (Addgene no.220739) plasmid instead of miRFP670nano3.

Soma-targeted dTomato-Fb_LAG16_ was engineered by PCR amplifying dTomato-Fb_LAG16_ and swapping it with the jGCaMP8s gene in the AAV-hSyn-RiboL1-jGCaMP8s (Addgene no.169247) plasmid by NEBuilder HiFi DNA Assembly Kit. To construct dTomato-Fb_LAG16_ expressed in the astrocytes or GABAergic interneurons, the dTomato-Fb_LAG16_ gene was amplified and inserted into the pAAV-3xcore2_eHGT_390m-minBglobin-SYFP2-4x2C-WPRE3-BGHpA (Addgene no.208167) or pAAV-DLX2.0-minBG-SYFP2-WPRE3-BGHpA (Addgene no.163505) plasmids, respectively.

To generate VIS-Fb to β-catenin with inserted sfGFP, sfGFP was cloned into the pcDNA plasmid by NEBuilder HiFi DNA Assembly Kit by swapping the miRFP670nano3 gene with sfGFP in the NIR-Fb_BC2_ plasmid^8^. sfGFP with Gly_2_Ser linkers was PCR amplified from p24-sfGFP.

### Construction of zebrafish plasmids

To engineer sfGFP-Fb_BC2_ under Tol2 transposon-based vector, sfGFP-Fb_BC2_ was cloned into pTol2-Ef1a-BlobinIntron-EGFP-LC3-RFP-LC3ΔG by NEBuilder HiFi DNA Assembly Kit by swapping the EGFP-LC3-RFP-LC3ΔG gene with sfGFP-Fb_BC2_.

### Mammalian cell culture and transfection

HeLa (CCL-2) were obtained from the ATCC and were not additionally authenticated or tested for mycoplasma contamination. Cells were cultured in a DMEM medium (Corning) supplemented with 10% FBS (HyClone), 0.5% penicillin-streptomycin (HyClone) and 2 mM glutamine at 37°C. For live-cell fluorescence microscopy, cells were plated in 35 mm glass-bottom Petri dishes (MatTek). Transient transfections were performed using polyethylenimine^72^ or Effectene reagent (Qiagen).

Green Pegassos (Addgene no.163114) and PyronicSF (Addgene no. 124812) plasmids were used for co-transfection with the jRGECO1a-Fb_LAG30_ construct.

For multicolor imaging of VIS-Fbs, HeLa cells were co-transfected with plasmids encoding NES-mTagBFP-Fb_GFP_, Mito-Cardinal-Fb_GFP_, H2B-CyOFP1-Fb_GFP_ and a cognate mEGFP antigen in a 1.6:1:1.6:1.6 ratio.

### Neuronal culture and transfection

Primary mouse hippocampal neuronal cultures were prepared from the P0-P1 C67BL/6J mice according to the protocol^73^. All animal work was performed under the NIH guidelines and was approved by the Albert Einstein College of Medicine Institutional Animal Care and Use Committee guidelines. Cells were plated at a density of ~60,000 cells per dish onto poly-D-lysine (Sigma) coated glass-bottomed dishes (MatTek). Neurons were cultured at 37°C and 5% CO2 in the Neurobasal A medium (Gibco) supplemented with 1% NeuroB27 (Millipore), 1 mM GlutaMAX (Gibco) and 0.5% penicillin-streptomycin (HyClone). Ten percent of the medium was exchanged twice per week. Transfection was performed on the day *in vitro* 12 or 14 (DIV12 or DIV14) using the Calcium Phosphate Transfection kit (Invitrogen) according to the previously published protocol^74^.

### Epifluorescence live-cell microscopy

Live mammalian cells were imaged using an Olympus IX81 inverted epifluorescence microscope equipped with a Lambda LS Xenon light source (Sutter). An ORCA-Flash4.0 V3 camera (Hamamatsu) was used for image acquisition. Cells were imaged using a 60× 1.35 NA oil immersion objective lens (UPlanSApo, Olympus) and 390/40 nm, 436/20 nm, 475/42 nm, 480/40 nm, 523/20 nm, 543/22 nm, 575/25 nm, or 605/30 nm excitation filters and 460/40 nm, 480/40 nm, 535/40 nm, 575/30, 580/30 nm, 588/21 nm, 615/30 nm, or 667/30 nm emission filters, respectively. Light power densities measured at the rear aperture of the objective lens were 0.76 mW cm^−2^ for 390/40 nm, 0.11 mW cm^−2^ for 436/20 nm, 4.14 mW cm^−2^ for 475/42 nm, 3.73 mW cm^−2^ for 480/40 nm, 0.09 mW cm^−2^ 523/20 nm, 2.21 mW cm^−2^ for 543/22 nm, 1.42 mW cm^−2^ for 575/25 nm, and 3.41 mW cm^−2^ for 605/40 nm excitation filters, which gives an estimate (FN is 26.5 for the objective lens) for the light power densities at the front focal objective plane of 0.47 W cm^−2^, 0.07 W cm^−2^, 2.57 W cm^−2^, 2.32 W cm^−2^, 0.06 W cm^−2^, 1.37 W cm^−2^, 0.88 W cm^−2^, and 2.12 W cm^−2^, respectively.

While imaging, HeLa cells were incubated in a DMEM medium and kept at 37°C. The microscope was operated with SlideBook2024 software (Intelligent Imaging Innovations). For imaging performance of jRGECO1a calcium biosensor inserted into a LAG30 Nb, HeLa cells were washed twice with pre-warmed 20 mM HEPES-buffered Hanks’ Balanced Salt Solution (HHBSS) (Corning) before imaging and incubated in HHBSS supplemented with 1 mM CaCl_2_. After the first frame was imaged, 5 μM ionomycin (Cayman Chemical) was added. Cells were maintained at 37 °C using the environmental chamber. When jRGECO1a-Fb_LAG30_ was co-expressed with Green Pegassos and PyronicSF, HeLa cells were incubated in HHBSS 30 min before imaging. After the first frame was imaged, a 1 mM or 10 mM solution of sodium pyruvate (Gibco) was added, respectively; 5 μM ionomycin (Cayman Chemical) was added to the same cells after the 5^th^ or the 10^th^ imaging frame. For imaging a glucose biosensor, iGlucoSnFr, co-expressed with jRGECO1a-Fb_LAG30_, HeLa cells were incubated in a DMEM medium without glucose and pyruvate (Gibco). After the first frame was imaged, a 50 mM glucose solution was added; 5 μM ionomycin (Cayman Chemical) was added to the same cells after the 5^th^ imaging frame.

For imaging mTagBFP2-Fb_GFP_, mTagBFP2-Fb_2E7_, mTagBFP2-Fb_ALFA_, mTagBFP2-Fb_LAM2_ mTagBFP2, and EBFP2, 390/40 nm excitation and 460/40 nm emission filters (Chroma) were used. For imaging mWasabi-Fb_GFP_, mNeonGreen-Fb_GFP_, green form of mEos4a-Fb_GFP_, mStayGold-Fb_ALFA_, mWasabi-Fb_LAM2_, mClover3-Fb_LAM2_, sfGFP-Fb_BC2_, mEGFP, mVenus, p24-sfGFP, gp41-sfGFP, GCaMP6s, Green Pegassos, PyronicSF, and iGlucoSnFr 480/40 nm excitation and 535/40 nm emission filters (Chroma) were used. For imaging mCherry-Fb_GFP_, mScarlet-Fb_GFP_, mScarlet-I-Fb_GFP_, PAmCherry-Fb_GFP_, red form of mEos4a-Fb_GFP_, TagRFP-T-Fb_ALFA_, mScarlet-Fb_ALFA_, jRGECO1a-Fb_LAG30_, dTomato-Fb_LAG16_, mCherry, and jRGECO1a 575/25 nm excitation and 615/30 nm emission filters (Chroma) were used. For imaging mTFP1-Fb_GFP_, 436/20 nm excitation and 480/40 emission filters (Chroma) were used. For imaging mOrange-Fb_GFP_, 543/22 nm excitation and 580/30 nm emission filters (Chroma) were used. For imaging LSSmOrange-Fb_GFP_, CyOFP1-Fb_GFP_, CyOFP1-Fb_ALFA_, and LSSmScarlet-Fb_GFP_, 436/20 nm, 523/20 nm and 475/42 nm excitation and 575/30 nm, 588/21 nm and 615/30 nm emission filters (Chroma) were used, respectively. For imaging mNeptune2-Fb_GFP_, mCardinal-Fb_GFP_, mCardinal-Fb_59H10_, 605/30 nm excitation and 667/30 nm emission filters (Chroma) were used. The data were analyzed using SlideBook2024 (Intelligent Imaging Innovations), GraphPad Prism v.10, and Origin 2017 SR1 software.

### Flow cytometry

Flow cytometry analysis was performed using an LSR II or FACSymphony A5 SE (BD Biosciences) flow cytometer. Before analysis, live cells were washed with cold PBS, trypsinized for 5 min at 37°C, and diluted in cold PBS containing 4% FBS and 2 mM EDTA to a density of 500,000 cells per ml.

About 50,000-100,000 cells per sample were recorded. Data were collected using BD FACSDiva v.8.0.1 (BD Biosciences) software. The fluorescence intensity of cells expressing mTagBFP2, EBFP2, and Nbs with inserted mTagBFP2, mTFP1 or LSSmOrange was analyzed using the 405 nm excitation laser and 450/50 nm (mTagBFP2, EBFP2) or 510/40 nm (mTFP1), 595/30 nm (LSSmOrange) emission filters, respectively. The fluorescence intensity of cells expressing mEGFP, mVenus, p24-sfGFP, gp41-sfGFP, ALFA-tag-sfGFP, and Nbs with inserted mWasabi, mNeonGreen, mClover3, mStayGold, sfGFP, CyOFP1 or LSSmScarlet was analyzed using the 488 nm excitation laser and 525/50 nm (mEGFP, mWasabi, mNeonGreen, mClover3), 510/20 nm (sfGFP) 537/32 nm (mVenus), 582/15 nm (CyOFP1) or 610/20 nm (LSSmScarlet) emission filters, respectively. The fluorescence intensity of mCherry and Nbs with inserted mOrange, mCherry, mScarlet, mScarlet-I, TagRFP-T, dTomato, mNeptune2, mCardinal was analyzed using a 561 nm laser for excitation, and its fluorescence was detected with a 582/15 nm (mOrange, mScarlet, and mScarlet-I), 610/20 nm (mCherry, TagRFP-T, and dTomato), 670/30 nm (mNeptune2 and mCardinal) emission filters, respectively. The fluorescence intensity of cells expressing Nbs with inserted miRFP670nano3 was analyzed using the 640 nm excitation laser and 660/20 nm emission filter. The data were analyzed using FlowJo v.7.6.2 software. To estimate the brightness of VIS-Fbs, flow cytometry gating was performed for intact cells, single cells, and live cells. These cells then were gated for non-transfected cells in green (for VIS-Fb_GFP_s, mCatdinal-Fb_59H10_, mTagBFP2-Fb_2E7_, VIS-Fb_ALFA_s, and dTomato-Fb_LAG16_; Ex. 488 nm, Em. 525/50 nm or Em. 510/20 nm), blue (VIS-Fb_GFP_s; Ex. 405 nm, Em. 450/50) or red (VIS-Fb_LAM2_s; Ex. 561 nm, Em. 610/20 nm or Em. 602/40 nm) channels for selecting antigen-positive cells. The latter cells were used to assess the brightness in the channel of an antigen and the channel of VIS-Fb^8^. The maximal fluorescence of antigen-bound form for each VIS-Fb was normalized to 100%.

### Animal subjects

The mouse strains used in this study included GFAP-Cre (RRID: IMSR_JAX:012886), Viaat-Cre (RRID: IMSR_JAX:017535), Ai95(RCL-GCaMP6f)-D (RRID: IMSR_JAX:024105), and Thy1-GCaMP6f mice (RRID: IMSR_JAX:025393). All mice were on a C57BL/6J background. For the *in vivo* imaging and immunostaining experiments, we utilized four male GFAP-GCaMP6f mice, two male Viaat-GCaMP6f mice, and four female Thy1-GCaMP6f mice, all of which were between 13-17 weeks old at the time of imaging (~4.5–5.5 weeks after stereotactic injections). The experimental mice used in individual experiments typically originated from different litters. Each mouse had unique identification marks. No specific criteria were applied for assigning mice to experimental groups. The mice were group-housed, provided with bedding and nesting materials, and maintained on a 12-h light-dark cycle in a temperature (22 ± 1°C) and humidity-controlled (45–65%) environment.

### Zebrafish husbandry

All protocols using zebrafish in this study were approved by the Albert Einstein College of Medicine Institutional Animal Care and Use Committees. Adult zebrafish and embryos up to 3 days postfertilization (dpf) were maintained as described previously^75^. For all experiments, larvae were anesthetized in E3 media without methylene blue supplemented with 0.16 mg/mL Tricaine (MS222/ethyl 3-aminobenzoate; Sigma-Aldrich).

### Virus production and stereotactic injection

Recombinant AAV9 production was carried out by the Salk Institute’s Viral Vector Core. The recombinant AAV9-3xCore2(390m)-dTomato-Fb_LAG16_, AAV9-DLX2.0-dTomato-Fb_LAG16_, and AAV9-hSyn-RiboL1-dTomato-Fb_LAG16_ viruses had a titer of 7.24E+13 GC ml-1, 1.16E+14 GC ml-1, and 9.3E+13 GC ml-1, respectively. 0.4 μl of 1:10 diluted AAV was injected into the somatosensory cortex (coordinates: AP −0.5 - (−0.8) mm; ML 1.45-1.55 mm; DV 0.28-0.30 mm and AP −1.5 - (−1.8) mm; ML 1.4-1.5 mm; DV 0.26-0.31 mm). Surgical procedures closely followed previously established protocols. Briefly, thin-wall glass pipettes were pulled on a Sutter Flaming/Brown micropipette puller (model P-97). Using a sterile technique, pipette tips were cut at an acute angle under 10× magnification. Tip diameters were typically 15–20 μm. Pipettes that did not result in sharp bevels nor had larger tip diameters were discarded. Millimeter tick marks were made on each pulled needle to measure the virus volume injected into the brain.

Mice were anesthetized with isoflurane (4–5% for induction; 1–1.5% for maintenance) and positioned in a computer-assisted stereotactic system with digital coordinate readout and atlas targeting (Leica Angle Two). Body temperature was maintained at 36–37 °C with a direct current (DC) temperature controller, and ophthalmic ointment was used to prevent the eyes from drying. A small amount of depilator cream (Nair) was used to remove hair over the dorsal areas of the injection site. The skin was cleaned and sterilized with a two-stage scrub of betadine and 70% ethanol, repeated three times.

A midline incision was made, beginning just posterior to the eyes and ending just past the lambda suture. The scalp was pulled open, and the periosteum was cleaned using a scalpel and forceps to expose the desired hemisphere for calibrating the digital atlas and coordinate marking. Once reference points (bregma and lambda) were positioned using the pipet needle and entered into the program, the desired target was set on the digital atlas. The injection pipet was carefully moved to the target site (using AP and ML coordinates). Next, the craniotomy site was marked, and an electrical micro-drill with a fluted bit (0.5 mm tip diameter) was used to thin a 0.5–1 mm diameter part of the bone over the target injection site. Once the bone was thin enough to flex gently, a 30 G needle with an attached syringe was used to carefully cut and lift a small (0.3–0.4 mm) segment of bone.

For injection, a drop of AAV was carefully pipetted onto parafilm (2–3 μl) to fill the pulled injection needle with the desired volume. Once loaded with sufficient volume, the injection needle was slowly lowered into the brain until the target depth was reached. Manual pressure was applied using a 30-ml syringe connected by shrink tubing, and 0.4 μl of the virus was slowly injected over 5–10 min. Once the virus was injected, the syringe’s pressure valve was locked. The position was maintained for approximately 10 min to allow the virus to spread and avoid backflow upon needle retraction. Each animal received two injections (~1 mm apart) in one hemisphere. Following the injections, head clamps were removed, muscles were approximated, and the skin was sutured along the incision. Mice were given subcutaneous Buprenorphine SR (0.5 mg per kg) and allowed to recover before placement in their cage.

### Animal preparation for in vivo two-photon imaging

Surgical procedures closely followed established protocols^8, 76, 77^. Briefly, mice were anesthetized with isoflurane (4–5% for induction; 1–1.5% for maintenance) on a custom surgical bed (Thorlabs). Body temperature was maintained at 36–37 °C with a DC temperature control system, and ophthalmic ointment was used to prevent the eyes from drying. Depilator cream (Nair) was used to remove hair above the imaging site. The skin was thoroughly cleansed and disinfected with a two-stage scrub of betadine and 70% ethanol, repeated three times.

A scalp portion was surgically removed to expose frontal, parietal, and interparietal skull segments. Scalp edges were attached to the lateral sides of the skull using a tissue-compatible adhesive (3M Vetbond). A custom-machined metal plate was affixed to the skull with dental cement (Coltene Whaledent, cat. no. H00335), allowing the head to be stabilized with a custom holder. Animals intended for awake imaging were allowed to recover for at least three days and then habituated to head fixation on a spherical treadmill (typically three sessions over three consecutive days). On the day of imaging, an approximately 2 mm W x 3 mm L diameter craniotomy was made over the AAV injection sites. A ~1.5% agarose solution and coverslip were applied to the exposed tissue. The coverslip was affixed to the skull with dental cement to control tissue motion. Mice imaged under awake conditions were allowed to recover from isoflurane anesthesia for at least 1–1.5 h before recordings commenced.

### In vivo two-photon microscopy

Live animal imaging was performed as previously described^5^. Briefly, a Sutter Movable Objective Microscope equipped with a pulsed femtosecond Ti:Sapphire laser (Chameleon Ultra II, Coherent) and two fluorescence detection channels were used for imaging (green emission filter: ET525/70M (Chroma); red emission filter: ET605/70M (Chroma) supplemented with ET645/75M (Chroma) and FF01-720/SP (Semrock); dichroic beamsplitter: 565DCXR (Chroma); photomultiplier tubes: H7422-40 GaAsP (Hamamatsu)). The laser excitation wavelength was set to either 920 or 1,000 nm. The average laser power was <10–15 mW at the tissue surface and adjusted with depth as needed to compensate for signal loss due to scattering and absorption. At the excitation intensities and durations used in this study, no signs of phototoxicity, such as a gradual increase in baseline fluorescence, lasting changes in activity rate, or blebbing of labeled cells, were apparent in our recordings. An Olympus 20× 1.0 NA water immersion objective was used for light delivery and collection. Z-stacks included up to 750 images, acquired at 1 μm axial step size, using a two-frame average, 512 × 512-pixel resolution, and 1.3× - 2.0× zoom (corresponding to 562 μm - 350 μm fields of view). Time-lapse recordings used a 256 × 256-pixel resolution, 1.0× zoom (corresponding to a 348 μm field of view), and 8.14 Hz frame rate. Awake recordings were performed in head-fixed mice on a spherical treadmill equipped with an optical encoder (E7PD-720-118; US Digital). Analog and imaging data were acquired synchronously, allowing the animal’s running speed to be related to cellular fluorescence changes.

### Brain tissue fixation, slicing, and immunostaining

Following American Veterinary Medical Association (AVMA) guidelines, mice were euthanized in their home cage. Transcardial perfusion was performed using 10% sucrose followed by 4% paraformaldehyde (PFA). Brain tissue was carefully extracted and incubated in 4% PFA overnight at 4°C. The tissue was then washed on a shaker with 1× PBS three times (15 min per wash). A Leica VT1000S vibratome was used to prepare 40-μm-thick coronal tissue sections. Immunostaining was performed on floating sections using standard techniques. Primary and secondary antibodies included GFP (1:200 dilution; Thermo Fisher Scientific; RRID: AB_2534023), NeuN (1:250 dilution; Novus Biologicals; RRID: AB_11036146), Sox9 (1:250 dilution; Abcam; RRID: AB_2728660) and Alexa Fluor 488 goat anti-chicken (1:100 dilution; Thermo Fisher Scientific; RRID: AB_2534096), Alexa Fluor Plus 405 goat anti-mouse (1:100 dilution; Thermo Fisher Scientific; RRID: AB_2890536), Alexa Fluor 633 goat anti-rabbit (1:100 dilution; Thermo Fisher Scientific; RRID: AB_2535732), respectively.

### Confocal microscopy of brain slices

Confocal imaging of brain sections (three per animal) was performed on a Zeiss LSM 710 with ZEN Black software (v2011) and an Olympus 20× 0.8 NA air-matched objective. Four-channel, 4 × 6 tiled z-stacks (~15 images at 1 μm axial step size) were acquired using 405 nm, 488 nm, 561 nm, and 633 nm laser lines. Each z-stack image had a 1024 × 1024-pixel resolution.

### Image and analog data processing

Data were processed, analyzed, and plotted in Fiji (2.0.0-rc- 61/1.54g; SciJava), Imaris (v10.0; Oxford Instruments), Igor Pro (v9.05; WaveMetrics), and Prism software (v10.0; GraphPad).

Immunostained tissue from AAV9-3xCore2(390m)-dTomato-Fb_LAG16_-injected GFAP-GCaMP6f mice was quantified by creating digital representations of labeled cells using Imaris’ surface creation wizard. We used a surface grain size of 0.5 µm for both GCaMP6f and dTomato. The seed points diameter was set to 8 µm, and the seed points quality filter included the top 15% of GCaMP6f+ and dTomato+ structures. Immunostained tissue from AAV9-DLX2.0-dTomato-Fb_LAG16_ and AAV9-hSyn-RiboL1-dTomato-Fb_LAG16_-injected Viaat-GCaMP6f and Thy1-GCaMP6f mice, respectively, was quantified by creating digital representations of labeled cell bodies using Imaris’ spot creation wizard. We applied an ‘XY Diameter’ spot size filter of 10 µm for GCaMP6f, dTomato, and NeuN and 9 µm for Sox9 to background-subtracted images. The ‘Quality’ filter was set to include the top 5% of Viatt-GCaMP6f+ and Thy1-GCaMP6f+ cells, the top 2% of DLX2.0-dTomato+ and hSyn-RiboL1-dTomato+ cells, the top 6% of NeuN+ cells, and the top 6% of Sox9+ cells, accounting for differences in labeling intensity and uniformity. Created spots and surfaces were then used to determine the percent colocalization with Imaris’ MATLAB colocalization function. Spots were considered colocalized with one another if the distance between them was ≤1 μm. Two surfaces were deemed colocalized if the distance between them was ≤1 μm. Spots were considered colocalized with a surface if the distance between them was ≤0.5 μm. Analysis was restricted to the top ~600 μm of the cortex because the typical AAV injection depth was ~300 μm.

Analog data from the spherical treadmill encoder were processed using Igor Pro.

### Zebrafish embryo injections and drug treatment

*Danio rerio* embryos derived from AB animals were injected with 1 nL of microinjection mix with 25 ng/μl of pTol2-sfGFP-Fb_BC2_ driven by the Ef1a promoter and 20 ng/μl of Transposase mRNA. Embryos were scored at 5-8 h and 24 post-fertilization (hpf) for GFP expression using a Leica M205 FCA THUNDER Imager Model Organism Large fluorescent dissecting scope.

To modulate Wnt/β-catenin signaling *in vivo*, zebrafish embryos were injected at the one-cell stage with pTol2-sfGFP-Fb_BC2_ driven by the Ef1a promoter. Embryos were maintained at 28°C in E3 medium and screened for mosaic green fluorescence at 5-8 h and 24 h post-injection (hpi).

For late-stage modulation, fluorescent embryos were randomly allocated to treatment or control groups at 24 hpi. The treatment group was incubated in 10 µM IWR-1 (Sigma-Aldrich) for 6 h, whereas control embryos remained untreated. Long-term imaging was performed at 30 hpi. For early-stage modulation, injected embryos at the 30–50 % epiboly stage (~6 hpi, before the appearance of detectable fluorescence) were divided into three groups: (i) untreated control, (ii) 10 µM IWR-1, or (iii) 0.2 mM LiCl (Thermo Fisher Scientific). Embryos were treated in 24-well plates containing fresh E3 medium and incubated for 24 h at 28.5 °C. Following treatments, embryos were examined by fluorescence stereomicroscopy or long-term intravital confocal microscopy to assess β-catenin reporter distribution and associated morphological phenotypes. Each experiment included at least two independent biological replicates with ≥ 150–200 injected embryos per replicate.

### Zebrafish imaging

All long-term imaging was performed using a zWEDGI device as previously described^56, 57^. Briefly, embryos were dechorionated and anesthetized in E3 supplemented with 0.16 mg/ml Tricaine. Embryos were loaded into a zWEDGI chamber for time-lapse imaging. The loading chamber was filled with 1% low-melting-point agarose (Sigma-Aldrich) in E3 to retain embryos in the proper position for imaging. Additional E3 supplemented with 0.16 mg/ml Tricaine was added as needed to avoid dryness and provide the required moisture to zebrafish during imaging acquisition. Time-lapse videos were acquired on a Spinning disk confocal microscope Nikon (CSU-W, Yokogawa) with a confocal scanhead on a Nikon Ti2 inverted microscope equipped with Photometrics Evolve EMCCD camera, and an EC Plan NeoFluar NA 0.3/10 x air objective, z-stacks, 5 μm optical sections, every 10 minutes, and 2048 x 2048 resolution. For image acquisition and assessment of the IWR-1 and LiCl 24-hour drug treatment effect at early developmental stages, images were taken at 30 hpf on a Leica M205 FCA THUNDER Imager Model Organism Large and an objective PlanApo 1.0x M-series. Embryos were dechorionated and then immersed in E3 supplemented with 0.16 mg/mL tricaine during imaging. Videos were processed on Imaris software version 10.2.

## Ethical statement

All live animal procedures were performed following the National Institutes of Health (NIH) guidelines and were approved by the Institutional Animal Care and Use Committee (IACUC) of the Salk Institute under protocol number 13-00022 and the Albert Einstein College of Medicine Institutional Animal Care and Use Committee guidelines. Mice were group-housed in a standard 12 h light/12 h dark cycle. Dissociated mouse hippocampal neuron cultures were prepared from male and female mice (P0-P1): C57BL/6J (Charles River).

## Statistics and reproducibility

For *in vivo* two-photon imaging experiments, the sample size after data exclusion (see below) was 2–4 biological (mice) and two technical replicates (injection sites). For immunostaining experiments, the sample size after data exclusion was 2–4 biological (mice) and three technical replicates (tissue sections). Sample sizes were not predetermined but aligned with numbers typically used for *in vivo* characterizations.

Mice with suboptimal surgical preparation (e.g., visible surface damage) were excluded from analysis because surgery quality can affect the imaging (e.g., signal-to-noise ratio, resolution) and immunostaining results (e.g., glial reactivity). Immunostained slices with weak or irregular staining were excluded from the data analysis.

The results described were reproducible within and across animals. Sex as a biological variable was not considered in the research design and analyses. The study’s primary goal was to demonstrate the approach’s technical capabilities.

The two-photon and confocal microscopy experiments did not involve treatment groups or identifiable covariates. Thus, randomization was not used, and covariates were not controlled in statistical analyses. The investigators were not blinded to allocation during the two-photon and confocal microscopy experiments and outcome assessment. Blinding was not necessary because no treatment groups existed. Immunostaining analyses were carried out independently from the *in vivo* image data collection. Data analysis was performed using automated software routines.

## Data availability

All data supporting the findings of this study are available within the article, its Supplementary Information, and the Source Data file. Major plasmids constructed in this study, with their sequences, are deposited at Addgene. Protein structures used in this study include PDB IDs: 3OGO [https://www.rcsb.org/structure/3ogo], 1ZGO [https://www.rcsb.org/structure/1ZGO], 3WLD [https://www.rcsb.org/structure/3WLD], 6LR7 [https://www.rcsb.org/structure/6lr7], 3M24 [https://www.rcsb.org/structure/3M24], and 7SAJ [https://www.rcsb.org/structure/7SAJ].

## Acknowledgments

We thank Dong Woo Hwang (Albert Einstein College of Medicine) for the hippocampal neuronal cultures, Fedor Subach (National Research Center “Kurchatov Institute”) for the pBAD/HisB-LSSmScarlet and pBAD/HisB-mNeonGreen plasmids, Koichi Kawakami (National Institute of Genetics) for the pTol2-Ef1a-BlobinIntron-EGFP-LC3-RFP-LC3ΔG plasmid, Edward Callaway (Salk Institute for Biological Studies) for the targeting approach discussions, the Flow Cytometry Core Facility (Albert Einstein College of Medicine) for the assistance with cell sorting, the Viral Vector Core Facility (Salk Institute for Biological Studies) for the virus production, Clinton W. DePaolo and Spartak Kalinin at the Zebrafish Core Facility (Albert Einstein College of Medicine) for the technical support, and the Analytical Imaging Facility (Albert Einstein College of Medicine). This work was supported by grants GM122567 (V.V.V.), NS123719 (A.N.) and GM118027 (SdO) from the US National Institutes of Health, 220011 from the Jane and Aatos Erkko Foundation, 360277 from the Research Council of Finland, MET-000000045 from the Chan Zuckerberg Initiative Foundation (all to V.V.V.), the NOMIS Foundation Neuroimmunology Initiative, and the Edwards-Yeckel Research Foundation (all to A.N.). Figure 6a and d were created with BioRender.com.

## Author contributions

N.V.B. developed VIS-Fbs, characterized and imaged them in mammalian cells. E.C. performed AAV injections and immunohistochemistry. A.N. planned and conducted the *in vivo* two-photon imaging. O.S.O. developed the prototypes of mCherry-Fb. J.M.G. and S.D.O. planned and conducted zebrafish experiments. V.V.V. planned and supervised the whole project, and together with N.V.B., S.D.O. and A.N. designed the experiments, analyzed the data, and wrote the manuscript. All authors reviewed the manuscript.

