## Supplementary information for "Synthetic multicolor antigen-stabilizable nanobody platform for intersectional labelling and functional imaging"

Supplemental Information

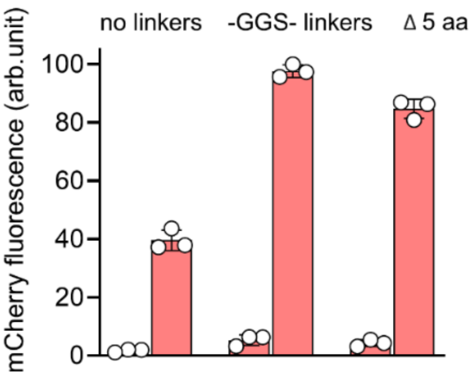

**Supplementary Figure 1. Evaluation of nanobodies with inserted mCherry in HeLa cells.** Red fluorescence intensity of cells transfected with mCherry inserted into $Nb_{GFP}$ without linkers (no linkers), with –GGS-linkers or with truncations of 5 a.a. from N- and C-ends (Δ 5 a.a.) and co-expressed with mEGFP (right column (+)) or mTagBFP2 (left column (-)). Fluorescence intensity was analyzed by flow cytometry using a 405 nm excitation laser and 450/50 nm emission filter for mTagBFP2; a 488 nm excitation laser and 525/50 nm emission filter for mEGFP; a 561 nm excitation laser and 610/20 nm emission filter for mCherry-$Fb_{GFP}$. The maximal fluorescence of antigen-bound form for mCherry-$Fb_{GFP}$ with GGS linkers was assumed to be 100%. Data are presented as mean values ± s.d. for $n = 3$ transfection experiments.

| | |
|---|---|
| mCherry-Fb$_{GFP}$ (no link) | MDQVQLVESGGALVQPGGSLRLSCAASGFPVNRYSMRWYRQAPGKEREWVAGMSSAGD |
| mCherry-Fb$_{GFP}$ (-GGS-) | MDQVQLVESGGALVQPGGSLRLSCAASGFPVNRYSMRWYRQAPGKEREWVAGMSSAGD |
| mCherry-Fb$_{GFP}$ (Δ5) | MDQVQLVESGGALVQPGGSLRLSCAASGFPVNRYSMRWYRQAPGKEREWVAGMSSAGD |
| mCherry-Fb$_{GFP}$ (no link) | RSSYEDS-----MVSKGEEDNMAIIKEFMRFKVHMEGSVNGHEFEIEGEGEGRPYEGT |
| mCherry-Fb$_{GFP}$ (-GGS-) | RSSYEDSGGGGSMVSKGEEDNMAIIKEFMRFKVHMEGSVNGHEFEIEGEGEGRPYEGT |
| mCherry-Fb$_{GFP}$ (Δ5) | RSSYEDS----------EEDNMAIIKEFMRFKVHMEGSVNGHEFEIEGEGEGRPYEGT |
| mCherry-Fb$_{GFP}$ (no link) | QTAKLKVTKGGPLPFAWDILSPQFMYGSKAYVKHPADIPDYLKLSFPEGFKWERVMNF |
| mCherry-Fb$_{GFP}$ (-GGS-) | QTAKLKVTKGGPLPFAWDILSPQFMYGSKAYVKHPADIPDYLKLSFPEGFKWERVMNF |
| mCherry-Fb$_{GFP}$ (Δ5) | QTAKLKVTKGGPLPFAWDILSPQFMYGSKAYVKHPADIPDYLKLSFPEGFKWERVMNF |
| mCherry-Fb$_{GFP}$ (no link) | EDGGVVTVTQDSSLQDGEFIYKVKLRGTNFPSDGPVMQKKTMGWEASSERMYPEDGAL |
| mCherry-Fb$_{GFP}$ (-GGS-) | EDGGVVTVTQDSSLQDGEFIYKVKLRGTNFPSDGPVMQKKTMGWEASSERMYPEDGAL |
| mCherry-Fb$_{GFP}$ (Δ5) | EDGGVVTVTQDSSLQDGEFIYKVKLRGTNFPSDGPVMQKKTMGWEASSERMYPEDGAL |
| mCherry-Fb$_{GFP}$ (no link) | KGEIKQRLKLKDGGHYDAEVKTTYKAKKPVQLPGAYNVNIKLDITSHNEDYTIVEQYE |
| mCherry-Fb$_{GFP}$ (-GGS-) | KGEIKQRLKLKDGGHYDAEVKTTYKAKKPVQLPGAYNVNIKLDITSHNEDYTIVEQYE |
| mCherry-Fb$_{GFP}$ (Δ5) | KGEIKQRLKLKDGGHYDAEVKTTYKAKKPVQLPGAYNVNIKLDITSHNEDYTIVEQYE |
| mCherry-Fb$_{GFP}$ (no link) | RAEGRHSTGGMDELYK-----VKGRFTISRDDARNTVYLQMNSLKPEDTAVYYCNVNV |
| mCherry-Fb$_{GFP}$ (-GGS-) | RAEGRHSTGGMDELYKGGGGSVKGRFTISRDDARNTVYLQMNSLKPEDTAVYYCNVNV |
| mCherry-Fb$_{GFP}$ (Δ5) | RAEGRHSTGGM----------VKGRFTISRDDARNTVYLQMNSLKPEDTAVYYCNVNV |
| mCherry-Fb$_{GFP}$ (no link) | GFEYWGQGTQVTVSS |
| mCherry-Fb$_{GFP}$ (-GGS-) | GFEYWGQGTQVTVSS |
| mCherry-Fb$_{GFP}$ (Δ5) | GFEYWGQGTQVTVSS |

**Supplementary Figure 2. Alignment of the amino acid sequences for mCherry-Fb$_{GFP}$ without linkers (no link), with GGS linkers (-GGS-) and with deletions of 5 a.a. from N- and C-termini (Δ5).** CDRs in Nb sequences are highlighted in blue. The sequence of inserted mCherry is highlighted with red. Linkers between Nbs and mCherry are highlighted in grey.

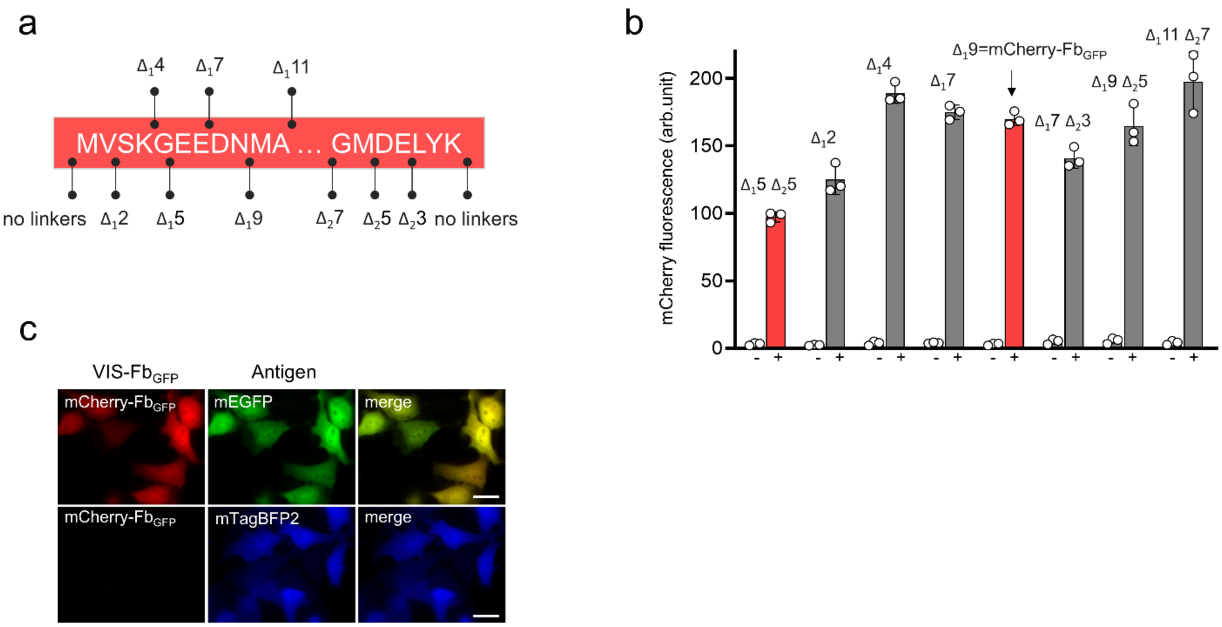

**Supplementary Figure 3. Optimization of mCherry termini in mCherry-Fb$_{GFP}$. (a)** Scheme of truncations from N- or/and C-end of mCherry part of mCherry-Fb$_{GFP}$. Black arrows indicate starting positions for truncations. **(b)** Red fluorescence intensity of cells transfected with mCherry-Fb$_{GFP}$ with truncations indicated in (a) and co-expressed with mEGFP (right column (+)) or mTagBFP2 (left column (-)). **(c)** Fluorescence images of HeLa cells co-expressing mCherry-Fb$_{GFP}$ with mTagBFP2 (negative control) or mEGFP (positive control). In (b), fluorescence intensity was analyzed by flow cytometry using a 405 nm excitation laser and 450/50 nm emission filter for mTagBFP2; a 488 nm excitation laser and 525/50 nm emission filter for mEGFP; a 561 nm excitation laser and 610/20 nm emission filter for mCherry-Fb$_{GFP}$. The maximal fluorescence of antigen-bound form for mCherry-Fb$_{GFP}$ with 5 truncated a.a. from each end was assumed to be 100%. Data are presented as mean values ± s.d. for $n = 3$ transfection experiments. In (c), for imaging of mTagBFP2, mEGFP, and mCherry-Fb$_{GFP}$, 390/40 nm excitation and 460/40 nm emission, 480/40 nm excitation and 535/40 nm emission, and 575/25 nm excitation and 615/30 nm emission filters were used, respectively. Scale bar, 40 μm.

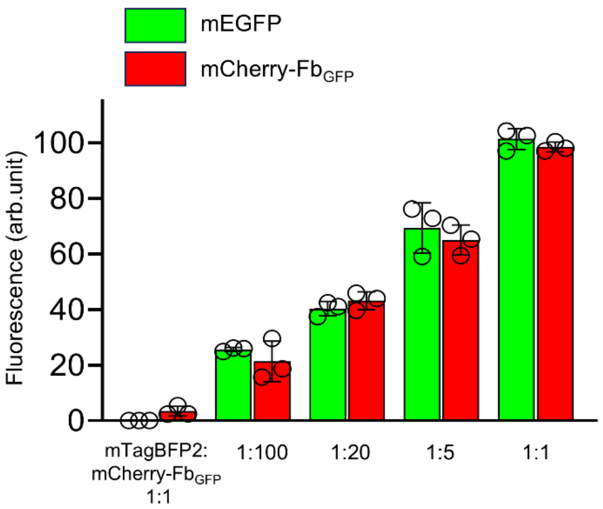

**Supplementary Figure 4. Fluorescence intensity of cells transfected with the fixed amount of the pmCherry-Fb$_{GFP}$ plasmid and varying ratios of the pmEGFP-N1 plasmid relative to the pmCherry-Fb$_{GFP}$ plasmid.** Fluorescence of mCherry-Fb$_{GFP}$ co-expressed with pmEGFP-N1 plasmid at a ratio of 1:1 was assumed to be 100%. mCherry-Fb$_{GFP}$ co-expressed with mTagBFP2 was assumed as 0%. Data are presented as mean values ± s.d. for $n = 3$ transfection experiments. Fluorescence intensity was analyzed by flow cytometry using a 405 nm excitation laser and 450/50nm emission filter for mTagBFP2, 488 nm excitation laser and 510/20 nm emission filter for mEGFP, a 561 nm excitation laser and 602/40 nm emission filter for mCherry-Fb$_{GFP}$.

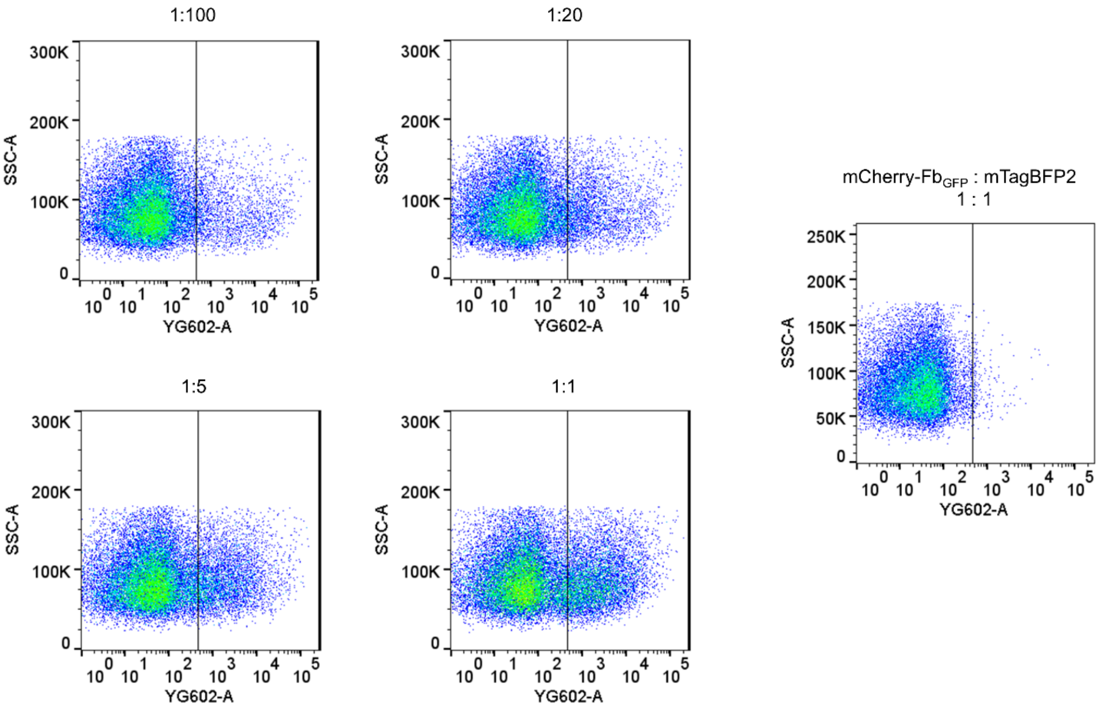

**Supplementary Figure 5. The intensity dot plot of cells transfected with the fixed amount of the pmCherry-Fb$_{GFP}$ plasmid and varying ratios of the pmEGFP-N1 plasmid relative to the pmCherry-Fb$_{GFP}$ plasmid.** Fluorescence intensity was analyzed by flow cytometry using a 405 nm excitation laser and 450/50 nm emission filter for mTagBFP2, 488 nm excitation laser and 510/20 nm emission filter for mEGFP, a 561 nm excitation laser and 602/40 emission filter for mCherry-Fb$_{GFP}$. The data are presented as the intensity dot plots of mCherry-Fb$_{GFP}$ (abscissa axis) versus Side Scatter Area (SSC-A) (ordinate axis) fluorescence.

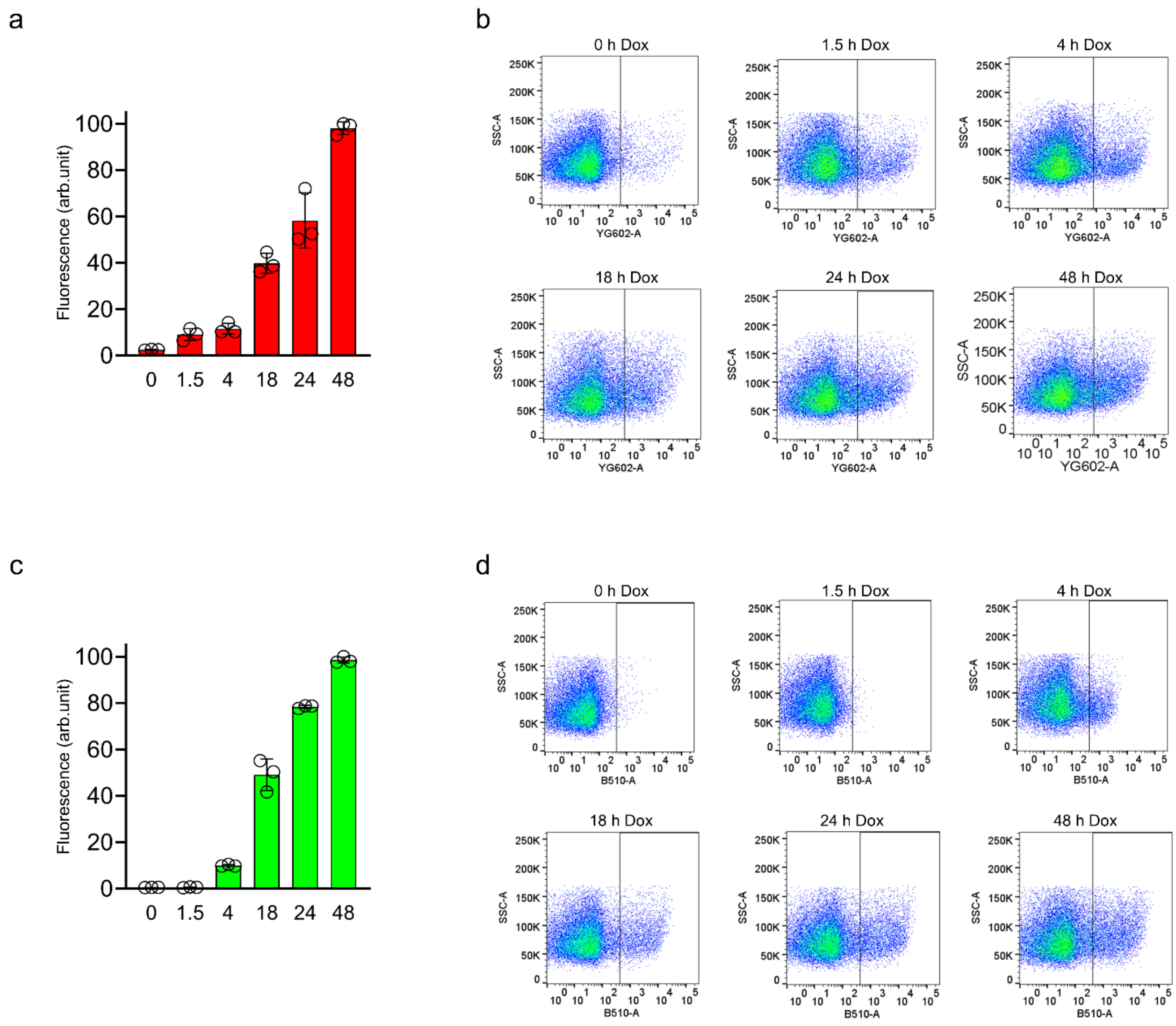

**Supplementary Figure 6. Time course of antigen-dependent mCherry-Fb$_{GFP}$ fluorescence in response to doxycycline-induced mEGFP expression in HeLa cells. (a)** Red fluorescence of HeLa cells co-expressing mCherry-Fb$_{GFP}$, mEGFP under tetracycline-inducible (TRE) promoter, and Tet On plasmid in response to 1 μg/mL doxycycline (Dox) for 0; 1.5; 4; 18; 24 or 48 h. The maximal fluorescence of HeLa cells treated with Dox for 48 h was assumed to be 100%. **(b)** The data from (a) are presented as the intensity dot plots of mCherry-Fb$_{GFP}$ (abscissa axis) versus Side Scatter Area (SSC-A) (ordinate axis) fluorescence. **(c)** Green fluorescence of HeLa cells co-expressing mCherry-Fb$_{GFP}$, mEGFP under tetracycline-inducible (TRE) promoter, and Tet On plasmid in response to 1 μg/mL doxycycline for 0; 1.5; 4; 18; 24 or 48 h. The maximal fluorescence of HeLa cells treated with Dox for 48 h was assumed to be 100%. **(d)** The data from (c) are presented as the intensity dot plots of mEGFP (abscissa axis) versus Side Scatter Area (SSC-A) (ordinate axis) fluorescence. Fluorescence intensity was analyzed by flow cytometry using a 488 nm excitation laser and 510/20 nm emission filter for mEGFP, a 561 nm excitation laser and 602/40 emission filter for mCherry-Fb$_{GFP}$. (a and c) Data are presented as mean values ± s.d. for $n = 3$ transfection experiments.

a

```
wtGFP         M-SKGEE----LFTGVVPI
mTagBFP2      MVSKGEE--MALIKENMHM
mTFP1         MVSKGEETTMGVIKPDMKI
mWasabi       MVSKGEETTMGVIKPDMKI
mNeonGreen    MVSKGEEDNMASLPATHEL
mOrange       MVSKGEENNMAIIKEFMRF
LSSmOrange    MVSKGEENNMAIIKEFMRF
CyOFP1        MVSKGEE--MALIKENMRS
mScarlet-I    MVSKGEA--MAVIKEFMRF
mScarlet      MVSKGEA--MAVIKEFMRF
LSSmScarlet   MVSKGEA--MAVIKEFMRF
mCherry       MVSKGEEDNMAIIKEFMRF
mNeptune2     MVSKGEE--MALIKENMHM
mCardinal     MVSKGEE--MALIKENMHM
PAmCherry     MVSKGEEDNMAIIKEFMRF
mEos4a        MVS------MAAIKPDMRI
```

b

```
wtGFP         VLLEFVTAAGITL--GMDELYK
mTagBFP2      EQHEVAVARYCDLP--SKLGHKLN
mTFP1         TVYESAVARNST--DGMDELYK
mWasabi       TVYEIAVARNST--DGMDELYK
mNeonGreen    NFKEWQKAFTDV--MGMDELYK
mOrange       EQYERAEGRHST--GGMDELYK
LSSmOrange    EQYERAEGRHST--GGMDELYK
CyOFP1        EQYEHAVARYSNLGGGMDELYK
mScarlet-I    EQYERSEGRHST--GGMDELYK
mScarlet      EQYERSEGRHST--GGMDELYK
LSSmScarlet   EQYERSEGRHST--GGMDELYK
mCherry       EQYERAEGRHST--GGMDELYK
mNeptune2     EQHEVAVARYCDLP--SKLGHKLNGMDELYK
mCardinal     EQHEVAVARYCDLP--SKLGHKLNGMDELYK
PAmCherry     EQYERAEGRHST--GGMDELYK
mEos4a        KLYEHAVA-HSGL--PDNARRYK
```

**Supplementary Figure 7. Alignment of the amino acid sequences for N- and C-ends of fluorescent proteins inserted into a nanobody for GFP. (a)** N-ends of fluorescent proteins inserted into an Nb for GFP **(b)** C-ends of fluorescent proteins inserted into an Nb for GFP. (a-b) The truncated a.a. are highlighted in red, and the inserted a.a. are highlighted in grey.

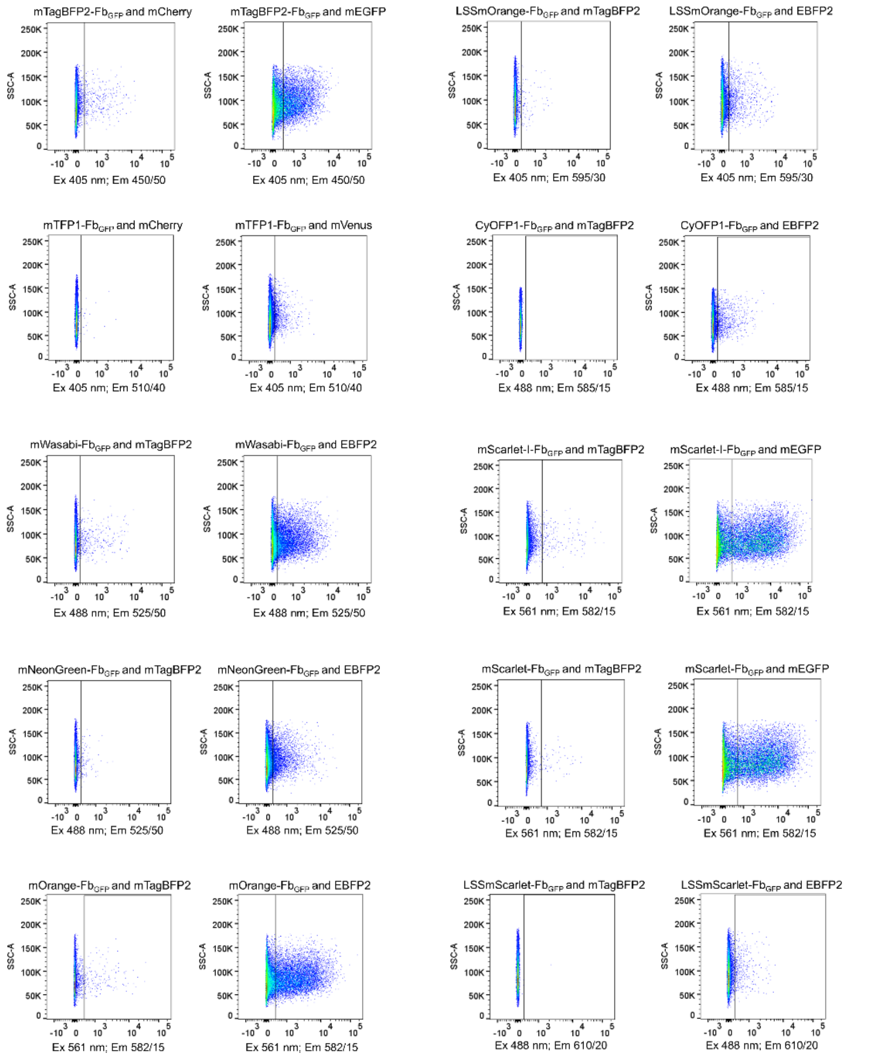

**Supplementary Figure 8. The intensity dot plots for cells transfected with 13 different VIS-Fbs to GFP from Figure 1b.** The following FPs were co-expressed with VIS-Fbs as positive controls: *mEGFP* for mTagBFP2-Fb$_{GFP}$, mScarlet-I-Fb$_{GFP}$, mScarlet-Fb$_{GFP}$, mCherry-Fb$_{GFP}$, mNeptune2-Fb$_{GFP}$, mCardinal-Fb$_{GFP}$; *mVenus* for mTFP1-Fb$_{GFP}$; *EBFP2* for mWasabi-Fb$_{GFP}$, mNeonGreen-Fb$_{GFP}$, mOrange-Fb$_{GFP}$, LSSmOrange-Fb$_{GFP}$, CyOFP1-Fb$_{GFP}$, LSSmScarlet-Fb$_{GFP}$. The following FPs were co-expressed with VIS-Fbs as negative controls: *mCherry* for mTagBFP2-Fb$_{GFP}$, mTFP1-Fb$_{GFP}$; *mTagBFP2* for other VIS-Fbs. Fluorescence intensity was analyzed by flow cytometry using a 405 nm excitation laser and 450/50 nm emission filter for mTagBFP2, EBFP2, mTagBFP2-Fb$_{GFP}$, 510/40 nm emission filter for mTFP1-Fb$_{GFP}$, 595/30 emission filter for LSSmOrange-Fb$_{GFP}$; a 488 nm excitation laser and 525/50 nm emission filter for mEGFP, mWasabi-Fb$_{GFP}$, mNeonGreen-Fb$_{GFP}$, 537/32 nm for mVenus; 582/15 nm emission filter for CyOFP1-Fb$_{GFP}$, 610/20 nm emission filter for LSSmScarlet-Fb$_{GFP}$; a 561 nm excitation laser and 582/15 nm emission filter for mOrange-Fb$_{GFP}$, mScarlet-Fb$_{GFP}$ and mScarlet-I-Fb$_{GFP}$; 610/20 nm emission filter for mCherry and mCherry-Fb$_{GFP}$; 670/30 nm emission filter for mNeptune2-Fb$_{GFP}$ and mCardinal-Fb$_{GFP}$. The data are presented as the intensity dot plots of corresponding VIS-Fb$_{GFP}$ (abscissa axis) versus Side Scatter Area (SSC-A) (ordinate axis) fluorescence.

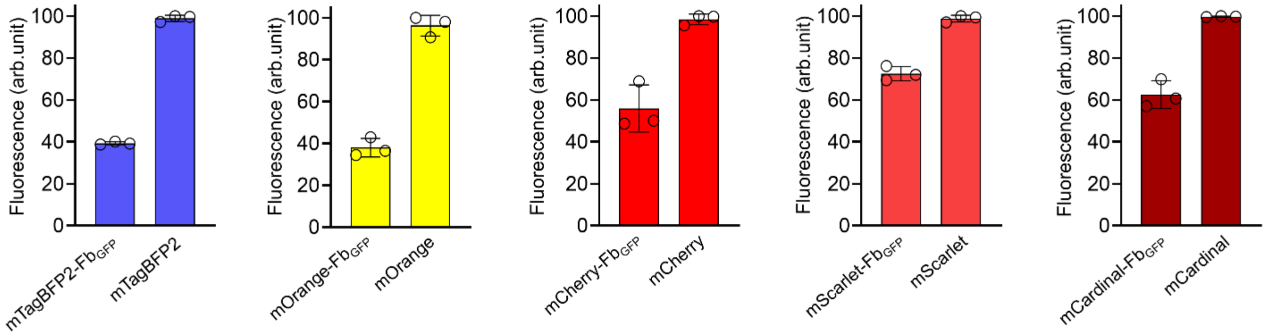

**Supplementary Figure 9. Comparison of the fluorescent intensity between representative VIS-Fbs$_{GFP}$ in antigen-stabilized condition and respective fluorescent proteins (FPs) that were inserted into the Nb$_{GFP}$.** HeLa cells were co-transfected with VIS-Fb or the respective parental FP with mEGFP, or EBFP2 in the case of mOrange-Fb$_{GFP}$, at a 1:1 ratio. Fluorescence intensity was analyzed by flow cytometry using a 405 nm excitation laser and 450/40 nm emission filter for mTagBFP2, EBFP2 and mTagBFP2-Fb$_{GFP}$; a 488 nm excitation laser and 510/20 nm emission filter for mEGFP; a 561 nm excitation laser and 585/30 nm emission filter for mOrange and mOrange-Fb$_{GFP}$; 602/40 nm emission filter for mCherry, mScarlet, mCherry-Fb$_{GFP}$ and mScarlet-Fb$_{GFP}$; 660/30 nm emission filter for mCardinal and mCardinal-Fb$_{GFP}$. The maximal fluorescence of corresponding parental FP was assumed to be 100%. Data are presented as mean values ± s.d. for $n = 3$ transfection experiments.

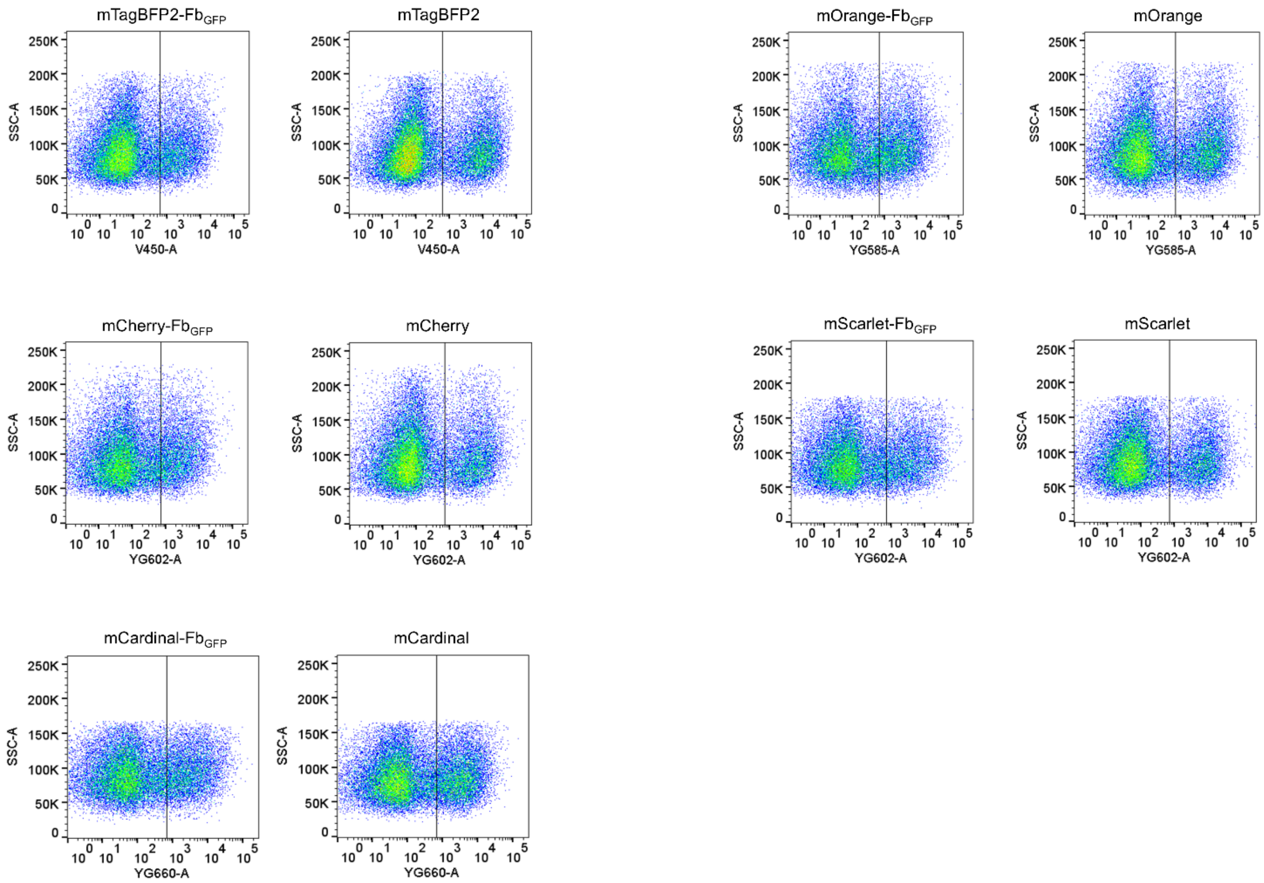

**Supplementary Figure 10. The intensity dot plots for representative VIS-Fbs$_{GFP}$ in antigen-stabilized condition and respective fluorescent proteins (FPs) that were inserted into the Nb$_{GFP}$ in transfected HeLa cells.** HeLa cells were co-transfected with VIS-Fb for GFP or the respective FP with mEGFP, or EBFP2 in the case of mOrange-Fb$_{GFP}$, at a 1:1 ratio. Fluorescence intensity was analyzed by flow cytometry using a 405 nm excitation laser and 450/40 nm emission filter for mTagBFP2, EBFP2 and mTagBFP2-Fb$_{GFP}$; a 488 nm excitation laser and 510/20 nm emission filter for mEGFP; a 561 nm excitation laser and 585/30 nm emission filter for mOrange and mOrange-Fb$_{GFP}$; 602/40 nm emission filter for mCherry, mScarlet, mCherry-Fb$_{GFP}$ and mScarlet-Fb$_{GFP}$; 660/30 nm emission filter for mCardinal and mCardinal-Fb$_{GFP}$. The data are presented as the intensity dot plots of corresponding VIS-Fb$_{GFP}$ or FP (abscissa axis) versus Side Scatter Area (SSC-A) (ordinate axis) fluorescence.

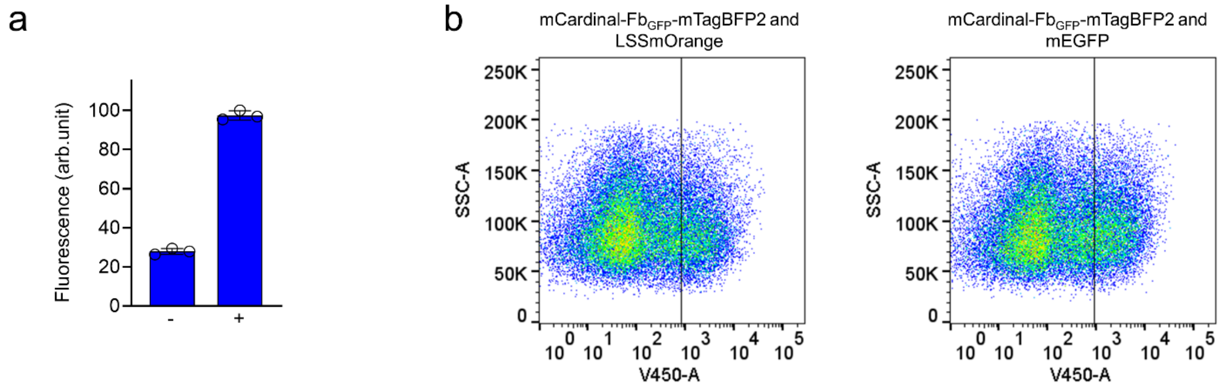

**Supplementary Figure 11. Fluorescence intensity of mCardinal-Fb$_{GFP}$ fused to mTagBFP2 in antigen-free and antigen-bound states. (a)** Fluorescence intensity of HeLa cells expressing mCardinal-Fb$_{GFP}$ fused with mTagBFP2 co-transfected with either mEGFP (+) or LSSmOrange (-). Data are presented as mean values ± s.d. for n = 3 transfection experiments. **(b)** The data from (a) are presented as the intensity dot plots of mCardinal-Fb$_{GFP}$ fused with mTagBFP2 in the blue channel (abscissa axis) versus Side Scatter Area (SSC-A) (ordinate axis) fluorescence. The fluorescence intensity was analyzed by flow cytometry using a 405 nm excitation laser and a 450/40 nm emission filter. The mTagBFP2 brightness in cells co-transfected with mEGFP was assumed to be 100%.

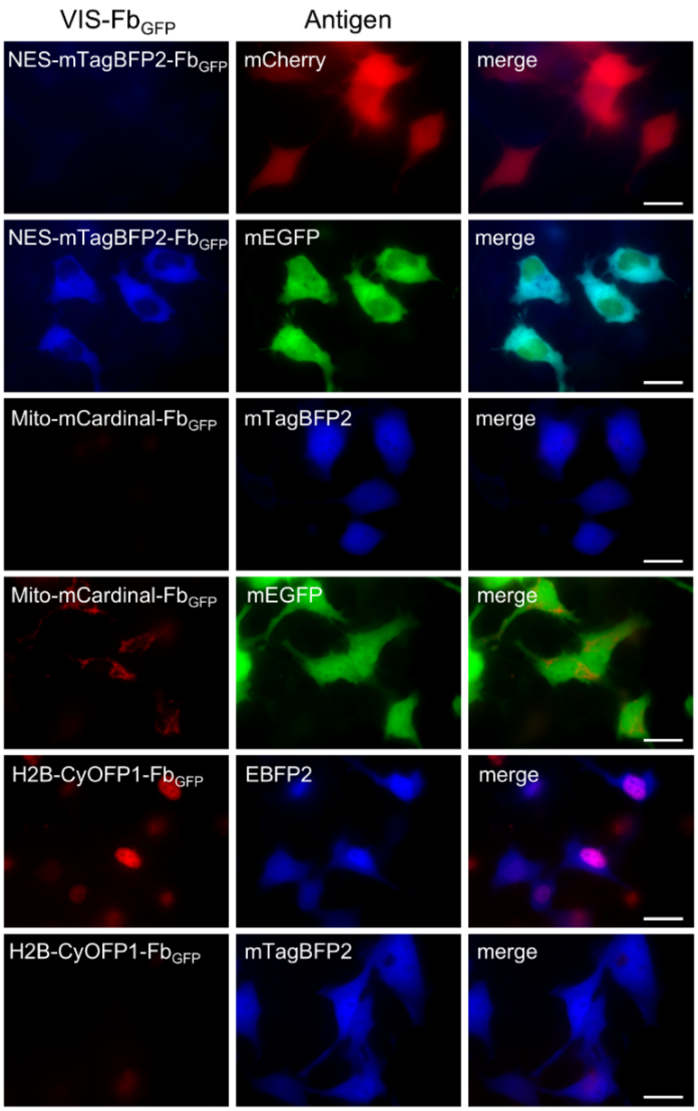

**Supplementary Figure 12. Controls for multicolor imaging with VIS-Fbs$_{GFP}$ in HeLa cells.** HeLa cells co-expressing NES-mTagBFP2-Fb$_{GFP}$ with mEGFP or mCherry, H2B-CyOFP1-Fb$_{GFP}$ with EBFP2 or mTagBFP2, mito-mCardinal-Fb$_{GFP}$ with mEGFP or mTagBFP2. The following filters were used: for imaging NES-mTagBFP2-Fb$_{GFP}$ and mTagBFP2 390/40 nm excitation and 460/40 nm emission; for imaging mEGFP 480/40 nm excitation and 535/40 nm emission; for imaging H2B-CyOFP1-Fb$_{GFP}$ 523/20 nm excitation and 588/21 nm emission; for imaging mCherry 575/25 nm excitation and 615/30 nm emission; for imaging mito-mCardinal-Fb$_{GFP}$ 605/30 nm excitation and 667/30 nm emission. Scale bar, 40 μm.

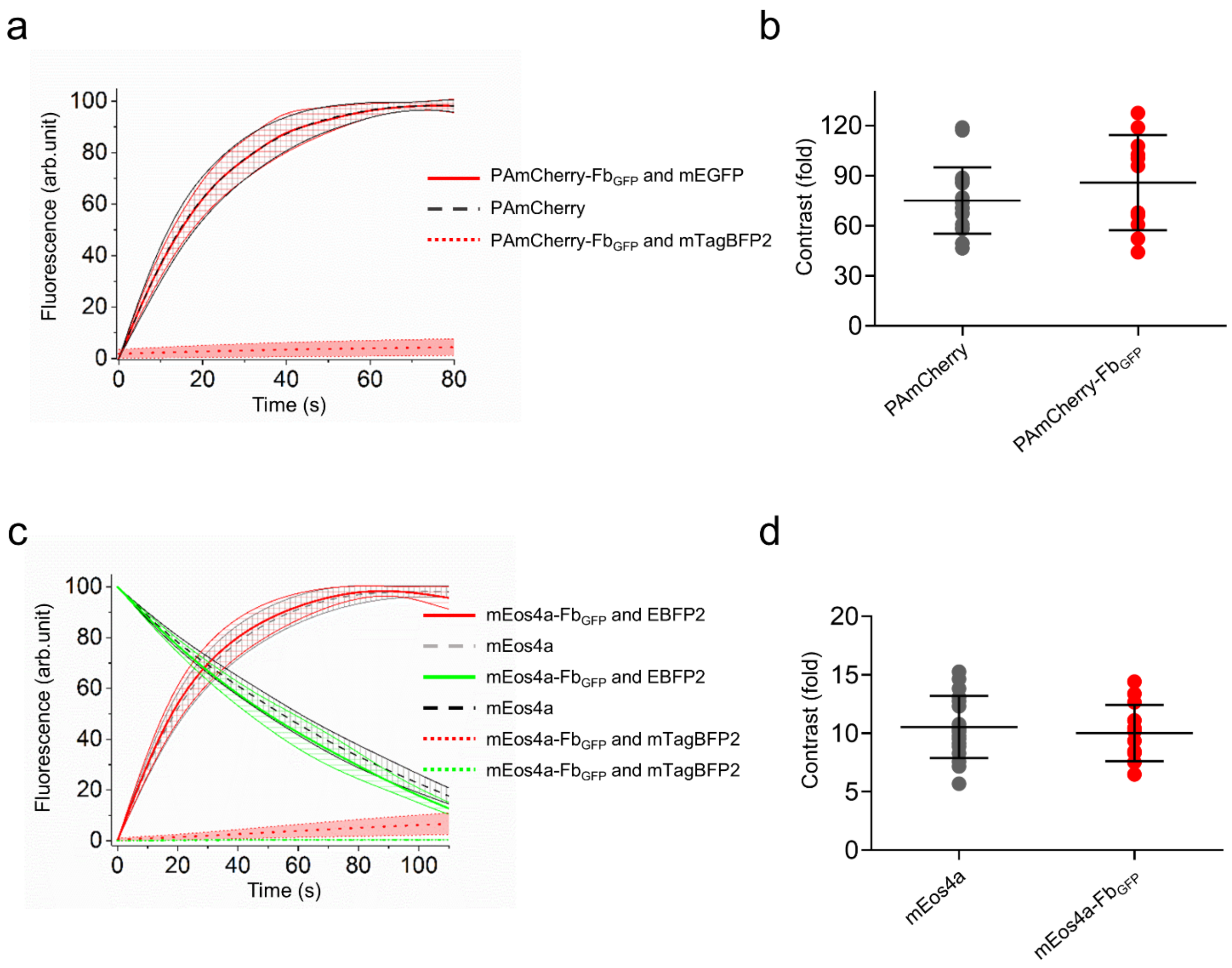

**Supplementary Figure 13. Comparison of activation kinetics and contrast for photoactivatable and photoswitchable fluorescent nanobodies with their parental fluorescent proteins expressed in HeLa cells. (a)** Change of fluorescence over time for PAmCherry FP (dashed black, *n=21*) and PAmCherry-based VIS-Fb$_{GFP}$ co-expressed with mEGFP (red, *n=11*) or mTagBFP2 (dotted red, *n=18*) in response to 390 nm irradiation. **(b)** Comparison of absolute contrasts for PAmCherry (*n=21*) and PAmCherry-Fb$_{GFP}$ co-expressed with mEGFP (*n=11*) between dark and light-activated forms. **(c)** Change of fluorescence over time for mEos4a FP (red form: dashes light grey, *n=16*; green form: dashes black, *n=16*) and mEos4a-based VIS-Fb$_{GFP}$ co-expressed with EBFP2 (red form: red, *n=13*; green from: green, *n=13*) or mTagBFP2 (red from: dotted red, *n=15*; green form: dotted green, *n=15*) in response to 390 nm irradiation. **(d)** Comparison of absolute green forms' contrasts for mEos4a (*n=16*) and mEos4a-Fb$_{GFP}$ co-expressed with EBFP2 (*n=13*) between dark and light-activated forms. The following filters were used: for imaging green form of mEos4a and mEos4a-Fb$_{GFP}$ 480/40 nm excitation and 535/40 nm emission; for imaging red form of mEos4a and mEos4a-Fb$_{GFP}$, PAmCherry, and PAmCherry-based VIS-Fb$_{GFP}$ 575/25 nm excitation and 615/30 nm emission. Data is presented as mean values ± s.d.

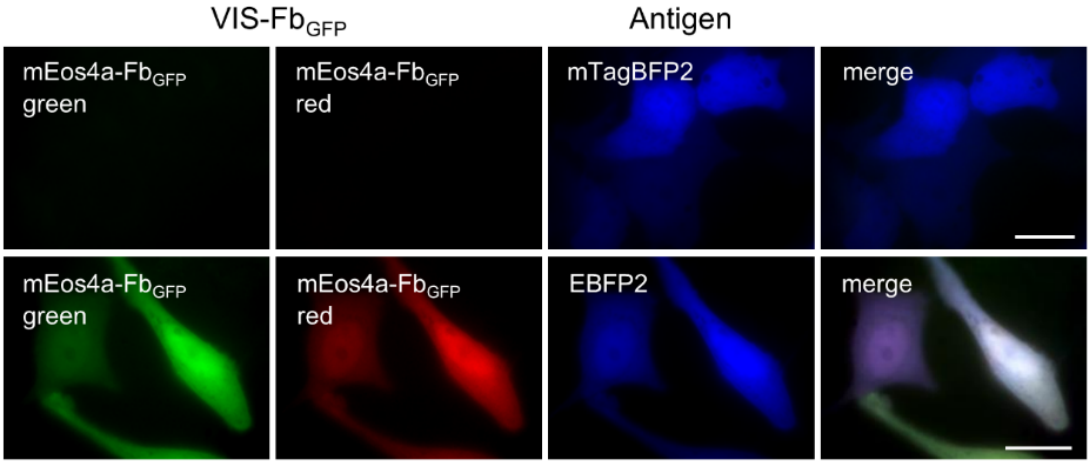

**Supplementary Figure 14. Controls for mEos4a-Fb$_{GFP}$ in HeLa cells.** The following filters were used: for imaging mTagBFP2 and EBFP2 390/40 nm excitation and 460/40 nm emission; for imaging mEos4a-Fb$_{GFP}$ green form 480/40 nm excitation and 535/40 nm emission; for imaging mEos4a-Fb$_{GFP}$ red form 575/25 nm excitation and 615/30 nm emission. Scale bar, 40 µm.

a

```
mTagBFP2-Fb_LAM2   MAQVQLVESGGGLVQAGGSLRLSCATSGFTFSDYAMGWFRQAPGKEREFVAAISWSGHVTDYADSMVSKGEE--MALIKENMHMK
mWasabi-Fb_LAM2    MAQVQLVESGGGLVQAGGSLRLSCATSGFTFSDYAMGWFRQAPGKEREFVAAISWSGHVTDYADSMVSKGEETTMGVIKPDMKIK
mClover3-Fb_LAM2   MAQVQLVESGGGLVQAGGSLRLSCATSGFTFSDYAMGWFRQAPGKEREFVAAISWSGHVTDYADSMVSKGEE--MALFTGVVPIL

mTagBFP2-Fb_LAM2   LYMEGTVDNHHFKCTSEGEGKPYEGTQTMRIKVVEGGPLPFAFDILATSFLYGSKTFINHTQGIP--DFFKQSFPEGFTWERVTT
mWasabi-Fb_LAM2    LKMEGNVNGHAFVIEGEGEGKPYDGTNTINLEVKEGAPLPFSYDILTTAFSYGNRAFTKYPDDIP--NYFKQSFPEGYSWERTMT
mClover3-Fb_LAM2   VELDGDVNGHKFSVRGEGEGDATNGKLTLKFICTTGK-LPVPWPTLVTTFGYGVACFSRYPDHMKQHDFFKSAMPEGYVQERTIS

mTagBFP2-Fb_LAM2   YEDGGVLTATQDTSLQDGCLIYNVKIRGVNFTSNGPVMQKKTLGWEAFTETLYP----ADGGLEGRNDMALKLVGGSHLIANA--
mWasabi-Fb_LAM2    FEDKGIVKVKSDISMEEDSFIYEIHLKGENFPPNGPVMQKETTGWDASTERMYV----RDGVLKGDVKMKLLLEGGGHHRVDF--
mClover3-Fb_LAM2   FKDDGTYKTRAEVKFEGDTLVNRIELKGIDFKEDGNILGHKLE-YNFNSHYVYITADKQKNCIKANFKIRHNVEDGSVQLADHYQ

mTagBFP2-Fb_LAM2   KTTYRSKKPAKNLKMPGVYYVDYRLERIKEANNET---YVEQHEVAVARYCDLPSKLGHKLNVKGRFTISRDNVKNTVYLQMNS
mWasabi-Fb_LAM2    KTIYRAKKA---VKLPDYHFVDHRIEIL---NHDKDYNKVTVYEIAVARNSTDGMDELYK--VKGRFTISRDNVKNTVYLQMNS
mClover3-Fb_LAM2   QNTPIGDGPVLLPDNHYLSHQSKLSKDPNEKRDHM--VLLEFVTAAGITH---GMDELYK--VKGRFTISRDNVKNTVYLQMNS

mTagBFP2-Fb_LAM2   LKPEDTAVYSCAAAKSGTWWYQRSENDFGSWGQGTQVTVS
mWasabi-Fb_LAM2    LKPEDTAVYSCAAAKSGTWWYQRSENDFGSWGQGTQVTVS
mClover3-Fb_LAM2   LKPEDTAVYSCAAAKSGTWWYQRSENDFGSWGQGTQVTVS
```

b

```
mTagBFP2(GGS)-Fb_LAM2
MAQVQLVESGGGLVQAGGSLRLSCATSGFTFSDYAMGWFRQAPGKEREFVAAISWSGHVTDYADSGGSMVSKGEELIKENMHMKLYMEGTVDNHHFKCTSE
GEGKPYEGTQTMRIKVVEGGPLPFAFDILATSFLYGSKTFINHTQGIPDFFKQSFPEGFTWERVTTYEDGGVLTATQDTSLQDGCLIYNVKIRGVNFTSNG
PVMQKKTLGWEAFTETLYPADGGLEGRNDMALKLVGGSHLIANAKTTYRSKKPAKNLKMPGVYYVDYRLERIKEANNETYVEQHEVAVARYCDLPSKLGHK
LNGGSVKGRFTISRDNVKNTVYLQMNSLKPEDTAVYSCAAAKSGTWWYQRSENDFGSWGQGTQVTVS
```

**Supplementary Figure 15. Alignments of the amino acid sequences for VIS-Fb$_{LAM2}$s. (a)** Alignment of amino acid sequences for mTagBFP2-Fb$_{LAM2}$, mWasabi-Fb$_{LAM2}$, mClover3-Fb$_{LAM2}$. mTagBFP2, mWasabi, and mClover3 sequences are highlighted in blue or green. **(b)** Alignment of amino acid sequence for mTagBFP2(GGS)-Fb$_{LAM2}$ with mTagBFP2 sequence highlighted in blue. (a, b) Truncated a.a. are highlighted in red, inserted a.a., including linkers, are highlighted in grey.

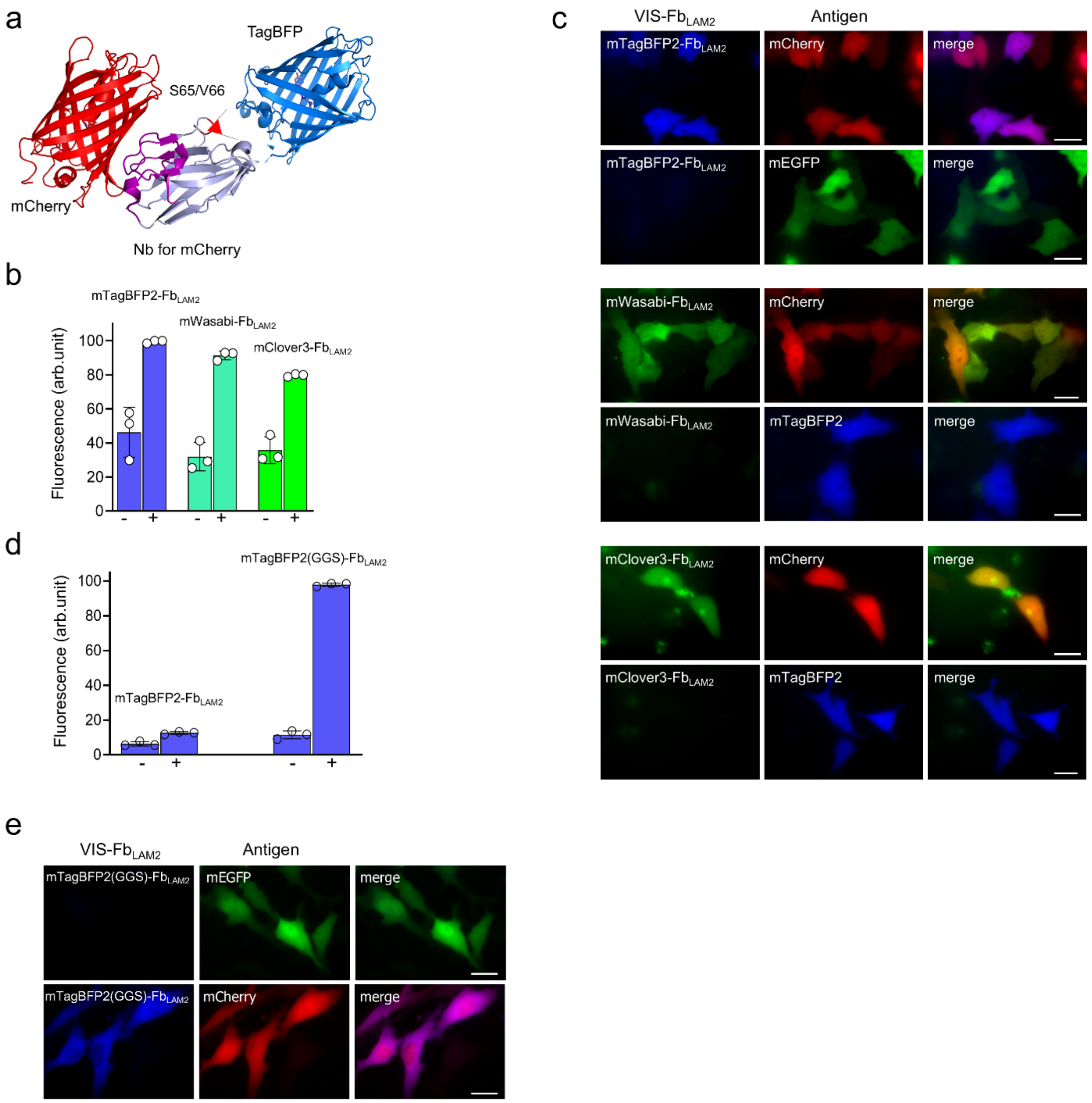

**Supplementary Figure 16. VIS-Fbs for mCherry. (a)** Scheme of a nanobody with TagBFP (PDB ID: 3M24) inserted into the nanobody for mCherry (PDB ID: 7SAJ) bound to mCherry FP. Complementarity-determining regions (CDRs) are highlighted in violet. The position of TagBFP insertion to the anti-mCherry nanobody is indicated with a red arrow. **(b)** Fluorescence intensity of cells expressing mTagBFP2-Fb$_{LAM2}$, mWasabi-Fb$_{LAM2}$, and mClover3-Fb$_{LAM2}$. For negative controls, mTagBFP2-Fb$_{LAM2}$ was co-expressed with mEGFP, mWasabi-Fb$_{LAM2}$ and mClover3-Fb$_{LAM2}$ – with mTagBFP2. The maximal fluorescence of HeLa cells co-expressing mTagBFP2-Fb$_{LAM2}$ with mCherry was assumed to be 100%. **(c)** Fluorescent images of mTagBFP2-Fb$_{LAM2}$, mWasabi-Fb$_{LAM2}$, and mClover3-Fb$_{LAM2}$ co-expressed with mCherry or mEGFP/mTagBFP2. **(d)** Fluorescence intensity of cells expressing mTagBFP2-Fb$_{LAM2}$ and mTagBFP2(GGS)$_{LAM2}$. For negative control, both VIS-Fb$_{LAM2}$s were co-expressed with mEGFP. The maximal fluorescence of HeLa cells co-expressing mTagBFP2-$Fb_{LAM2}$ with mCherry was assumed to be 100%. **(e)** Fluorescent images of mTagBFP2(GGS)-$Fb_{LAM2}$ co-expressed with mEGFP or mCherry. In (b and d) fluorescence intensity was analyzed by flow cytometry using a 405 nm excitation laser and 450/50 nm or 450/40 nm emission filter for mTagBFP2-$Fb_{LAM2}$, mTagBFP2(GGS)-$Fb_{LAM2}$, and mTagBFP2; a 488 nm excitation laser and 525/50 nm emission filter for mWasabi-$Fb_{LAM2}$, mClover3-$Fb_{LAM2}$, and mEGFP; a 561 nm excitation laser and 610/20 nm emission filter for mCherry. Data are presented as mean values ± s.d. for $n = 3$ transfection experiments. The following filters were used in (c and e): for imaging mTagBFP2-$Fb_{LAM2}$, mTagBFP2(GGS)-$Fb_{LAM2}$, and mTagBFP2 390/40 nm excitation and 460/40 nm emission; for imaging mWasabi-$Fb_{LAM2}$, mClover3-$Fb_{LAM2}$, and mEGFP 480/40 nm excitation and 535/40 nm emission; for imaging mCherry 575/25 nm excitation and 615/30 nm emission. Scale bar, 40 μm.

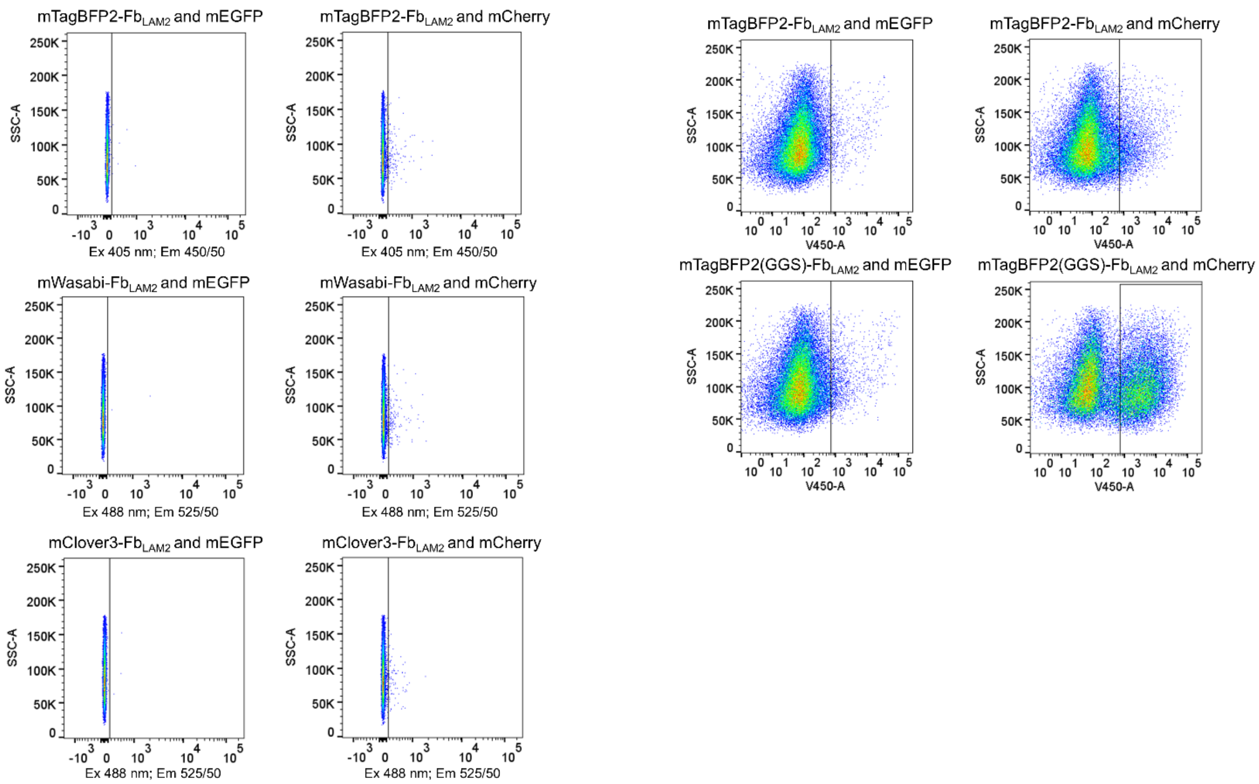

**Supplementary Figure 17. The intensity dot plots for VIS-Fb$_{LAM2}$s in antigen-stabilized or -destabilized conditions in transfected HeLa cells.** Fluorescence intensity was analyzed by flow cytometry using a 405 nm excitation laser and 450/50 nm or 450/40 nm emission filter for mTagBFP2-Fb$_{LAM2}$, mTagBFP2(GGS)-Fb$_{LAM2}$, and mTagBFP2; a 488 nm excitation laser and 525/50 nm emission filter for mWasabi-Fb$_{LAM2}$, mClover3-Fb$_{LAM2}$, and mEGFP; a 561 nm excitation laser and 610/20 nm emission filter for mCherry. The data are presented as the intensity dot plots of VIS-Fb$_{LAM2}$s (abscissa axis) versus Side Scatter Area (SSC-A) (ordinate axis) fluorescence.

**a**

mCardinal-$Fb_{59H10}$

MDVQLQESGGGLVQAGGSLRLSCAASGSISRFNAMGWWRQAPGKEREFVARIVKGFDPVLADSGGSMVSKGEELIKENM
HMKLYMEGTVNNHHFKCTTEGEGKPYEGTQTQRIKVVEGGPLPFAFDILATCFMYGSKTFINHTQGIPDFFKQSFPEGF
TWERVTTYEDGGVLTVTQDTSLQDGCLIYNVKLRGVNFPSNGPVMQKKTLGWEATTETLYPADGGLEGRCDMALKLVGG
GHLHCNLKTTYRSKKPAKNLKMPGVYFVDRRLERIKEADNETYVEQHEVAVARYCDLPSKLGHKLNGMDELYKGGSVKG
RFTISIDSAENTLALQMNRLKPEDTAVYYCFAALDTAYWGQGTQVTVSS

**b**

mTagBFP2-$Fb_{2E7}$

MDVQLQESGGGLVQPGGSLRLSCAASGNIVSIDAAGWFRQAPGKQREPVATILTGGATNYADSGGSMVSKGEELIKENM
HMKLYMEGTVDNHHFKCTSEGEGKPYEGTQTMRIKVVEGGPLPFAFDILATSFLYGSKTFINHTQGIPDFFKQSFPEGF
TWERVTTYEDGGVLTATQDTSLQDGCLIYNVKIRGVNFTSNGPVMQKKTLGWEAFTETLYPADGGLEGRNDMALKLVGG
SHLIANAKTTYRSKKPAKNLKMPGVYYVDYRLERIKEANNETYVEQHEVAVARYCDLPSKLGHKLNGGSVKGRFTISRD
NAKNTVYLQMNSLKPEDTAVYYCYAPMIYYGGRYSDYWGQGTQVTVSS

**c**

| | |
|---|---|
| mTagBFP2-$Fb_{ALFA}$ | MVQLQESGGGLVQPGGSLRLSCTASGVTISALNAMAMGWYRQAPGERRVMVAAVSERGNAMYR |
| CyOFP1-$Fb_{ALFA}$ | MVQLQESGGGLVQPGGSLRLSCTASGVTISALNAMAMGWYRQAPGERRVMVAAVSERGNAMYR |
| TagRFP-T-$Fb_{ALFA}$ | MVQLQESGGGLVQPGGSLRLSCTASGVTISALNAMAMGWYRQAPGERRVMVAAVSERGNAMYR |
| mScarlet-$Fb_{ALFA}$ | MVQLQESGGGLVQPGGSLRLSCTASGVTISALNAMAMGWYRQAPGERRVMVAAVSERGNAMYR |
| mStayGold-$Fb_{ALFA}$ | MVQLQESGGGLVQPGGSLRLSCTASGVTISALNAMAMGWYRQAPGERRVMVAAVSERGNAMYR |

| | |
|---|---|
| mTagBFP2-$Fb_{ALFA}$ | ESGGSMVSKGEELIKENMHMKLYMEGTVDNHHFKCTSEGEGKPYEGTQTMRIKVVEGGPLPFA |
| CyOFP1-$Fb_{ALFA}$ | ESGGSMVSKGEELIKENMRSKLYLEGSVNGHQFKCTHEGEGKPYEGKQTNRIKVVEGGPLPFA |
| TagRFP-T-$Fb_{ALFA}$ | ESGGSMVSKGEELIKENMHMKLYMEGTVNNHHFKCTSEGEGKPYEGTQTMRIKVVEGGPLPFA |
| mScarlet-$Fb_{ALFA}$ | ESGGSMVSKGEAVIKEFMRFKVHMEGSMNGHEFEIEGEGEGRPYEGTQTAKLKVTKGGPLPFS |
| mStayGold-$Fb_{ALFA}$ | ESGGSMVSTGEELFTGVVPFKFQLKGTINGKSFTVEGEGEGNSHEGSHKGKYVCTSGKLPMSW |

| | |
|---|---|
| mTagBFP2-$Fb_{ALFA}$ | FDILATSFLYGSKTFINHTQGIPDFFKQSFPEGFTWERVTTYEDGGVLTATQDTSLQDGCLIY |
| CyOFP1-$Fb_{ALFA}$ | FDILATHFMYGSKVFIKYPADLPDYFKQSFPEGFTWERVMVFEDGGVLTATQDTSLQDGELIY |
| TagRFP-T-$Fb_{ALFA}$ | FDILATSFMYGSRTFINHTQGIPDFFKQSFPEGFTWERVTTYEDGGVLTATQDTSLQDGCLIY |
| mScarlet-$Fb_{ALFA}$ | WDILSPQFMYGSRAFTKHPADIPDYYKQSFPEGFKWERVMNFEDGGAVTVTQDTSLEDGTLIY |
| mStayGold-$Fb_{ALFA}$ | AALGTSFGYGMKYYTKYPSGLKNWFHEVMPEGFTYDRHIQYKGDGSIHAKHQHFMKNGTYHNI |

| | |
|---|---|
| mTagBFP2-$Fb_{ALFA}$ | NVKIRGVNFTSNGPVMQKKTLGWEAFTETLYPADGGLEGRNDMALKLVGGSHLIANAKTTYR |
| CyOFP1-$Fb_{ALFA}$ | NVKVRGVNFPANGPVMQKKTLGWEPSTETMYPADGGLEGRCDKALKLVGGGHLHVNFKTTYK |
| TagRFP-T-$Fb_{ALFA}$ | NVKIRGVNFPSNGPVMQKKTLGWEANTETLYPADGGLEGRTDMALKLVGGGHLICNFKTTYR |
| mScarlet-$Fb_{ALFA}$ | KVKLRGTNFPPDGPVMQKKTMGWEASTERLYPEDGVLKGDIKMALRLKDGGRYLADFKTTYK |
| mStayGold-$Fb_{ALFA}$ | VEFTGQDFKENSPVLTGDMDVSLPNEVQHIPIDDGVECTVTLQYPLLSDESKCVEAYQNTII |

| | |
|---|---|
| mTagBFP2-$Fb_{ALFA}$ | SKKPAKNLKMPGVYYVDYRLERIKEANNETYVEQHEV-AVARYCDLPSKLGHKLN------- |
| CyOFP1-$Fb_{ALFA}$ | SKKPV---KMPGVHYVDRRLERIKEADNETYVEQYEH-AVARYSNL-------GGGMDELYK |
| TagRFP-T-$Fb_{ALFA}$ | SKKPAKNLKMPGVYYVDHRLERIKEADKETYVEQHEV-AVARYCDLPSKLGHKLNGMDELYK |
| mScarlet-$Fb_{ALFA}$ | AKKPV---QMPGAYNVDRKLDITSHNEDYTVVEQYER-SEGRHSTGGMDELYK--------- |
| mStayGold-$Fb_{ALFA}$ | KPLHN---QPAPDVPFHWIRKQYTQSKDDTEERDHIIQSETLEAHL---------------- |

| | |
|---|---|
| mTagBFP2-$Fb_{ALFA}$ | GGSVQGRFTVTRDFTNKMVSLQMDNLKPEDTAVYYCHVLEDRVDSFHDYWGQGTQVTVSS |
| CyOFP1-$Fb_{ALFA}$ | GGSVQGRFTVTRDFTNKMVSLQMDNLKPEDTAVYYCHVLEDRVDSFHDYWGQGTQVTVSS |
| TagRFP-T-$Fb_{ALFA}$ | GGSVQGRFTVTRDFTNKMVSLQMDNLKPEDTAVYYCHVLEDRVDSFHDYWGQGTQVTVSS |
| mScarlet-$Fb_{ALFA}$ | GGSVQGRFTVTRDFTNKMVSLQMDNLKPEDTAVYYCHVLEDRVDSFHDYWGQGTQVTVSS |
| mStayGold-$Fb_{ALFA}$ | GGSVQGRFTVTRDFTNKMVSLQMDNLKPEDTAVYYCHVLEDRVDSFHDYWGQGTQVTVSS |

**Supplementary Figure 18. Amino acid sequences for VIS-Fbs for p24, gp41, and ALFA-tag. (a)** Amino acid sequence for mCardinal-$Fb_{59H10}$. mCardinal sequence is highlighted with pink. **(b)** Amino acid sequence for mTagBFP2-$Fb_{2E7}$. mTagBFP2 sequence is highlighted with blue. **(c)** Alignment of amino acid sequences for VIS-$Fb_{ALFA}$s. mTagBFP2, CyOFP1, TagRFP-T, mScarlet, and mStayGold sequences are highlighted with blue, orange, and green. In (a - c) truncated amino acids are highlighted with red; linkers between Nb and FP are highlighted with grey.

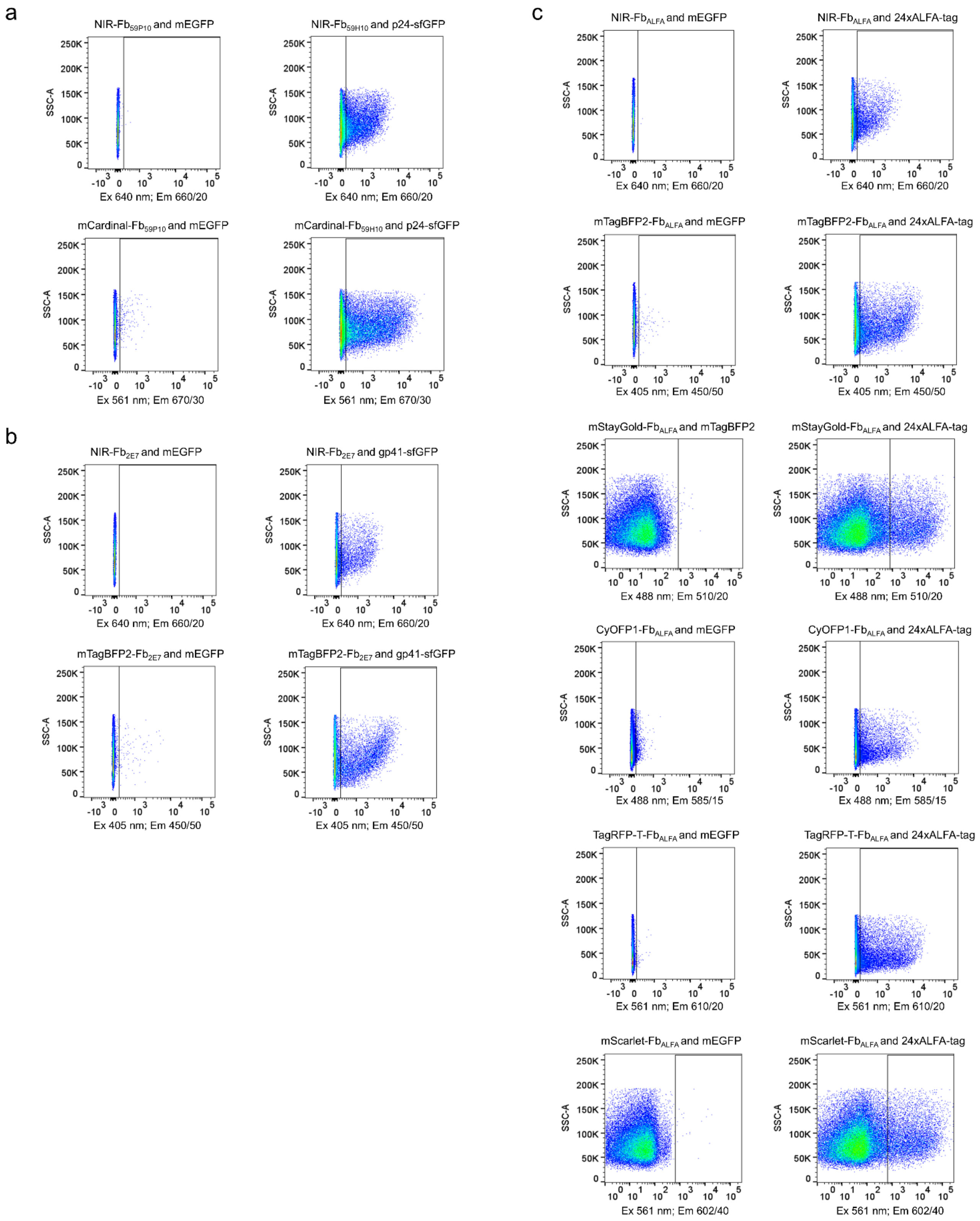

**Supplementary Figure 19. The intensity dot plots for mCardinal-Fb$_{59H10}$, mTagBFP2-Fb$_{2E7}$, and VIS-Fb$_{ALFA}$s in antigen-stabilized or -destabilized conditions in transfected HeLa cells.**

Fluorescence intensity was analyzed by flow cytometry using a 405 nm excitation laser and 450/50 nm emission filter for mTagBFP2-$Fb_{2E7}$ and mTagBFP2-$Fb_{ALFA}$; a 488 nm excitation laser and 525/50 nm emission filter for mEGFP, sfGFP and 510/20 nm emission filter for mStayGold-$Fb_{ALFA}$, 582/15 nm emission filter for CyOFP1-$Fb_{ALFA}$; a 561 nm excitation laser and 602/40 nm emission filter for mScarlet-$Fb_{ALFA}$, 610/20 nm emission filter for TagRFP-T-$Fb_{ALFA}$, 670/30 nm emission filter for mCardinal-$Fb_{59H10}$; a 640 nm excitation laser and a 660/20 nm emission filter for NIR-$Fb_{59H10}$, NIR-$Fb_{2E7}$ and NIR-$Fb_{ALFA}$. The data are presented as the intensity dot plots of corresponding VIS-Fbs (abscissa axis) versus Side Scatter Area (SSC-A) (ordinate axis) fluorescence.

jRGECO1a-Fb$_{LAG30}$

MAQVQLVESGGGLVQAGGSLRLSCAASGRTFSTSAMGWFRQAPGREREFVAAITWTVGNTIYGDSRRKWNKAGHAVRAIGRLSSPVVSERMY
PEDGALKSEIKKGLRLKDGGHYAAEVKTTYKAKKPVQLPGAYIVDIKLDIVSHNEDYTIVEQCERAEGRHSTGGMDELYKGGTGGSLVSKGE
EDNMAIIKEFMRFKVHMEGSVNGHEFEIEGEGEGRPYEAFQTAKLKVTKGGPLPFAWDILSPQFMYGSKAYIKHPADIPDYFKLSFPEGFRW
ERVMNFEDGGIIHVNQDSSLQDGVFIYKVKLRGTNFPPDGPVMQKKTMGWEATRDDLTEEQIAEFKEAFSLFDKDGDGTITTKELGTVFRSL
GQNPTEAELQDMINEVDADGDGTFDFPEFLTMMARKMNDTDSEEEIREAFRVFDKDGNGYIGAAELRHVMTDLGEKLTDEEVDEMIRVADID
GDGQVNYEEFVQMMTAKMKGRFTISRDRTKNTVDLQMDSLKPEDTAVYYCTARSRGFVLSDLRSVDSFDYKGQGTQVTVS

**Supplementary Figure 20. Amino acid sequence for jRGECO1a inserted into the LAG30 Nb for GFP.** M13 peptide in jRGECO1a sequences is highlighted in green. The sequence of mApple is highlighted in red. CaM in jRGECO1a is highlighted in blue.

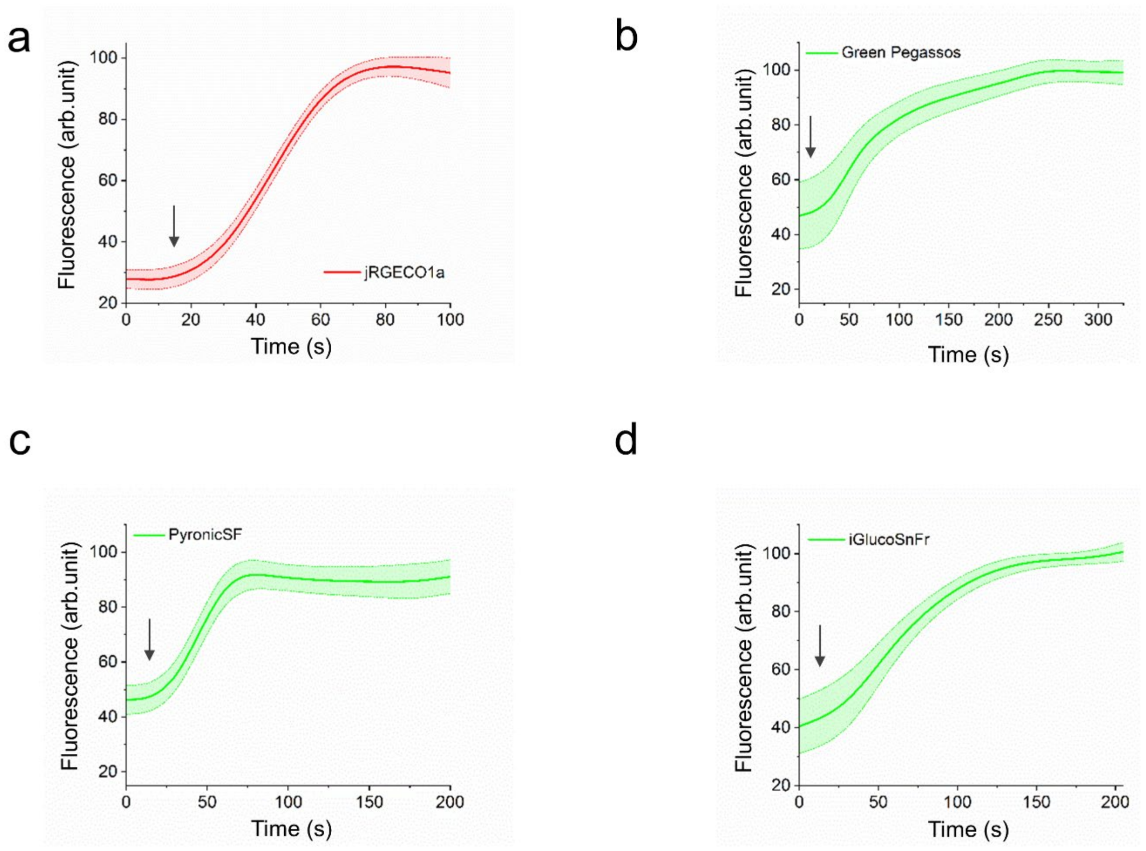

**Supplementary Figure 21. Performances of jRGECO1a in response to 5 µL ionomycin (a, *n=16*), Green Pegassos – to 1 mM pyruvate (b, *n=9*), PyronicSF – to 10 mM pyruvate (c, *n=10*), and iGlucoSnRF – to 20 mM glucose (d, *n=11*) in live HeLa cells.** The following filters were used: for imaging Green Pegassos, PyronicSF, and iGlucoSnFr 480/40 nm excitation and 535/40 nm emission; for imaging jRGECO1a 575/25 nm excitation and 615/30 nm emission. Data is presented as mean values ± s.d.

dTomato-$Fb_{LAG16}$

MAQVQLVESGGRLVQAGDSLRLSCAASGRTFSTSAMAWFRQAPGREREFVAAITWTVGNTILGDSGGSMVSKGEEVIKEFMRFKVRMEGSMNG
HEFEIEGEGEGRPYEGTQTAKLKVTKGGPLPFAWDILSPQFMYGSKAYVKHPADIPDYKKLSFPEGFKWERVMNFEDGGLVTVTQDSSLQDGT
LIYKVKMRGTNFPPDGPVMQKKTMGWEASTERLYPRDGVLKGEIHQALKLKDGGHYLVEFKTIYMAKKPVQLPGYYYVDTKLDITSHNEDYTI
VEQYERSEGRHHLFLYGMDELYKGGSVKGRFTISRDRAKNTVDLQMDNLEPEDTAVYYCSARSRGYVLSVLRSVDSYDYWGQGTQVTVS

**Supplementary Figure 22. Amino acid sequence for dTomato inserted into the LAG16 Nb for GFP.** dTomato sequence is highlighted with red, linkers are highlighted with grey.

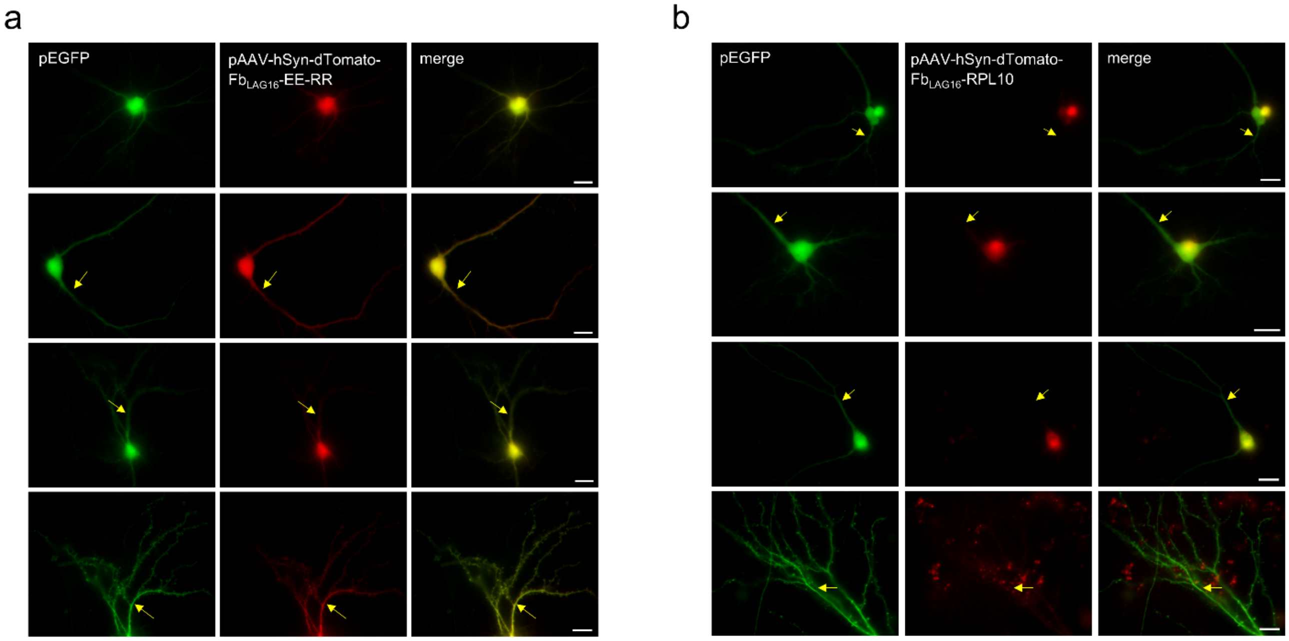

**Supplementary Figure 23. Expression of dTomato-Fb$_{LAG16}$ with different soma-localization signals. (a)** Co-expression of dTomato-Fb$_{LAG16}$ fused to EE-RR tag and mEGFP in the soma of hippocampal neuronal culture. **(b)** Co-expression of dTomato-Fb$_{LAG16}$ fused to RPL10 tag and mEGFP in the soma of hippocampal neuronal culture. The following filters were used: for imaging mEGFP 480/40 nm excitation and 535/40 nm emission; for imaging dTomato-Fb$_{LAG16}$-EE-RR and dTomato-Fb$_{LAG16}$-RPL10 575/25 nm excitation and 615/30 nm emission. Neuronal processes are marked with yellow arrows. Scale bar, 20 µm.

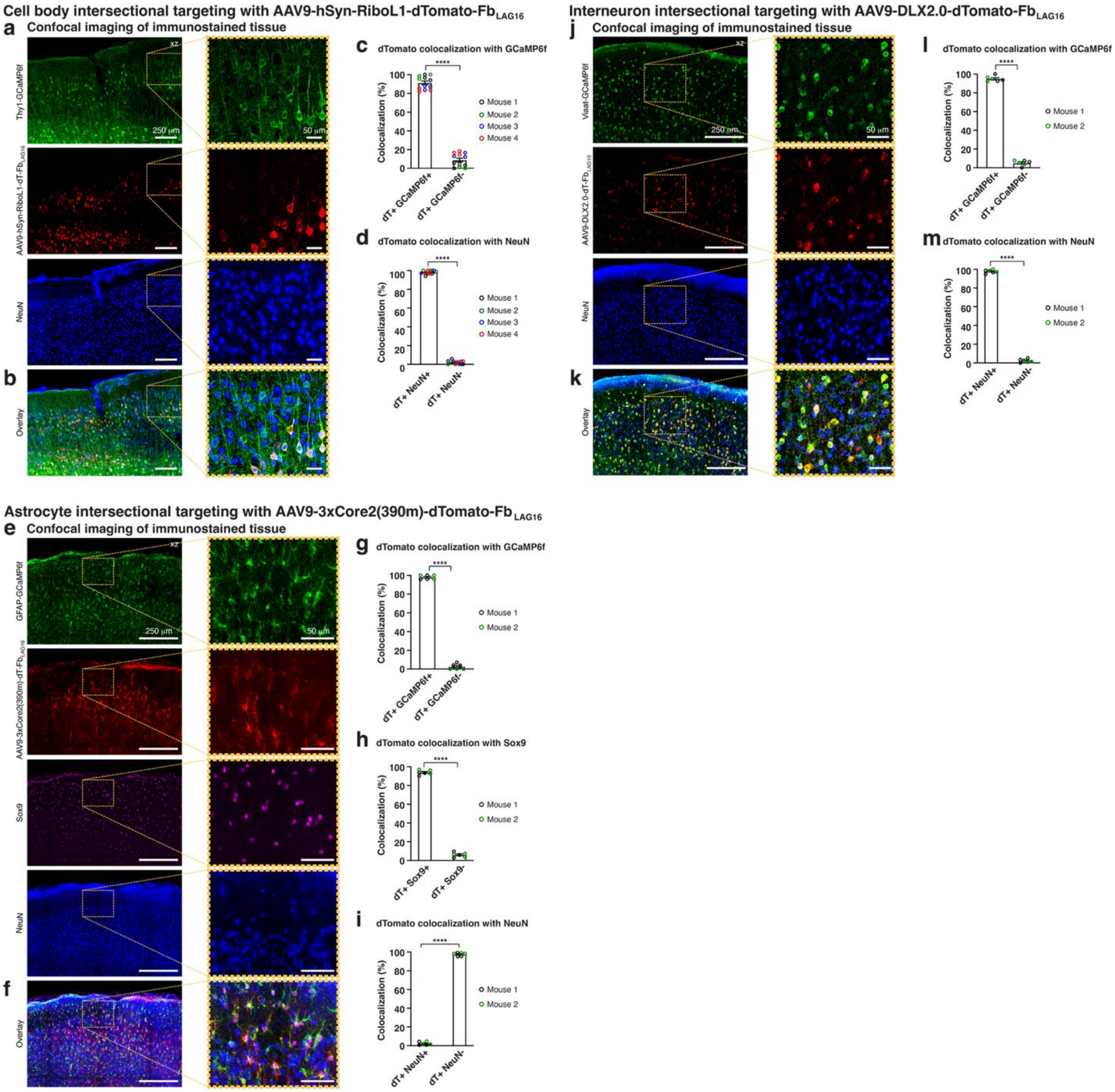

**Supplementary Figure 24. Immunostaining validation of intersectional targeting of specific cell compartments and populations with dTomato-Fb$_{LAG16}$ in GCaMP6f reporter mice. (a)** Example confocal fluorescence images showing GCaMP6f- (green) and dTomato-expressing cells (red) in a cortical tissue section from a *Thy1*-GCaMP6f mouse injected with an AAV vector driving dTomato-Fb$_{LAG16}$ expression under the control of the hSyn promoter and the soma-targeting peptide RiboL1, ~5 weeks after vector injection. The tissue was co-immunostained with NeuN (blue). Right, zoom-ins of the indicated regions. Scale bars, 250 µm (left) and 50 µm (right). **(b)** Overlay of the images in (a). **(c)** Population analysis showing the percent overlap between dTomato-Fb$_{LAG16}$ and GCaMP6f-positive cell bodies (dTomato+/GCaMP6f+: 91.2% ± 7.0%; dTomato+/GCaMP6f-: 8.8% ± 7.0%; *n=12* tissue sections from four mice). **(d)** Population analysis showing the percent overlap between dTomato-Fb$_{LAG16}$ and NeuN-positive cell bodies (dTomato+/NeuN+: 98.1% ± 1.9%; dTomato+/NeuN-: 1.9% ± 1.9%; *n=12* tissue sections from four mice). **(e)** Example confocal fluorescence images showing GCaMP6f- (green) and dTomato-expressing cells (red) in a cortical tissue section from a *GFAP*-GCaMP6f mouse injected with an AAV vector driving dTomato-$Fb_{LAG16}$ expression under the control of the astrocyte enhancer 3xCore2(390m), ~4.5 weeks after vector injection. The tissue was co-immunostained with Sox9 (purple) and NeuN (blue). Right, zoom-ins of the indicated regions. Scale bars, 250 µm (left) and 50 µm (right). **(f)** Overlay of the images in (e). **(g)** Population analysis showing the percent overlap between dTomato-$Fb_{LAG16}$ and GCaMP6f-positive cells (dTomato+/GCaMP6f+: 98.0% ± 2.2%; dTomato+/GCaMP6f-: 2.1% ± 2.2%; *n=6* tissue sections from two mice). **(h)** Population analysis showing the percent overlap between dTomato-$Fb_{LAG16}$ and Sox9-positive cells (dTomato+/Sox9+: 93.9% ± 2.4%; dTomato+/Sox9-: 6.1% ± 2.4%; *n=6* tissue sections from two mice). **(i)** Population analysis showing the percentage overlap between dTomato-$Fb_{LAG16}$ and NeuN-positive cells (dTomato+/NeuN+: 2.4% ± 1.7%; dTomato+/NeuN-: 97.6% ± 1.8%; *n=6* tissue sections from two mice). **(j)** Example confocal fluorescence images showing GCaMP6f- (green) and dTomato-expressing cells (red) in a cortical tissue section from a *Viaat*-GCaMP6f mouse injected with an AAV vector driving dTomato-$Fb_{LAG16}$ expression under the control of the DLX2.0 enhancer, ~4.5 weeks after vector injection. The tissue was co-immunostained with NeuN (blue). Right, zoom-ins of the indicated regions. Scale bars, 250 µm (left) and 50 µm (right). **(k)** Overlay of the images in (j). **(l)** Population analysis showing the percent overlap between dTomato-$Fb_{LAG16}$ and GCaMP6f-positive cell bodies (dTomato+/GCaMP6f+: 94.9% ± 2.9%; dTomato+/GCaMP6f-: 5.1% ± 2.9%; *n=6* tissue sections from two mice). **(m)** Population analysis showing the percent overlap between dTomato-$Fb_{LAG16}$ and NeuN-positive cell bodies (dTomato+/NeuN+: 97.7% ± 1.9%; dTomato+/NeuN-: 2.3% ± 1.9%; *n=6* tissue sections from two mice). Data are presented as mean values ± SD.

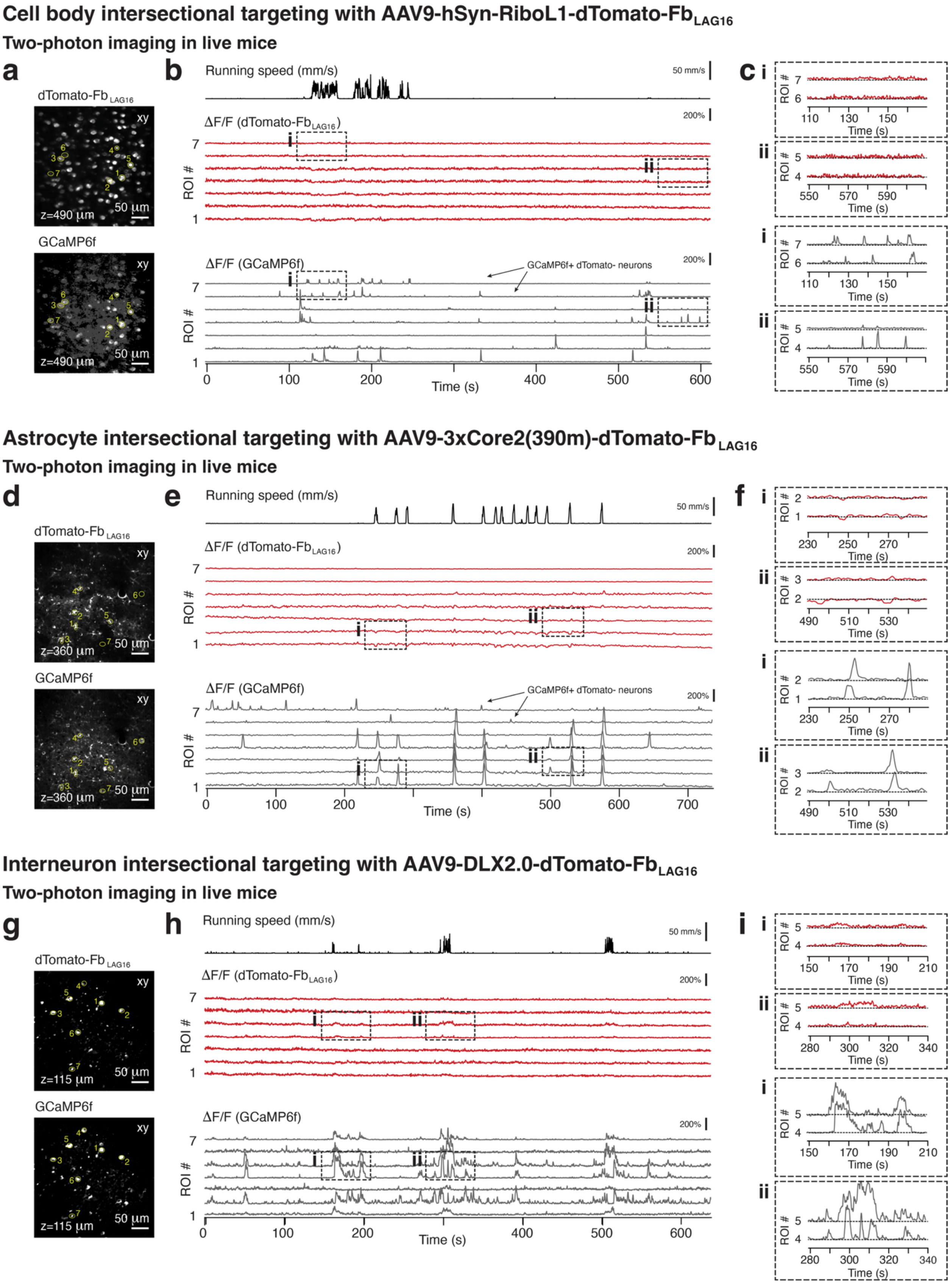

Cell body intersectional targeting with AAV9-hSyn-RiboL1-dTomato-Fb LAG16
Two-photon imaging in live mice
a
dTomato-Fb LAG16
xy
z=490 µm
50 µm
GCaMP6f
b
Running speed (mm/s)
50 mm/s
ΔF/F (dTomato-Fb LAG16)
200%
ROI #
ΔF/F (GCaMP6f)
GCaMP6f+ dTomato- neurons
Time (s)
c
Time (s)
Astrocyte intersectional targeting with AAV9-3xCore2(390m)-dTomato-Fb LAG16
Two-photon imaging in live mice
d
z=360 µm
e
f
Interneuron intersectional targeting with AAV9-DLX2.0-dTomato-Fb LAG16
Two-photon imaging in live mice
g
z=115 µm
h
i

**Supplementary Figure 25. In vivo imaging validation of intersectional targeting of specific cell compartments and populations with dTomato-Fb$_{LAG16}$ in GCaMP6f reporter mice. (a)** Example two-photon fluorescence images from a dual-color time-lapse recording showing GCaMP6f- (bottom) and dTomato-expressing cells (top) in the somatosensory cortex of a behaving Thy1-GCaMP6f mouse injected with an AAV vector driving dTomato-Fb$_{LAG16}$ expression under the control of the hSyn promoter and the soma-targeting peptide RiboL1. Imaging was performed ~5 weeks after vector injection. Recording depth (z) from the pial surface and seven somatic regions of interest (ROIs) is indicated. Scale bars, 50 µm. **(b)** Fluorescence transients in the indicated ROIs are shown as ΔF/F (red and gray) for the individual channels. The simultaneously recorded mouse's locomotor activity on a spherical treadmill is shown above the fluorescence traces. Scale bars, 50 mm/s and 200%. **(c)** Zoom-ins of the two periods indicated in (b). **(d)** Example two-photon fluorescence images from a dual-color time-lapse recording showing GCaMP6f- (bottom) and dTomato-expressing cells (top) in the somatosensory cortex of a behaving GFAP-GCaMP6f mouse injected with an AAV vector driving dTomato-Fb$_{LAG16}$ expression under the control of the astrocyte enhancer 3xCore2(390m). Imaging was performed ~4.5 weeks after vector injection. Recording depth (z) from the pial surface and seven somatic regions of interest (ROIs) is indicated. Scale bars, 50 µm. **(e)** Fluorescence transients in the indicated ROIs are shown as ΔF/F (red and gray) for the individual channels. The simultaneously recorded mouse's locomotor activity on a spherical treadmill is shown above the fluorescence traces. Scale bars, 50 mm/s and 200%. **(f)** Zoom-ins of the two periods indicated in (e). **(g)** Example two-photon fluorescence images from a dual-color time-lapse recording showing GCaMP6f- (bottom) and dTomato-expressing cells (top) in the somatosensory cortex of a behaving Viaat-GCaMP6f mouse injected with an AAV vector driving dTomato-Fb$_{LAG16}$ expression under the control of the DLX2.0 enhancer. Imaging was performed ~4.5 weeks after vector injection. Recording depth (z) from the pial surface and seven somatic regions of interest (ROIs) is indicated. Scale bars, 50 µm. **(h)** Fluorescence transients in the indicated ROIs are shown as ΔF/F (red and gray) for the individual channels. The simultaneously recorded mouse's locomotor activity on a spherical treadmill is shown above the fluorescence traces. Scale bars, 50 mm/s and 200%. **(i)** Zoom-ins of the two periods indicated in (h).

sfGFP-Fb$_{BC2}$

MQVQLVESGGGLVQPGGSLTLSCTASGFTLDHYDIGWFRQAPGKEREGVSCINNSDDDTYYADSGGSMVSKGEELFTGVVPILVELDGDVNGH
KFSVRGEGEGDATNGKLTLKFICTTGKLPVPWPTLVTTLTYGVQCFSRYPDHMKRHDFFKSAMPEGYVQERTISFKDDGTYKTRAEVKFEGDT
LVNRIELKGIDFKEDGNILGHKLEYNFNSHNVYITADKQKNGIKANFKIRHNVEDGSVQLADHYQQNTPIGDGPVLLPDNHYLSTQSKLSKDP
NEKRDHMVLLEFVTAAGITHGMDELYKGGSVKGRFTIFMNNAKDTVYLQMNSLKPEDTAIYYCAEARGCKRGRYEYDFWGQGTQVTVSS

**Supplementary Figure 26.** Amino acid sequence for sfGFP inserted into the BC2 Nb for β catenin. sfGFP sequence is highlighted with green, linkers are highlighted with grey.

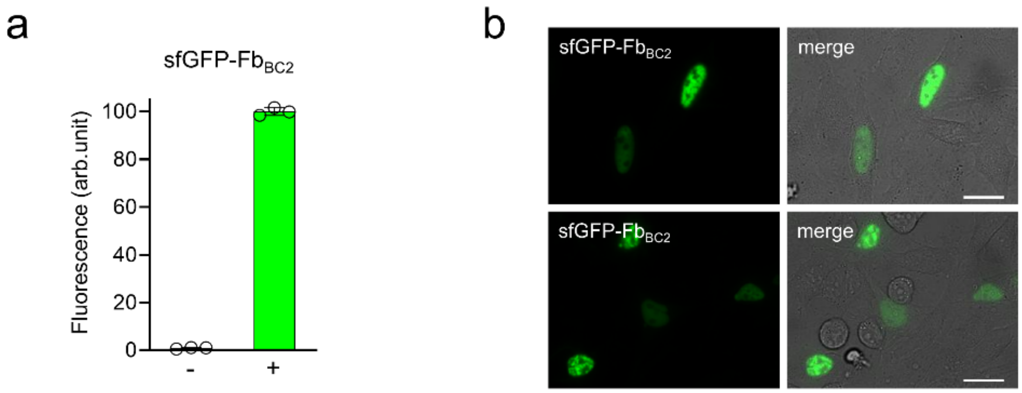

**Supplementary Figure 27. Performance of sfGFP-Fb$_{BC2}$ for β-catenin in HeLa cells.** (**a**) Fluorescence intensity of HeLa cells expressing sfGFP-Fb$_{BC2}$ for β-catenin. Co-transfection with a plasmid encoding β-catenin-sfGFP was used as a positive control (+), and co-transfection with pmTagBFP2-N1 was used as a negative control (-). (**b**) Fluorescent images of HeLa cells co-expressing sfGFP-Fb$_{BC2}$ for β-catenin and β-catenin. In (a) fluorescence intensity was analyzed by flow cytometry using a 405 nm excitation laser and 450/40 nm emission filter for mTagBFP2, a 488 nm excitation laser and 525/50 nm emission filter for sfGFP-Fb$_{BC2}$. The maximal fluorescence of antigen-bound form for sfGFP-Fb$_{BC2}$ was assumed to be 100%. Data are presented as mean values ± s.d. for $n = 3$ transfection experiments. In (b), the following filters were used: for imaging sfGFP-Fb$_{BC2}$ and mEGFP 480/40 nm excitation and 535/40 nm emission. Scale bar, 40 μm.

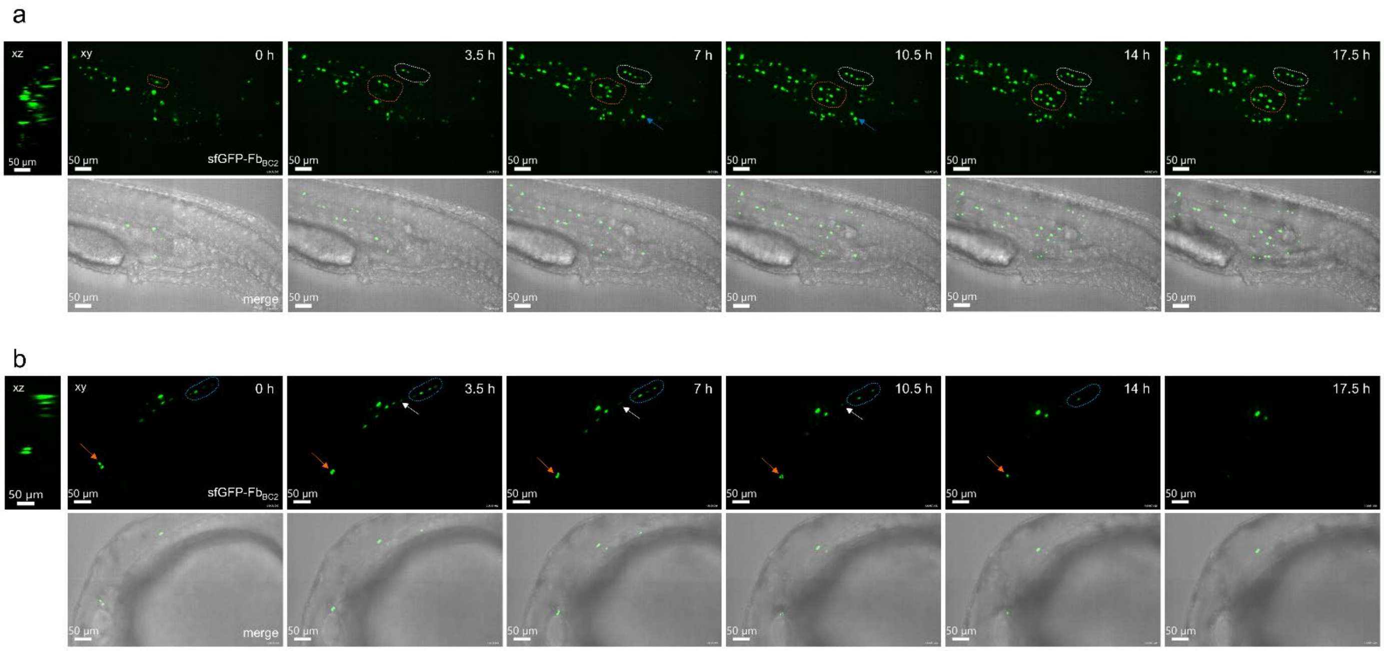

**Supplementary Figure 28. In vivo tracking of endogenous β-catenin in untreated zebrafish mosaically expressing sfGFP-Fb$_{BC2}$ in different tissues. (a, b)** 3D view time-lapsed images from intravital spinning disk fluorescent confocal microscopy of 30 hpi wild-type untreated zebrafish larvae previously injected at 1 cell-stage with pTol2-sfGFP-Fb$_{BC2}$ for β-catenin. Time-lapse images acquired every 3.5 hours are presented. Shown are xz (0 h) and xy projections for the sfGFP-Fb$_{BC2}$ channel, as well as xy projection of the merged brightfield and green fluorescence channels. (a) In vivo tracking of endogenous β-catenin in zebrafish trunk. Dashed white and orange ROIs indicate cells displaying an increase in endogenous β-catenin signal over time. Blue arrow indicates cells displaying a decrease in endogenous β-catenin signal over time. (b) In vivo tracking of endogenous β-catenin in zebrafish head. Dashed blue ROI and orange arrows indicate cells displaying a decrease in endogenous β-catenin signal over time; white arrow indicates cells displaying an increase in endogenous β-catenin signal over time. Scalebar, (a, b) 50 µm.

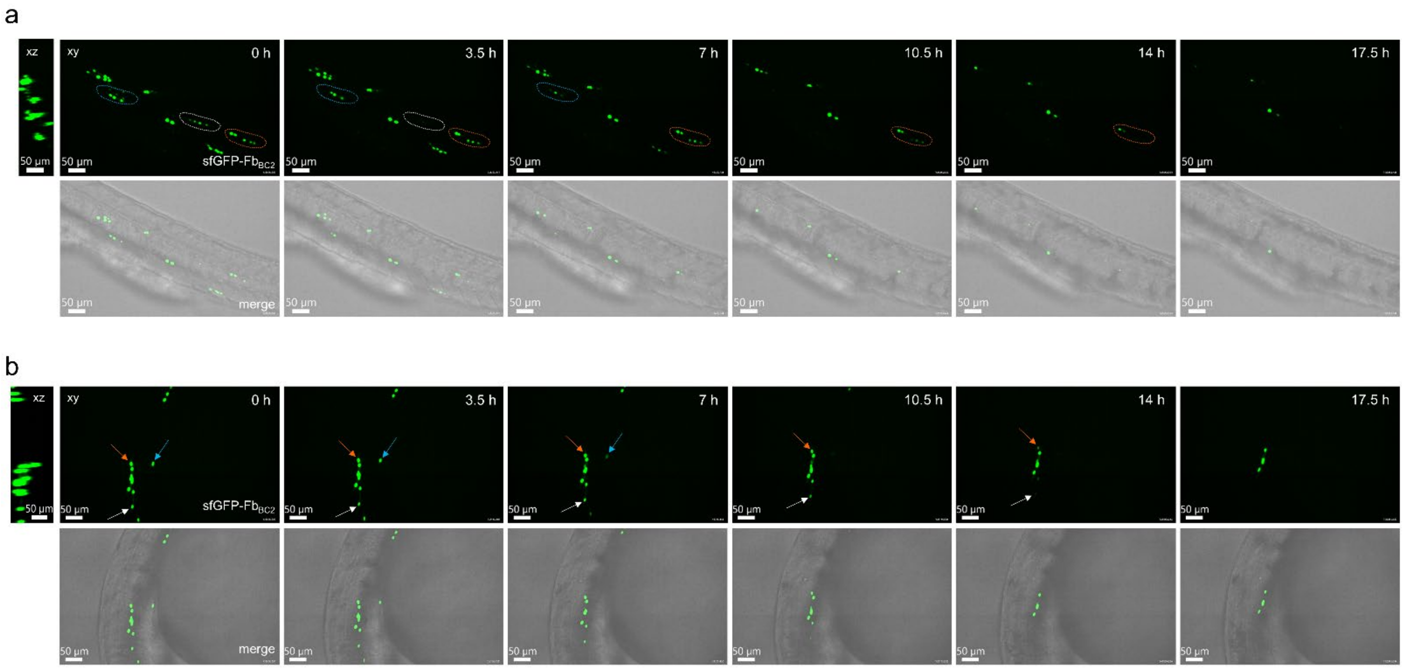

**Supplementary Figure 29. In vivo tracking of endogenous β-catenin in zebrafish mosaically expressing sfGFP-Fb$_{BC2}$ and treated with 10 µM IWR-1 in different tissues. (a, b)** 3D view time-lapsed images from intravital spinning disk fluorescent confocal microscopy of 30 hpi wild-type untreated zebrafish larvae previously injected at 1 cell-stage with pTol2-sfGFP-Fb$_{BC2}$ for β-catenin. Time-lapse images acquired every 3.5 hours are presented. Shown are xz (0 h) and xy projections for the sfGFP-Fb$_{BC2}$ channel, as well as xy projection of the merged brightfield and green fluorescence channels. (a) In vivo tracking of endogenous β-catenin in zebrafish trunk. Dashed white, orange and blue ROIs indicate cells displaying a decrease in endogenous β-catenin signal over time. (b) In vivo tracking of endogenous β-catenin in zebrafish head. White, orange and blue arrows indicate cells displaying a decrease in endogenous β-catenin signal over time. Scalebar, (a, b) 50 µm.

**Supplementary Table 1.** Sequences of oligonucleotides used in this study.

| Oligonucleotide | Sequence (5'-3') |
|---|---|
| mCherry-dMVSKGEEDN-fw | gatcgtagctcctatgaagactccatggccatcatcaaggagttcatg |
| mCherry-GBP1-rv | catggacgagctgtacaaggtgaagggccgtttcaccatc |
| GBP1-mTagBFP2-fw | ctcctatgaagactccatggccctgattaaggagaacatg |
| GBP1-mTagBFP2-rv | catgttctccttaatcagggccatggagtcttcataggag |
| mTagBFP2-GBP1-fw | ctagcaaactggggcacaaggtgaagggccgtttcac |
| mTagBFP2-GBP1-rv | gtgaaacggcccttcaccttgtgccccagtttgctag |
| GBP1-mTFP1-fw | gtagctcctatgaagactccatgggcgtaatcaagcccgac |
| GBP1-mTFP1-rv | gtcgggcttgattacgcccatggagtcttcataggagctac |
| mCerulean3-GBP1-fw | ggacgagctgtacaaggtgaagggccgtttcacc |
| mCerulean3-GBP1-rv | ggtgaaacggcccttcaccttgtacagctcgtcc |
| GBP1-NG-fw | gtagctcctatgaagactccatggctagtctgcctg |
| GBP1-NG-rv | caggcagactagccatggagtcttcataggagctac |
| NG-GBP1-fw | ctttacggacgtgatgggcatggacgagctgtacaaggtgaagggccgtttcac |
| NG-GBP1-rv | gtgaaacggcccttcaccttgtacagctcgtccatgcccatcacgtccgtaaag |
| GBP1-mOrange-fw | gtagctcctatgaagactccatggccatcatcaaggag |
| GBP1-mOrange-rv | ctccttgatgatggccatggagtcttcataggagctac |
| GBP1-mScarlet-fw | ctcctatgaagactccatggccgtgatcaaggagttcatg |
| GBP1-mScarlet-rv | catgaactccttgatcacggccatggagtcttcataggag |
| mNeptune2-GBP1-fw | ctagcaaactggggcacaaagtgaagggccgtttcac |
| mNeptune2-GBP1-rv | gtgaaacggcccttcactttgtgccccagtttgctag |
| GBP1-mCardinal-fw | ctcctatgaagactccatggccctgatcaaggagaacatg |
| GBP1-mCardinal-rv | catgttctccttgatcagggccatggagtcttcataggag |
| GBP1-PAmCherry-fw | gtagctcctatgaagactccatggccatcattaaggag |
| GBP1-PAmCherry-rv | ctccttaatgatggccatggagtcttcataggagctac |
| GBP1-mEos4a-fw | ctcctatgaagactccatggccgcgattaagccagacatg |
| GBP1-mEos4a-rv | catgtctggcttaatcgcggccatggagtcttcataggag |
| mEos4a-GBP1-fw | gattgcctgacaatgccagatacaaggtgaagggccgtttcac |
| mEos4a-GBP1-rv | gtgaaacggcccttcaccttgtatctggcattgtcaggcaatc |
| mTag-Fb-BamHI-fw | catggatccgcctccggcgaccaagtccaactggtg |
| mTag-Fb-XhoI-rv | ctgtctcgagctagctggagacggtgacctg |
| CyOFP-Fb-BamHI-fw | ctaggatccaccggtcgccaccatggaccaagtccaactg |
| CyOFP-Fb-XbaI-rv | cttctagattagctggagacggtgacctgggtg |
| mito-mScI-Fb-fw | gatccaccggtcgccaccatggaccaagtccaac |
| mito-mScI-Fb-rv | gttggacttggtccatggtggcgaccggtggatc |
| mScI-Fb-DNA-fw | gtcaccgtctccagctaaagcggccgcgactctag |
| mScI-Fb-DNA-rv | ctagagtcgcggccgctttagctggagacggtgac |
| mEGFP-Y66G-fw | cgtgaccaccctgaccggcggcgtgcagtgcttc |
| mEGFP-Y66G-rv | gaagcactgcacgccgccggtcagggtggtcacg |
| LAM2-mTagBFP-fw | cacggattacgcggatagcatggccctgattaaggagaacatg |
| LAM2-mTagBFP-rv | catgttctccttaatcagggccatgctatccgcgtaatccgtg |

| | |
|---|---|
| mTagBFP-LAM2-fw | caaactggggcacaaggtgaaaggtcggtttac |
| mTagBFP-LAM2-rv | gtaaaccgacctttcaccttgtgccccagtttg |
| LAM2-Wasabi-fw | cacggattacgcggatagcatgggcgtaatcaagcccgac |
| LAM2-Wasabi-rv | gtcgggcttgattacgcccatgctatccgcgtaatccgtg |
| Wasabi-LAM2-fw | catggacgagctgtacaaggtgaaaggtcggtttac |
| Wasabi-LAM2-rv | gtaaaccgacctttcaccttgtacagctcgtccatg |
| LAM2-Clover-fw | cacggattacgcggatagcatggccctgttcaccggggtggtg |
| LAM2-Clover-rv | caccaccccggtgaacagggccatgctatccgcgtaatccgtg |
| Nbactin-Scarlet-L(3)-fw | caaactatggagacttcgggggtagcatggtgagcaagggcgag |
| Nbactin-Scarlet-L(3)-rv | ctcgcccttgctcaccatgctacccccgaagtctccatagtttg |
| Card-Nbact-fw1 | caaactggggcacaaaggggggtagcgtgaagggccgattc |
| Card-Nbact-rv1 | gaatcggcccttcacgctacccccctttgtgccccagtttg |
| CMVd1-Nbactin-fw | cgtcagatccgctagcatggctcaggtgcagctg |
| CMVd1-Nbactin-rv | cagctgcacctgagccatgctagcggatctgacg |
| Nbactin-CMVd1-fw | caggtcaccgtctcctcataacccgggatccac |
| Nbactin-CMVd1-rv | gtggatcccgggttatgaggagacggtgacctg |
| 59H10-Cardinal-fw | cagtgctcgccgatagtgggggtagcatggtgagcaag |
| 59H10-Cardinal-rv | cttgctcaccatgctacccccactatcggcgagcactg |
| Cardinal-59H10-fw | caaactggggcacaaaggggggtagcgtgaagggcag |
| Cardinal-59H10-rv | ctgcccttcacgctacccccctttgtgccccagtttg |
| gp41-BFP2-fw | caactacgccgatagcgggggtagcatggtgagcaagggcgaggagctgattaaggagaac |
| gp41-BFP2-rv | gttctccttaatcagctcctcgcccttgctcaccatgctacccccgctatcggcgtagttg |
| BFP2-gp41-fw | caaactggggcacaagggggggtagcgttaagggtagattc |
| BFP2-gp41-rv | gaatctacccttaacgctacccccccttgtgccccagtttg |
| ALFA-BFP2-fw | catgtatcgcgaaagcgggggtagcatggtgagcaagggcgaggagctgattaaggagaac |
| ALFA-BFP2-rv | gttctccttaatcagctcctcgcccttgctcaccatgctacccccgctttcgcgatacatg |
| BFP2-ALFA-fw | caaactggggcacaagggggggtagcgttcaaggccgctttac |
| BFP2-ALFA-rv | gtaaagcggccttgaacgctacccccccttgtgccccagtttg |
| ALFA-CyOFP-fw | gaaagcgggggtagcatggtgagcaagggcgag |
| ALFA-CyOFP-rv | ctcgcccttgctcaccatgctacccccgctttc |
| CyOFP-ALFA-fw | ctccaacctgggcatggacgagctgtacaagggggggtagcgttcaaggccgctttac |
| CyOFP-ALFA-rv | gtaaagcggccttgaacgctacccccccttgtacagctcgtccatgcccaggttggag |
| TagRFP-ALFA-fw | caaactggggcacaaaggggggtagcgttcaag |
| TagRFP-ALFA-rv | cttgaacgctacccccctttgtgccccagtttg |
| ALFA-mSG-fw | catgtatcgcgaaagcgggggtagcatggtgtctac |
| ALFA-mSG-rv | gtagacaccatgctacccccgctttcgcgatacatg |
| mSG-ALFA-fw | gaggcccacctgggggggtagcgttcaaggccgctttac |
| mSG-ALFA-rv | gtaaagcggccttgaacgctacccccccaggtgggcctc |
| NES-gp41-fw | gatgaggtggatggagccatgtctggtatagtgcag |
| NES-gp41-rv | ctgcactataccagacatggctccatccacctcatc |
| sfGFP-pcDNA-fw | gacgagctgtacaagtaactcgagtctagagggcccg |
| sfGFP-pcDNA-rv | cgggccctctagactcgagttacttgtacagctcgtc |
| H2B-ALFA-fw | gatccaccggtcgccaccatgcatctagatacctc |
| H2B-ALFA-rv | gaggtatctagatgcatggtggcgaccggtggatc |
| mGFP-pcDNA-fw | gacgagctgtacaagtaatctagagggccctattc |

| | |
|---|---|
| mGFP-pcDNA-rv | gaatagggccctctagattacttgtacagctcgtc |
| DNA-p24-fw | gaataaggagcgatcgccatgagccctaccagcattc |
| DNA-p24-rv | gaatgctggtagggctcatggcgatcgctccttattc |
| GFP-CAAX-fw | catggacgagctgtacaaggctagcggtaaaaagaag |
| GFP-CAAX-rv | cttctttttaccgctagccttgtacagctcgtccatg |
| plasmid-IgK-fw | ctagttaagcttggtaccatggagacagacacactc |
| plasmid-IgK-rv | gagtgtgtctgtctccatggtaccaagcttaactag |
| IgK-GBP1-fw | caggttccactggtgacatggaccaagtccaactg |
| IgK-GBP1-rv | cagttggacttggtccatgtcaccagtggaacctg |
| LAG30-M13-fw | catctatggcgacagccgtcgtaagtggaataag |
| LAG30-M13-rv | cttattccacttacgacggctgtcgccatagatg |
| CaM-LAG30-fw | caaatgatgacagcgaagatgaaaggtcgcttc |
| CaM-LAG30-rv | gaagcgacctttcatcttcgctgtcatcatttg |
| LAG16-dTomato-fw | cacgattctgggcgattcgatggtgagcaagggcgaggaggtcatcaaagag |
| LAG16-dTomato-rv | ctctttgatgacctcctcgcccttgctcaccatcgaatcgcccagaatcgtg |
| dTomato-LAG16-fw | catggacgagctgtacaaggtgaaagggcgtttcac |
| dTomato-LAG16-rv | gtgaaacgccctttcaccttgtacagctcgtccatg |
| LAG16-dTomato-fw2 | cacgattctgggcgattcgggggggtagcatggtgagcaagggcgag |
| LAG16-dTomato-rv2 | ctcgcccttgctcaccatgctacccccccgaatcgcccagaatcgtg |
| dTomato-LAG16-fw2 | catggacgagctgtacaagggggggtagcgtgaaagggcgtttcac |
| dTomato-LAG16-rv2 | gtgaaacgccctttcacgctacccccccttgtacagctcgtccatg |
| LAG16-AgeI-fw | caaccggtcgccaccatggcgcaagtccagctg |
| LAG16-AscI-rv | ctggcgcgccgtcgacttaagaaacggtcacctgagtg |
| EERR-LAG16-fw | cagtcggatccgccaccatggcgcaagtccagctg |
| EERR-LAG16-rv | cagctggacttgcgccatggtggcggatccgactg |
| LAG16-EERR-fw | cactcaggtgaccgtttctgagctcgggggcagcggtg |
| LAG16-EERR-rv | caccgctgcccccgagctcagaaacggtcacctgagtg |
| RiboL-LAG16-fw | gaggatccacccgccaccatggcgcaagtccagctg |
| RiboL-LAG16-rv | cagctggacttgcgccatggtggcgggtggatcctc |
| LAG16-RiboL-fw | cactcaggtgaccgtttctgggggcagcggtgggtc |
| LAG16-RiboL-rv | gacccaccgctgcccccagaaacggtcacctgagtg |
| BamHI-LAG16-fw | gaggatccggcgccaccatggcgcaagtccagctg |
| BamHI-LAG16-rv | cagctggacttgcgccatggtggcgccggatcctc |
| LAG16-RPL10-fw | cactcaggtgaccgtttcttcaggccggactcagatc |
| LAG16-RPL10-rv | gatctgagtccggcctgaagaaacggtcacctgagtg |
| BC2-sfGFP-fw | tacatattatgccgactctgggggtagcatggtgagcaagggcgag |
| BC2-sfGFP-rv | ctcgcccttgctcaccatgctacccccagagtcggcataatatgta |
| sfGFP-BC2-fw | catggacgagctgtacaagggggggtagcgtcaaaggaagattcac |
| sfGFP-BC2-rv | gtgaatcttcctttgacgctacccccccttgtacagctcgtccatg |
| Tol2-BC2-fw | gaattcctcgacggatccaccatgcaggttcagcttgtag |
| Tol2-BC2-rv | ctacaagctgaacctgcatggtggatccgtcgaggaattc |
| BC2-Tol2-fw | caggtgacagtgagtagttagggatcccccgggtcgcg |
| BC2-Tol2-rv | cgcgacccgggggatccctaactactcactgtcacctg |
